# SORT disentangles multilevel gene programs across heterogeneous spatial transcriptomic atlases

**DOI:** 10.64898/2026.10.04.756488

**Authors:** Lei Zhang, Limin Li, Shuqin Zhang

## Abstract

Understanding how transcriptional programs are organized across biological contexts is essential for deciphering tissue development, disease progression, and species-specific adaptations. Spatial transcriptomic atlases spanning patients, developmental stages, and species provide an opportunity to resolve gene expression programs (GEPs) that are sample-specific, group-shared, or region-associated, yet strong sample-specific variation can obscure weaker shared programs. Here we introduce SORT, a matrix factorization framework that jointly recovers multilevel GEPs and their spatial activities, using orthogonality to reduce program redundancy and graph total variation to preserve spatial boundaries. In simulations with strong patient variation, SORT more accurately recovered patient-specific, subtype-shared, and region-associated programs than existing methods. Across 108 pancreatic ductal adenocarcinoma sections, SORT resolved tumor and stromal programs with distinct and overlapping activities and linked these activities to survival across four external cohorts. In a cross-species cerebellar atlas, SORT distinguished conserved layer-associated programs from species-specific activity variation. Across eight stages of mouse organogenesis, SORT revealed GEPs with anatomical redeployment and graded spatial activity, as well as stage-dependent local co-deployment of distinct programs.

## Introduction

Tissue organization and function are shaped not only by the expression of individual genes, but more importantly by the coordinated spatial deployment of transcriptional programs. Spatial transcriptomic studies have uncovered coordinated gene expression programs (GEPs) associated with metabolic zonation in liver, injury-associated microenvironments and localized cellular interactions in kidney, and spatiotemporal organization in the developing limb [1–3]. Beyond individual genes, GEPs provide a higher-order view of tissue state, helping reveal which programs recur across contexts and where they are spatially deployed or redeployed. With rapid advances in spatial transcriptomics, increasingly comprehensive multi-section atlases now span patients, disease states, developmental stages and species [4–6]. This expansion enables systematic investigation of GEPs across heterogeneous sections and biological contexts, providing opportunities to uncover transcriptional programs shared across sections and biological groups as well as those specialized for particular tissues, disease states or developmental processes.

Recovering GEPs from heterogeneous spatial atlases is challenging because gene programs can exhibit different patterns and levels of sharing: some may be confined to individual sections, others may recur within biological groups, and still others may mark anatomical regions observed in multiple sections. At the same time, program activities can vary continuously in magnitude and spatial extent, and multiple programs may overlap within the same tissue. These sources of variation can confound the distinction between broadly shared programs and section-specific signals, particularly when strong sample-specific variation obscures weaker programs shared across biological groups or anatomical regions. In addition, correlated gene programs can lead to redundant representations, making it difficult to recover a compact and interpretable set of programs. Atlas-scale analysis therefore requires a common gene-program profile space that resolves nonredundant programs at different levels of sharing, together with section-specific activity maps that capture local spatial coherence while preserving boundaries between adjacent tissue structures.

Existing methods capture different aspects of transcriptional variation and spatial organization. Traditional matrix decomposition methods such as principal component analysis (PCA) [7], nonnegative matrix factorization (NMF) [8] and semi-nonnegative matrix factorization (semi-NMF) [9] provide low-dimensional expression representations without explicitly modeling spatial relationships among measurements. GraphPCA extends PCA with spatial graph structure to obtain spatially informed components [10], whereas spatial factorization methods such as nonnegative spatial factorization (NSF) [11] and neighborhood nonnegative matrix factorization (NNMF) [12] incorporate spatial dependence into matrix factorization to recover spatially coherent or overlapping programs. These methods are primarily designed for individual sections. When extended to multisample settings, methods differ in how they share information across sections. Multi-sample non-negative spatial factorization (mNSF) shares nonnegative gene loadings across samples while modeling sample-specific spatial factors with separate Gaussian processes [13]. STAMP learns spatial topics across samples and can model sample-associated variation as well as gradual changes in gene modules along continuous biological processes such as development [14]. Although these approaches provide increasingly flexible representations of shared and adaptive transcriptional structure, they do not explicitly recover multilevel GEPs across sections while maintaining spatially coherent, boundary-aware activity maps and avoiding redundant program representations.

To address these challenges, we developed SORT, a matrix factorization model that jointly learns shared transcriptomic profiles and section-specific activity maps with spatial regularization and orthogonality constraints. Orthogonality promotes distinct gene-program profiles and reduces redundant representations, while graph total variation regularization encourages spatially coherent activities and preserves anatomical boundaries. This formulation enables joint recovery of GEPs at different levels of shared specificity and comparison of their spatial deployment across heterogeneous sections. We evaluated SORT in simulations with known ground truth, demonstrating its ability to recover patient-specific, disease subtype-shared and region-associated programs under strong sample variation and assessing the contributions of orthogonality and spatial regularization. We then applied SORT to pancreatic ductal adenocarcinoma, cross-species cerebellar, and mouse organogenesis atlases [4–6] to reveal context-dependent program deployment across patients, species and developmental stages. These analyses linked program activities to patient survival, distinguished conserved tissue organization from species-specific spatial variation, and revealed developmental changes in spatial deployment.

## Results

### Overview of SORT

In heterogeneous spatial transcriptomic atlases, GEPs can recur at multiple levels of shared specificity. A given GEP may be restricted to a single sample, shared within a biological group, or spatially associated with a distinct tissue region. SORT is designed to recover these multilevel GEPs while capturing the heterogeneous structure of their spatial activities. It jointly factorizes the expression matrices of all sections into shared transcriptomic profiles and corresponding section-specific activity maps, with each profile and its activity maps together defining a GEP. This unified decomposition enables biological interpretation of the recovered programs, facilitates comparison of their activities across samples and conditions, and provides a common framework for downstream analyses linking GEP activity to clinical outcomes or developmental time (Fig. 4).

**Figure 1.**
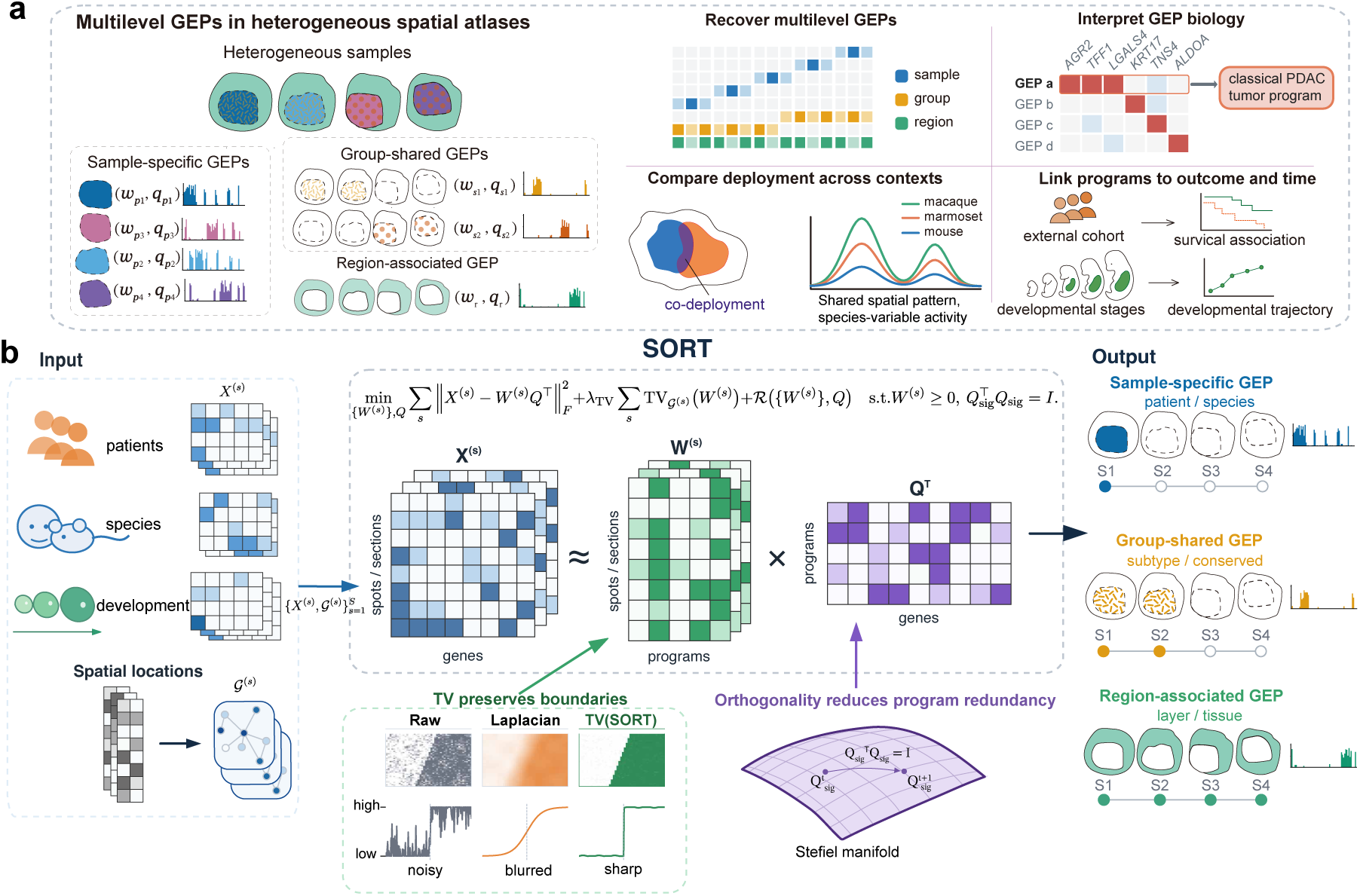
Gene expression programs (GEPs) across levels of shared specificity and the SORT framework. **(a)** GEPs can be restricted to a single sample, shared within a biological group or associated with a tissue region in heterogeneous spatial atlases. Each GEP is defined by a transcriptomic profile and corresponding spatial activity maps. Downstream analyses include accurate recovery of GEPs across levels of shared specificity, gene-based biological interpretation, comparison of spatial deployment across contexts, and integration with survival or developmental time. **(b)** SORT jointly factorizes multi-section spatial transcriptomic data into shared transcriptomic profiles (*Q*) and section-specific activity matrices (*W* ^(*s*)^), with regularization penalties and orthogonality constraints to promote spatial coherence and reduce redundancy among signal programs, enabling joint recovery of multilevel GEPs and comparison of their spatial deployment across sections.

**Figure 2.**
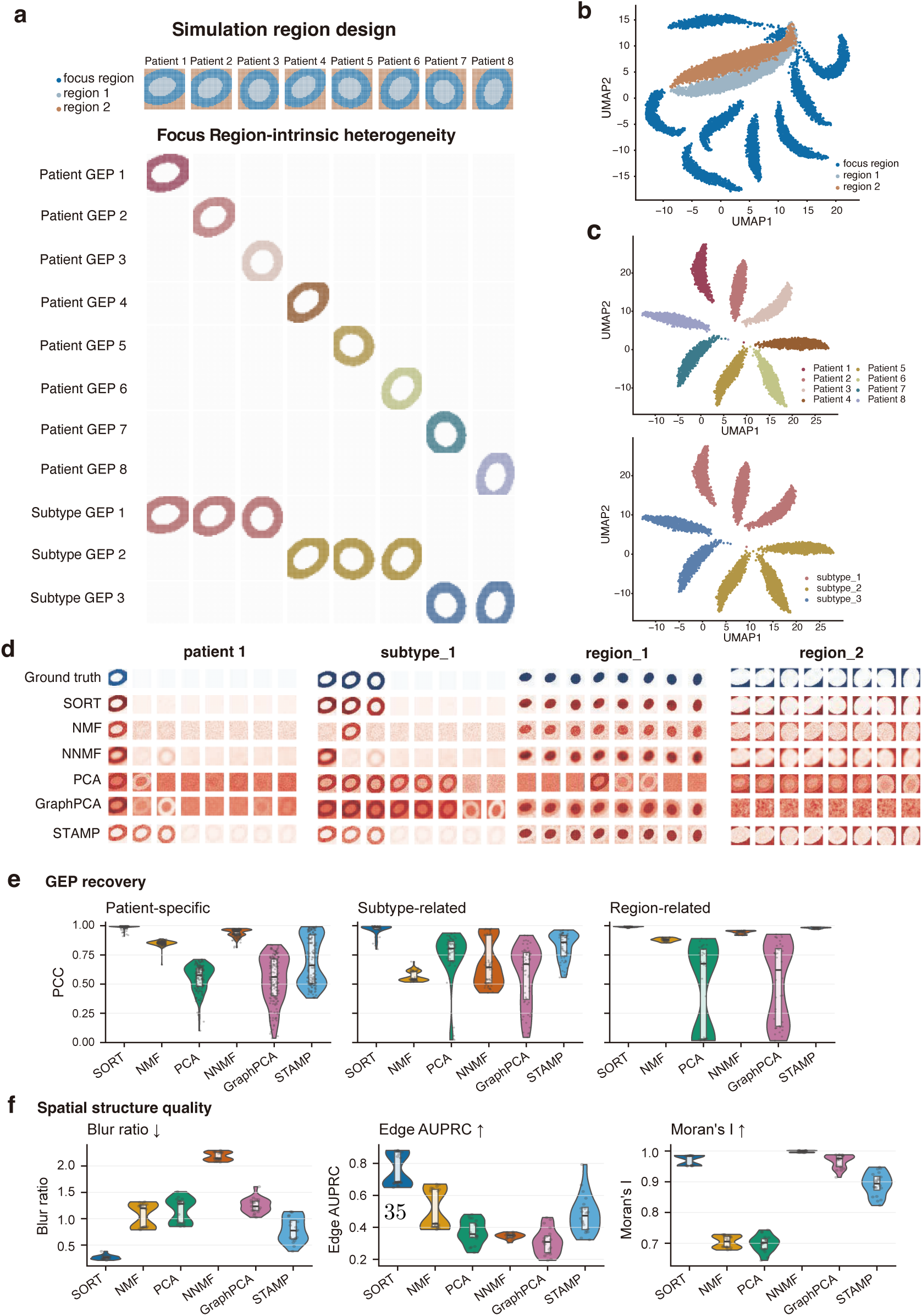
Benchmarking SORT on simulated multipatient spatial transcriptomics data. **(a)** Simulation design. Each simulated dataset comprised eight patient sections, three spatial regions per section, and 13 ground-truth programs: eight patient-specific, three subtype-related, and two region-related programs. The focus region contained the patient-specific and subtype-related programs, while the region-related programs were localized outside the focus. **(b)** Uniform manifold approximation and projection (UMAP) of all spatial locations from a representative replicate, colored by region. **(c)** UMAP of locations within the focus region from the same replicate, colored by patient (top) or subtype (bottom). **(d)** Ground-truth and recovered activities for one patient-specific, one subtype-related, and two region-related programs in the representative replicate. **(e)** Quantitative recovery across replicates. Signed Pearson correlation between each ground-truth activity vector and its best-matching inferred counterpart across 20 independent simulations. Each point represents one ground-truth program in one replicate (patient-specific: *n* = 160; subtype-related: *n* = 60; region-related: *n* = 40). **(f)** Spatial fidelity measured by Blur ratio (lower is better), edge area under the precision–recall curve (AUPRC; higher is better) and Moran’s *I*. Each point represents one simulation replicate after aggregation across matched programs (*n* = 20). Violin plots in **(e,f)** show the full distributions, with medians and interquartile ranges indicated.

**Figure 3.**
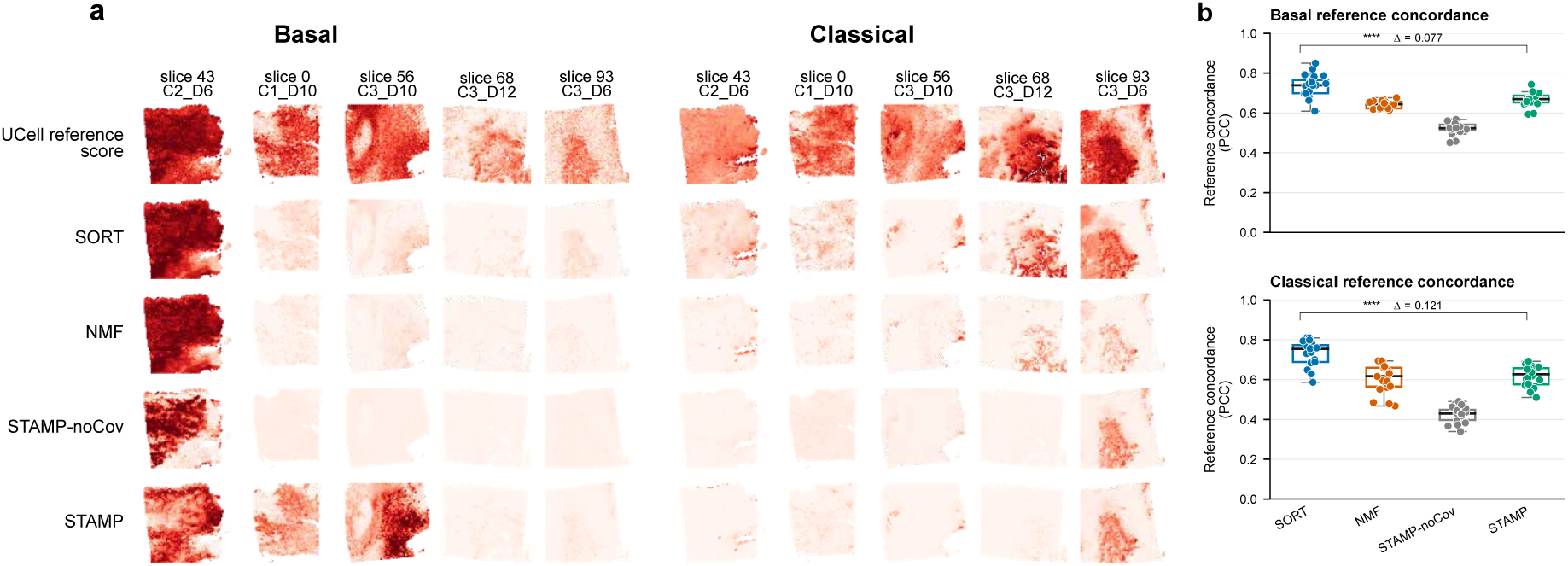
Benchmarking SORT on basal-like and classical PDAC reference signals. **(a)** Spatial maps for five of the ten sections in one representative stratified subset. Rows display basal-like or classical UCell reference scores computed from published signatures and the recovered activity of the best-matching component, for SORT, NMF, STAMP without patient covariates (STAMP-noCov), and STAMP with patient identity as a covariate. **(b)** Concordance between the best-matching component activities and the reference scores across 20 stratified subsets for basal-like and classical subtypes. Each point represents one subset. ‘****’ indicates *P <* 10*^−^*^4^ from paired two-sided Wilcoxon signed-rank tests. ‘Δ’ is the median within-subset difference between SORT and STAMP across the 20 subsets.

**Figure 4.**
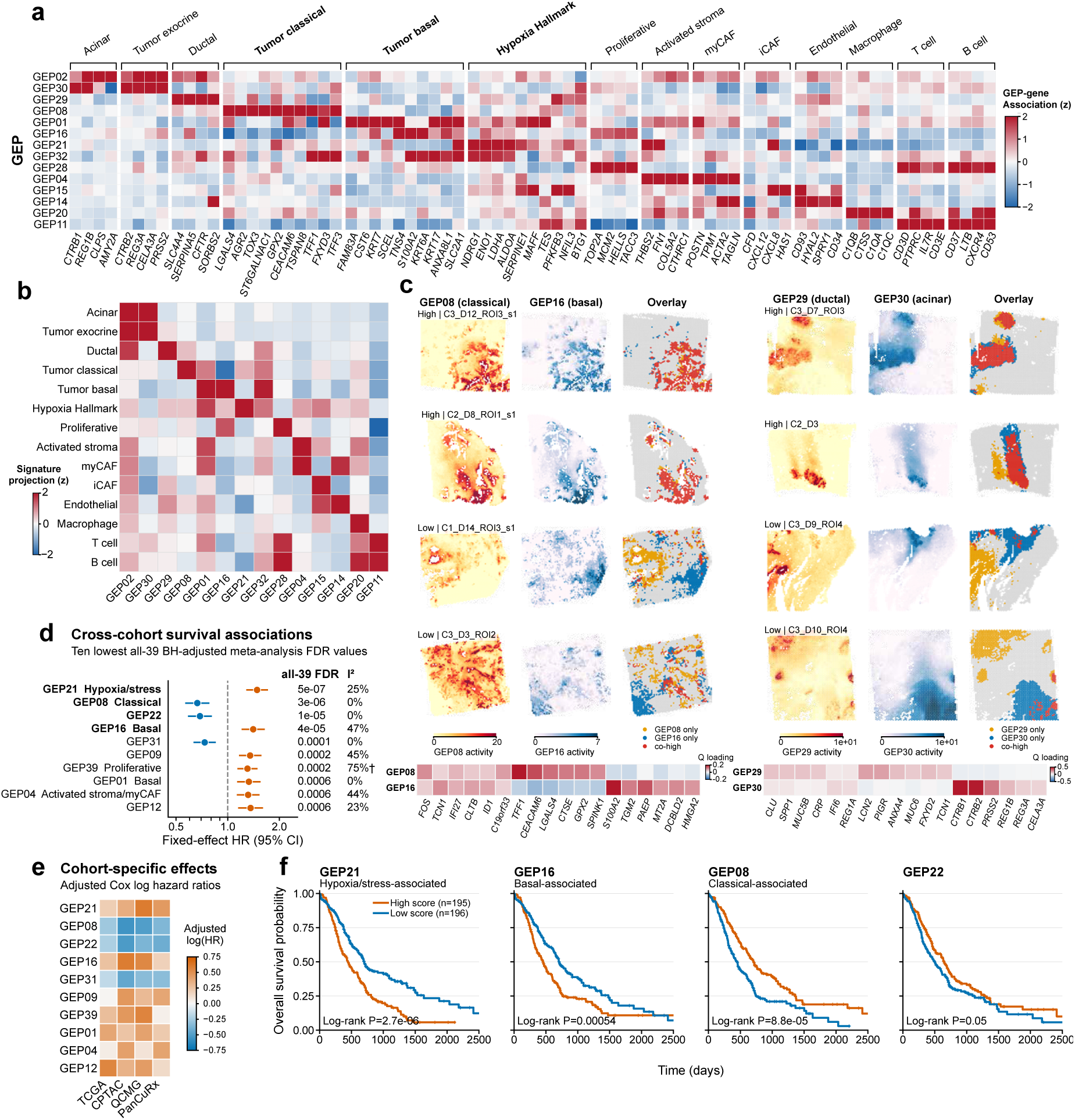
PDAC programs show variable spatial deployment and survival associations. **(a)** Associations of 14 selected GEP activities with 74 representative genes drawn from published PDAC and tumor microenvironment signatures. Genes are grouped by source. Colors show the ordinary least-squares coefficient *B_gk_*, quantifying the association between each gene and GEP activity after controlling for other programs. Coefficients were standardized within each gene across all 39 signal GEPs and clipped to [−2, 2]. **(b)** Projection of published PDAC and tumor microenvironment signatures onto the GEPs shown in **(a)**. Colors show the signed mean of *B_gk_* across genes in the signature, standardized across the displayed GEPs. **(c)** Sections with relatively high or low colocalization between GEP08 and GEP16, and between GEP29 and GEP30. High activity was defined using section-specific 80th-percentile thresholds. Colors indicate locations high only for the first GEP (orange), high only for the second GEP (blue), high for both (red), or high for neither (gray). Lower heatmaps show *Q* profile values for selected genes. **(d)** Survival associations for the ten GEPs with the smallest Benjamini–Hochberg FDRs among all 39 signal GEPs. Points and bars show hazard ratios and 95% confidence intervals from fixed-effect inverse-variance meta-analysis. Blue and orange indicate protective and adverse associations, respectively. *I*^2^ estimates the proportion of variation among cohort effects attributable to heterogeneity. Daggers mark *I*^2^ ≥ 60%. **(e)** Covariate-adjusted log hazard ratios for the ten GEPs across four cohorts. **(f)** Pooled Kaplan–Meier curves for the four GEPs with the smallest FDRs across all 39 signal GEPs. Cohort-standardized scores were split at the pooled median. Censor marks are shown, and log-rank *P* values are reported descriptively. Supplementary Figs. S12 and S13 provide complete results for all 39 GEPs and risk tables.

**Figure 5.**
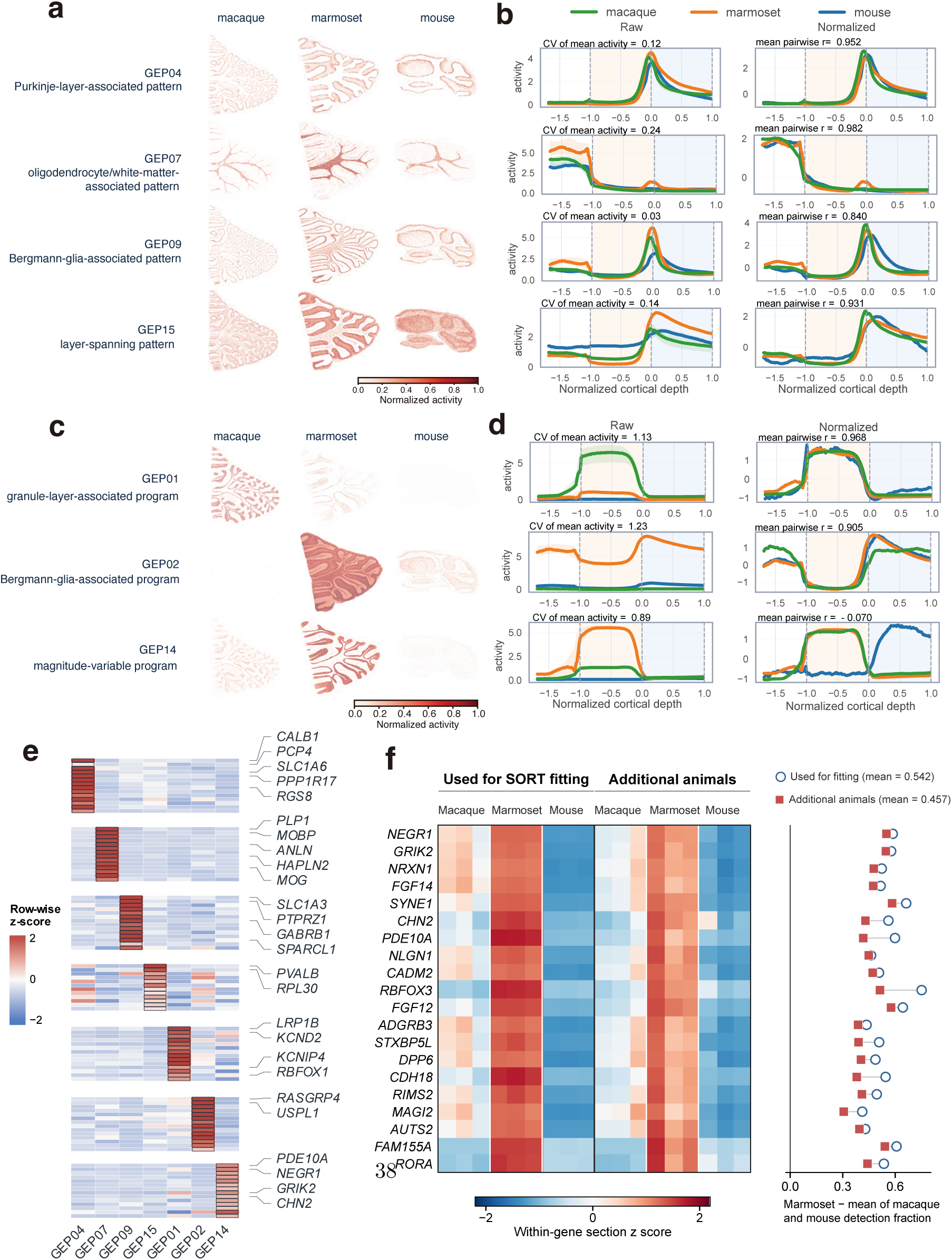
Conserved cerebellar layers and activity variation among selected specimens. **(a)** Spatial maps of selected GEPs associated with cerebellar layers in macaque, marmoset, and mouse. Color indicates normalized activity. **(b)** Cortical depth profiles over the interval between the 0.5th and 99.5th percentiles of the pooled cortical-depth coordinates. Raw profiles show activity magnitude, whereas profiles normalized within each GEP show spatial shape. Zero marks the Purkinje cell layer. Annotations report spatial profile similarity and the coefficient of variation (CV) of mean activity. **(c)** Spatial maps of selected GEPs with activity magnitude variation among the specimens. **(d)** Cortical depth profiles for the GEPs in **(c)**, calculated as in **(b)**. **(e)** Transcriptomic profiles for the seven GEPs shown in **(a,c)**. Rows are genes grouped by marker assignment or function, and columns are GEPs. Color shows profile values standardized within each gene. **(f)** Detection of GEP14-prioritized genes within GEP14 spatial domains in fitted and additional animals. Rows show the 20 prioritized GEP14 genes. Columns show three sections from one fitted and one additional animal of each species. Heatmap values are detection fractions standardized within each gene across all 18 sections. The adjacent plot shows the marmoset detection fraction minus the mean detection fraction in macaque and mouse.

**Figure 6.**
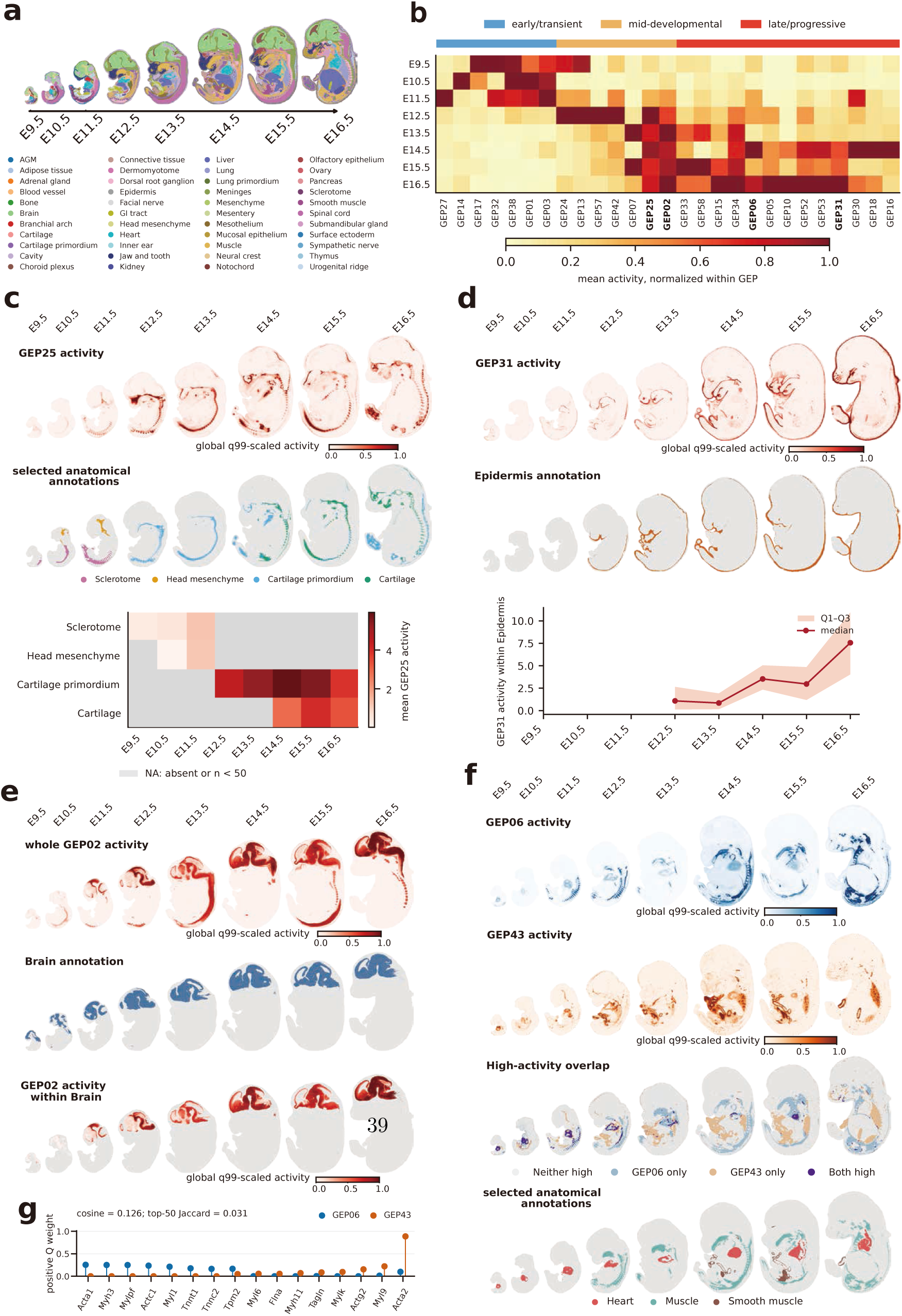
Graded and context-dependent program deployment during mouse embryogenesis. **(a)** Anatomical annotations across eight stages of mouse embryogenesis. **(b)** Stage-wise mean activity of 27 signal GEPs, normalized within each GEP and ordered from early peaks to intermediate maxima and later increases. **(c)** GEP25 spatial activity, selected anatomical annotations, and mean activity by stage and anatomical context. **(d)** GEP31 spatial activity and the epidermis annotation. Median activity and quartiles are shown for supported epidermis strata with at least 50 locations. **(e)** Whole-embryo GEP02 activity, the brain annotation, and GEP02 activity within brain locations. **(f)** GEP06 activity, GEP43 activity, high-activity overlap, and anatomical annotations for heart, muscle and smooth muscle. Activity maps are scaled by each GEP’s global 99th percentile and clipped at 1. The overlap map distinguishes locations high for GEP06 only, GEP43 only, both or neither, using each GEP’s 90th-percentile activity threshold within each stage. **(g)** Positive transcriptomic profile weights for selected high-weight genes from GEP06 and GEP43.

Formally, for each spatial section *s* = 1*,…, S*, let *X*^(*s*)^ ∈ R*^ns×p^* represent the expression of *p* genes in *n_s_* spatial locations. SORT solves the following optimization problem:

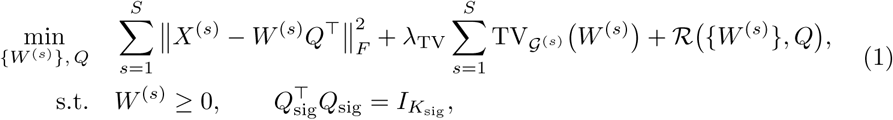

where *Q* = (*q*_0_*, Q*_sig_) ∈ R*^p×K^* contains the *K* shared transcriptomic profiles, with *q*_0_ as the background component and *Q*_sig_ as the *K*_sig_ = *K* − 1 signal profiles. The matrix *W^(s)^ = (w_0_^(s)^, W^(s)^_sig_) ∈ R^ns×K^_+_* contains their section-specific activity scores. Each signal GEP *k* thus comprises the transcriptomic profile *q_k_*and its corresponding spatial activity maps *{w^(s)^_k_}^S^_s=1_*, which define its gene-level identity and its deployment across sections, respectively. In the following, ‘GEP’ refers to signal GEPs unless otherwise noted.

The model is estimated by minimizing a compound objective that balances reconstruction fidelity with two regularization terms and the constraints. Graph total variation (TV) is applied independently on each section’s spatial graph G^(*s*)^ to encourage locally coherent activity while allowing sharp transitions at tissue boundaries. The auxiliary regularizer R(·, ·) promotes sparse signal profiles and activity scores, and discourages negative signal profile values. The orthogonality constraint *Q^T^_sig_Q_sig_ = I* reduces redundancy among signal GEPs by enforcing their linear independence. Unlike the nonnegative activity scores *W* ^(*s*)^, the signal profiles *Q*_sig_ can contain positive or negative weights within a profile, representing increases or decreases in their fitted expression as the GEP becomes more active. Because orthogonality is imposed on the profiles rather than the activity maps, distinct GEPs can remain active in the same spatial region. A detailed description of the model and optimization is provided in the Methods.

We first validated SORT on simulated data with patient-level effects designed to obscure weaker shared programs. We then applied SORT to three complementary biological contexts: a PDAC cohort, where we identified subtype-associated GEPs and evaluated survival associations with external bulk cohorts; a cross-species cerebellar atlas, where we separated conserved layer organization from species-specific activity variation; and a mouse organogenesis atlas, where we characterized recurrent, graded, and spatially overlapping GEPs across changing anatomical contexts. Across these diverse settings, SORT recovered shared transcriptomic programs and resolved their spatial deployment, enabling biological interpretation, cross-sample comparison, and integration with external clinical outcomes.

### SORT accurately recovers multilevel GEPs in simulated atlases

To benchmark recovery of multilevel GEPs, we generated simulated spatial transcriptomic atlases with known ground truth, where strong inter-patient heterogeneity was designed to obscure weaker programs shared across patient subtypes or tissue regions. Each of the 20 independent replicates comprised eight patient sections, with three spatial regions in each section. The focus region harbored eight patient-specific programs and three programs shared by patient subtypes, with two additional programs localized outside the focus. The ground truth thus comprised 13 signal GEPs per replicate in total. Replicates were generated with either annular or wedge-sector spatial layouts. Details of count generation, preprocessing, and the prespecified fitted rank are provided in the Methods.

We first illustrated the simulated structure using a representative replicate. The ground-truth spatial activity maps illustrated the designed localization of the 13 programs across regions (Fig. 4a). UMAP embedding of the expression data from the same replicate revealed a clear separation between the focus region and the other two (Fig. 4b). Within the focus region, patient labels defined eight finer branches, whereas subtype labels grouped these branches into three broader sets (Fig. 4c). This organization was consistent with the underlying hierarchy in which inter-patient variation dominated over subtype-level variation.

We next compared SORT against several baseline methods, including PCA, GraphPCA [10], NMF, NNMF [12] and STAMP [14], on the same representative replicate, evaluating the spatial activity maps of the inferred components that best matched the ground-truth GEPs (Fig. 4d and Supplementary Fig. S1). SORT accurately recovered the intended spatial localization of the displayed programs, with activity concentrated in their designated regions. NMF and NNMF tended to mix patient-specific and subtype-shared programs within the same components, blurring the hierarchical structure. PCA and GraphPCA yielded more diffuse activity maps that extended beyond the intended boundaries. STAMP recovered the subtype and region-associated patterns reasonably well, but spread patient-specific activities across multiple sections, failing to preserve their spatial specificity.

Beyond qualitative comparison, we quantitatively evaluated the recovery performance of SORT and the baseline methods by comparing their inferred spatial activity maps against the ground truth. Specifically, we computed the signed Pearson correlation between each ground-truth activity score vector and its best-matching inferred counterpart across all methods (Methods). Across 20 simulation replicates, SORT achieved consistently high recovery across all three program types, with median correlations of 0.994 for patient-specific programs (IQR, 0.992–0.996; *n* = 8 × 20 = 160), 0.987 for subtype-shared programs (IQR, 0.968–0.994; *n* = 3 × 20 = 60), and 0.992 for region-associated programs (IQR, 0.988–0.994; *n* = 2 × 20 = 40) (Fig. 4e). Among the baseline methods, the best median correlations were 0.952 for patient programs (NNMF), 0.857 for subtype programs (STAMP), and 0.982 for regional programs (STAMP). SORT showed its largest improvement over the best baseline for subtype programs, the level most strongly obscured by patient-specific variation. As a complementary specificity assessment, we compared, for each inferred subtype-shared GEP, its correlation with the matching ground-truth counterpart against its maximum correlation with any ground-truth patient-specific GEP. SORT showed the largest positive median difference, while NMF and NNMF yielded negative medians, suggesting that their subtype components were more correlated with patient-specific GEPs. The remaining methods showed smaller or more variable positive differences (Supplementary Fig. S2).

We further assessed boundary preservation in the recovered spatial activity maps using three complementary metrics: edge AUPRC, blur ratio, and Moran’s *I* (Methods). AUPRC measures boundary detection and higher is better, blur ratio measures activity-gradient spread around true boundaries and lower is better. We interpreted Moran’s *I* alongside the boundary metrics, as excessive smoothing can inflate spatial autocorrelation without preserving true boundaries. SORT achieved the best boundary preservation performance among all methods, with the lowest blur ratio (median 0.263, IQR 0.234–0.280, *n* = 20) and the highest edge AUPRC (median 0.684, IQR 0.677–0.872, *n* = 20). STAMP ranked second on both metrics (blur ratio 0.775; edge AUPRC 0.473). In contrast, NNMF produced the highest Moran’s *I* but performed poorly on boundary metrics (blur ratio 2.148; edge AUPRC 0.349) (Fig. 4f), illustrating that high spatial autocorrelation alone does not indicate accurate boundary recovery.

To assess the sensitivity of SORT to its hyperparameters and model components, we performed matched sensitivity and ablation analyses using ten simulation seeds. We first varied the graph TV weight *λ*_TV_ across a range of values. Recovery was highest and spatial blurring was lowest at *λ*_TV_ = 1.0, with edge AUPRC remaining similar to that at 0.6 (Supplementary Fig. S3). Lower TV weights reduced subtype recovery and increased spatial blurring, with the largest deterioration at *λ*_TV_ = 0.3, suggesting that graph total variation is critical for accurately recovering spatially coherent programs. We then evaluated the contribution of individual model components by systematically removing the graph TV penalty, the sparsity penalty on *W* ^(*s*)^, or both. Removing graph TV, either alone or together with the sparsity penalty, substantially reduced both recovery accuracy and boundary preservation, underscoring the importance of spatial regularization. In contrast, removing the sparsity penalty on *W* ^(*s*)^ alone had little effect on performance (Supplementary Fig. S4), indicating that activity sparsity played a secondary role in these simulations. To further examine the contribution of the orthogonality constraint *Q^⊤^ Q*_sig_ = *I*, we compared SORT against a variant that removed pairwise orthogonality while retaining unit norm constraints (Methods). Removing orthogonality had little effect on recovery of patient-specific and region-associated programs, both with correlations near 0.99. In contrast, median correlation of subtype-shared programs dropped substantially from 0.963 to 0.660. Relative reconstruction error remained nearly unchanged. Boundary metrics showed mixed results: edge AUPRC favored the original SORT, whereas blur ratio favored the variant (Supplementary Fig. S5). These results indicate that orthogonality primarily reduces redundancy among weaker shared programs, such as those at the subtype level, whereas boundary preservation is governed by graph total variation.

Together, these simulations show that SORT recovered nonredundant programs across three levels of shared specificity while preserving spatial boundaries when patient variation dominated.

### SORT resolves PDAC programs with variable spatial deployment and survival associations across cohorts

Pancreatic ductal adenocarcinoma (PDAC) is characterized by substantial interpatient heterogeneity, with tumors exhibiting variable proportions of basal-like and classical transcriptional states and diverse stromal compositions that shape the tumor microenvironment. Resolving how these GEPs are spatially organized across patients can reveal the joint deployment of tumor and microenvironmental programs and inform studies of PDAC progression and therapeutic response. To investigate this, we fitted SORT to a published atlas of treatment-naive primary tumors containing 108 Visium sections from 39 patients and 352,102 retained spatial locations [4].

Before analyzing the full atlas, we first benchmarked SORT against baseline methods on their ability to recover the basal-like and classical reference signals reported by the original study. UCell scores computed from the published signatures served as reference signals [4, 15]. To assess the robustness of SORT across patient and section composition, we constructed 20 stratified subsets, each containing ten sections from ten patients (Methods). As shown in a representative subset, SORT activity maps closely followed the spatial distribution of high basal-like and classical reference scores across patients (Fig. 4a). In contrast, recovered spatial activities from the comparison methods were either more restricted to individual sections or showed weaker concordance with the reference scores, suggesting that SORT more reliably recovered the published subtype reference patterns. To quantify this, we computed the correlation between each inferred activity score vector and the reference signals for each of the 20 subsets. SORT consistently showed the strongest agreement with both subtype references. Median Pearson correlations with the basal-like reference were 0.739 for SORT and 0.670 for STAMP, the best-performing baseline method (*P* = 5.72 × 10*^−^*^6^, paired Wilcoxon). For the classical reference, the medians were 0.755 for SORT and 0.627 for STAMP (*P* = 1.91 × 10*^−^*^6^, paired Wilcoxon; Fig. 4b). As a complementary specificity check, we compared each inferred activity’s correlation with its intended subtype reference against its correlation with the alternative. SORT achieved the highest target specificity for both basal-like and classical subtypes (Supplementary Fig. S6).

We next applied SORT to the full dataset of 108 sections. The fit recovered 39 signal GEPs, capturing a range of biological processes including epithelial states and stromal, vascular and immune programs, as indicated by their associations with published PDAC and tumor microenvironment signatures. Over-representation analysis (ORA) of the top positive genes in *Q* provided functional annotations (Methods). Summary of the ten top positive-weight genes in *Q* and representative enriched terms for all 39 GEPs is provided in Supplementary Table 1 (Methods). To provide a global overview of the fitted output, Supplementary Fig. S7 reports the mean activity score and the fraction of active locations for each of the 39 signal GEPs across all sections. Spatial activity maps for a representative panel of 16 sections, selected independently of GEP activity, are shown in Supplementary Fig. S8.

To quantify the association of each gene with each recovered GEP while controlling for other programs, we regressed normalized expression for each gene on all fitted activity maps in *W*_sig_, yielding coefficients *B_gk_*(Methods). Standardized *B_gk_*values for 74 representative genes from published PDAC and tumor microenvironment signatures reveal specific gene–GEP–signature relationships (Fig. 4a) [4]. For example, GEP02 captured a pancreatic exocrine program enriched in digestion and absorption, and GEP30 had high positive profile weights for exocrine-associated genes such as *CTRB2*, *REG3A*, *CELA3A*, and *PRSS2*. The expression of these genes showed strong associations with GEP02 in *B_gk_*, indicating shared exocrine characteristics between GEP02 and GEP30. GEP29 captured a pancreatic ductal program, marked by ductal-associated genes such as *SLC4A4*, *SERPINA5*, *CFTR* and *SORBS2*. *SLC4A4* and *CFTR* are involved in ductal ion transport [16, 17], whereas *SERPINA5* and *SORBS2* are associated with ductal features and cell adhesion and migration, respectively [4, 18]. To assess associations between each signature and each GEP, we calculated the signed mean of *B_gk_*across overlapping genes and z-scored the resulting values across the 14 displayed GEPs (Fig. 4b; Methods). These analyses linked GEP04 to activated stroma and myofibroblastic cancer-associated fibroblast (myCAF) signatures, consistent with its enrichment in extracellular matrix organization, collagen formation, integrin cell surface interactions, and epithelial–mesenchymal transition (Supplementary Fig. S10a). GEP11 is linked to T-cell and B-cell signatures, with top positive-weight immune-related genes such as *PTPRC* and *IL7R*, and enriched in adaptive immune system and cytokine signaling. Supplementary Fig. S9 extends the signature summaries to all 39 signal GEPs.

Among the epithelial GEPs, pathway, signature and gene-level associations distinguished a classical program with secretory features from basal-like programs with different accompanying features. GEP08 was enriched in protein secretion and O-linked glycosylation of mucins (Supplementary Fig. S10c; Methods). It also showed a positive projection onto the tumor classical signature and positive gene-level associations with *AGR2*, *TFF1*, *LGALS4*, and *TFF3* (Fig. 4a,b). These transcriptional and functional features were consistent with the mucus-producing classical state of PDAC [4, 19]. GEP01, GEP16 and GEP32 all projected positively onto the tumor basal-like signature, but exhibited distinct gene-level and functional features. GEP01 included top-ranked basal-like-associated genes such as *KRT7* and *KRT17*, and showed gene-level associations with *FAM83A*, *CST6* and *SCEL*. It was enriched in keratinization and type I hemidesmosome assembly, highlighting a keratinized epithelial program. GEP16 showed gene-level associations with *KRT7*, *S100A2*, *TNS4* and *KRT6A*, and was enriched in epithelial–mesenchymal transition, keratinization and hypoxia. Similarly, GEP32 had high positive profile weights for *KRT17* and hypoxia-associated *SLC2A1*, and showed a gene-level association with *S100A2*, *KRT6A* and *ANXA8L1*, with enrichment in hypoxia and glycolysis, highlighting a hypoxia-related metabolic component of the basal-like program in PDAC [4, 20]. ORA showed concordant enrichment for these processes (Supplementary Fig. S10e). Together, the integrated annotations distinguished a classical PDAC tumor program with secretory features from basal-like programs with distinct keratinized, mesenchymal and hypoxia-related features.

We next examined whether the transcriptionally distinct programs showed different degrees of spatial colocalization. To illustrate this, we examined the spatial overlap of activity for two representative pairs: the classical-associated GEP08 with the basal-associated GEP16, and the ductal-associated GEP29 with the acinar-associated GEP30 (Fig. 4c). For each section, locations with activity score at or above the 80th percentile were classified as occupied by either program or by both. We calculated a composite colocalization score summarized from high-activity overlap and activity correlation within each section (Methods). For GEP08 and GEP16, the displayed two low-colocalization sections had scores of 0.244 and 0.344, and the two high-colocalization sections had scores of 0.875 and 0.898, respectively. For GEP29 and GEP30, the corresponding scores were 0.159 and 0.167, and 0.663 and 0.770. This highlights that the degree of spatial colocalization between the same two programs can vary widely across sections and between program pairs. To further explore this heterogeneity, we compared GEP16 and GEP32 across the complete atlas (Supplementary Fig. S11). Although these two programs were transcriptionally distinct, their spatial activity correlations varied considerably across sections. Sections with stronger correlations also exhibited greater overlap between high-activity locations for both GEPs. These results demonstrate that transcriptionally distinct programs can exhibit a spectrum of spatial relationships, from largely segregated to extensively colocalized, depending on the local tissue context.

To assess whether the recovered GEPs had clinical relevance, we next evaluated associations between the spatially resolved programs and patient outcomes in four external bulk PDAC cohorts. For each GEP, we computed a bulk projection score for each patient using a fixed gene set selected prior to any survival analysis (Methods). All 39 scores corresponding to the 39 GEPs were then tested for association with overall survival within each cohort using Cox proportional hazards models adjusted for available covariates, and the cohort-level effect estimates were combined via fixed-effect meta-analysis. Benjamini–Hochberg correction controlled the false discovery rate (FDR) across all 39 GEPs, with cohort details and full numerical results reported in Supplementary Table 2. Fig. 4d,e summarizes the ten GEPs with the strongest associations (smallest FDR values), and Supplementary Fig. S12 visualizes the full screen across all 39 GEPs, including FDR values, cohort-specific hazard ratios (HRs), and *I*^2^ statistics quantifying heterogeneity across cohorts.

Higher bulk projection scores for GEP21, associated with hypoxia and stress, and GEP16, associated with basal features, were associated with shorter survival. Their meta-analysis HRs were 1.482 and 1.408, with FDR values of 4.74 × 10*^−^*^7^ and 3.65×10*^−^*^5^, respectively. The corresponding *I*^2^ values were 25.2% and 47.1%. Pathway annotation for GEP21 highlighted hypoxia and glycolysis, with selected genes from *Q* including *DDIT* 4, *GAPDH*, *ALDOA* and *TPI*1 (Supplementary Fig. S10b). By contrast, higher scores for GEP08 and GEP22 were associated with longer survival (HR 0.665 and 0.689; FDR 3.17 × 10*^−^*^6^ and 1.47 × 10*^−^*^5^; *I*^2^ = 0% for both; Fig. 4d,e). GEP08 showed tumor classical associations. GEP22 showed modest positive projections for macrophage, B-cell and ductal signatures (Supplementary Fig. S9), while its effect estimates were protective across all four cohorts.

For Kaplan–Meier visualization, we selected GEP21, GEP08, GEP22 and GEP16, the four GEPs with the smallest FDR values, and stratified patients by the median of the corresponding bulk projection scores standardized within each cohort. This stratification produced the expected adverse or protective survival separation for each GEP, consistent with the direction of its meta-analysis HR (Fig. 4f). Supplementary Fig. S13 shows the four complete curves with censor marks and numbers at risk. These curves complement the continuous Cox models and meta-analysis. Supplementary Fig. S14 applies the same pooled median split to all 17 GEPs with FDR *<* 0.05. These cross-cohort associations support further evaluation of the prognostic relevance of the corresponding programs.

Together, SORT recovered recurrent tumor state and microenvironmental GEPs across patients in one common profile space, resolved variation in their spatial overlap among sections, and linked their bulk projections to survival across external cohorts.

### SORT distinguishes conserved cerebellar layer organization from spatial activity variation across species

The cerebellum is a highly conserved brain structure across vertebrates, yet the specific transcriptional programs that are conserved or species-specific are not fully resolved. To address this, we applied SORT to a large cross-species cerebellar atlas of macaque, marmoset and mouse, jointly analyzing nine Stereo-seq sections comprising three adjacent sections from one animal per species to compare shared layer-associated organization and activity differences among the sampled animals [5]. The dataset contained 1,546,637 spatial locations and 4,000 genes retained for model fitting. To facilitate spatial interpretation, we aligned the recovered GEP activities to the cortical depth coordinate provided by the atlas, with the Purkinje cell layer at zero. Supplementary Fig. S15 displays spatial activity maps for all 25 fitted GEPs across the nine sections. Supplementary Table 3 provides the summary of the top positive-weight genes and enriched terms for the 24 signal GEPs (Methods).

SORT identified four signal GEPs with conserved layer-associated spatial patterns across the three sampled animals (Fig. 4a). GEP04 localized to the Purkinje cell layer, GEP07 to oligodendrocyte-rich white matter and GEP09 to Bergmann glia. GEP15 showed a broader pattern spanning several layers. Their normalized cortical depth profiles remained similar across the three animals over a common interval of 160 bins, with mean pairwise correlations ranging from 0.840 to 0.982. Their mean activities also varied little, with coefficients of variation (CVs) ranging from 0.03 to 0.24 (Fig. 4b; Methods).

By contrast, GEP01, GEP02, and GEP14 showed larger differences in activity magnitude among the sampled animals (Fig. 4c,d; Methods). GEP01 and GEP02 retained similar spatial depth profiles across species, with mean pairwise correlations of 0.968 and 0.905, respectively, but their activity strengths varied markedly: GEP01 was strongest in macaque, whereas GEP02 was strongest in marmoset, with CVs of mean activity score of 1.13 and 1.23, respectively. GEP14 was strongest in marmoset, weaker in macaque and nearly undetectable in mouse, with a CV of 0.89. These CV values are substantially higher than those of the conserved layer-associated GEPs. Supplementary Fig. S16 ranks all 25 GEPs by CV and shows the section-level means underlying the seven profiles discussed here.

Transcriptomic profiles of the seven displayed GEPs provided gene-level evidence for their spatial deployment (Fig. 4e). The conserved layer-associated GEPs contained markers consistent with their spatial localizations. GEP04 combined the calcium-related genes *CALB1* and *PCP4*, together with *SLC1A6*, a glutamate transporter expressed by Purkinje cells [21], and was enriched for synaptic signaling, transport along microtubules, and neuron development. GEP07 contained the oligodendrocyte and myelin genes *PLP1*, *MOBP* and *MOG*, consistent with its activity in white matter [22], and was enriched for myelination, oligodendrocyte development, and glial cell development. GEP09 contained *SLC1A3* and the extracellular matrix gene *SPARCL1*. *SLC1A3* mediates glutamate clearance around synapses [23], and GEP09 was enriched for positive regulation of synaptic transmission, neuron projection development, and gliogenesis. The GEPs with greater activity variation (GEP01 and GEP14) showed distinct neuronal gene combinations. GEP01 included genes *KCND2*, *KCNIP4* and *RBFOX1*, and was enriched for synaptic signaling, neuron projection development, and synaptic vesicle cycle. GEP14 included *GRIK2*, *CHN2* and *NEGR1*, and was enriched for synaptic membrane adhesion, synapse organization, and synaptic signaling (Supplementary Table 3). These gene and pathway results supported the biological interpretation of the displayed GEPs.

We next examined whether the activity differences for GEP02 and GEP14 were accompanied by concordant detection patterns for genes prioritized from their profiles. For each GEP, we fixed the 20 genes with the largest positive profile weights among those meeting the prespecified selection criterion before cross-species evaluation (Methods). High-activity domains were defined by fitted GEP activities in the original animals. In one additional animal per species, gene-masked projection onto the original GEP profiles located corresponding high-activity domains without using the evaluated genes or refitting SORT (Methods). Within these domains, we compared the detection fractions of the prioritized genes across the sampled animals. For GEP14, all 20 prioritized genes showed higher detection fractions in marmoset than the mean of macaque and mouse in both the original and additional animal sets (Fig. 4f). The mean differences were 0.542 in the animals used for SORT fitting and 0.457 in the additional animals. GEP02 showed the same direction for all 20 genes, with mean differences of 0.408 and 0.225 in the two animal sets (Supplementary Fig. S17a). For both GEPs, the mean detection difference across the 20 selected genes exceeded those of all 10,000 matched control sets and persisted after restriction to confirmed one-to-one orthologues (Supplementary Fig. S17b,c). These results confirm that species with higher GEP activity showed higher detection fraction of the corresponding genes. Supplementary Fig. S17d provides a separate reciprocal contrast within the shared granule layer domain of GEP00 and GEP01. Genes prioritized from GEP01 showed greater detection in macaque, where GEP01 activity was also greater. These analyses linked GEP deployment to coherent gene detection patterns in both the fitting and additional animal sets.

The runtime across six balanced input sizes, including the complete atlas, is shown in Supplementary Fig. S18. On a single NVIDIA A30 graphics processing unit (GPU), SORT processed the complete nine-section atlas, comprising 1,546,637 spatial locations and 4,000 genes, in 25.0 minutes end to end.

Together, the cerebellar analysis demonstrated that a single SORT fit can separate conserved layer organization from species-specific activity differences, link selected GEPs to gene detection within corresponding spatial domains, and scale to more than 1.5 million locations.

### SORT resolves graded and context-dependent spatial deployment of GEPs during mouse embryogenesis

Mammalian organogenesis involves dynamic transcriptional programs precisely coordinated across space and time. Characterizing these programs and their spatial organization is essential for understanding tissue patterning and cell fate specification. To systematically investigate this, we applied SORT to a mouse organogenesis atlas spanning eight cross-sectional stages from E9.5 to E16.5, with 5,902–121,752 retained spatial locations per stage after filtering [6] (Fig. 4a; Methods). Joint fitting using SORT recovered 59 signal GEPs. Their top positive-weight genes and enriched terms are summarized in Supplementary Table 4 (Methods).

Across development, SORT revealed shared transcriptomic profiles whose spatial deployment recurred across changing anatomical contexts and formed graded patterns within developing tissues. The 27 GEPs displayed in Fig. 4b showed diverse temporal trajectories, including sharp early peaks, intermediate maxima, and sustained increases at later stages. For example, GEP27 peaked at E11.5, GEP25 increased from E12.5 onward, and GEP31 and GEP06 reached their highest mean activities later in development. Supplementary Fig. S19 shows the temporal trajectories of all 59 signal GEPs. Supplementary Fig. S20 maps every fitted component across all eight stages and shows spatial domains ranging from broad embryonic regions to individual organs and narrow tissue layers.

GEP25 activity marked the E11.5 sclerotome and head mesenchyme, increased in the E12.5 cartilage primordium, and remained high in later cartilage-related compartments (Fig. 4c). To quantify this association, we compiled a literature-informed gene set comprising the chondrogenic regulators *Sox5*, *Sox6* and *Sox9* and the cartilage matrix genes *Acan*, *Col2a1*, *Col9a1* and *Col11a1* [24, 25] (Supplementary Table 2 documents the gene set membership, biological rationale, and literature sources). We summarized the set by averaging gene-wise z-scored log expression. Across nine combinations of developmental stage and anatomical annotation, correlations with GEP25 activity ranged from 0.774 to 0.941 after adjustment for detected-gene coverage (Supplementary Fig. S21a; Methods). Gene-set scores also increased across ten equally sized groups ordered by GEP25 activity in representative precursor, primordium and later cartilage contexts (Supplementary Fig. S21b). A complementary ORA of genes with positive profile values in *Q* emphasized cartilage development, collagen formation, and extracellular matrix organization (Supplementary Fig. S22a), which are consistent with the enriched terms for GEP25 (Supplementary Table 4). Together, the concordant spatial, gene-set, and pathway patterns support a chondrogenic-associated interpretation of GEP25, from early mesenchymal to later cartilage-related compartments.

The common transcriptional profile space also resolved developmental changes within a single anatomical structure. Within the epidermis, we assessed GEP31 activity against a fixed epithelial-associated gene set that included the basal keratins *Krt5*, *Krt14* and *Krt15*, along with the adhesion-associated genes *Perp* and *Dsp* [26, 27]. Median GEP31 activity increased overall from 1.09 at E12.5 to 7.56 at E16.5 (Fig. 4d). Coverage-adjusted rank correlations between GEP31 activity and this gene set ranged from 0.782 to 0.887 across all five supported stages (Supplementary Fig. S21c,d; Supplementary Table 2). ORA of genes with positive profile values also emphasized keratinization and epidermis development (Supplementary Fig. S22b). These results show that GEP31 activity increased steadily during epidermal development, capturing stage-associated dynamics within a stable anatomical region. GEP31 was correspondingly enriched for keratinocyte differentiation, epidermis development, and keratinization.

GEP02 resolved graded spatial variation within the broad brain annotation across all eight stages (Fig. 4e). To relate this spatial variation to neural gene expression, we compiled a gene set that included the neuronal differentiation genes *Map2* and *Dcx*, along with the cytoskeletal genes *Mapt* and *Ina*, selected as established neural markers from the literature [6, 22]. Coverage-adjusted rank correlations between GEP02 activity and this gene set ranged from 0.345 to 0.870 across the eight stages, with weaker support at E9.5 and E10.5 and stronger support from E11.5 onward. Higher GEP02 activity remained associated with higher gene-set scores across alternative activity thresholds and spatial grid resolutions (Supplementary Fig. S23). ORA of genes with positive profile weights in *Q* further emphasized neuronal differentiation, axonogenesis, and nervous system development (Supplementary Fig. S22c), consistent with the enriched terms for GEP02, which include neuron projection morphogenesis, axonogenesis, and synapse organization. These results show that GEP02 captured graded neural-associated expression, revealing spatial heterogeneity within the broad brain region.

Transcriptomically distinct programs could overlap spatially at particular developmental stages. GEP06 and GEP43 illustrated this pattern, with overlapping activity in the heart during E9.5–E12.5 despite their different overall spatial distributions (Fig. 4f). Across whole sections, their colocalization score decreased from 0.644 at E9.5 to 0.260 at E12.5 and remained lower at later stages (0.118–0.161, Supplementary Fig. S24b). Their positive-profile cosine similarity was 0.126, and their top-50 positive gene sets had a Jaccard overlap of 0.031 (Fig. 4g), indicating that their transcriptomic profiles were largely distinct. This distinction was also reflected in their gene compositions: GEP06 contained sarcomeric and excitation–contraction genes, including *Myh*3, *Tnnt*2, and *Ryr*1, whereas GEP43 contained smooth-contractile genes, including *Acta*2, *Tagln*, and *Myh*11 [28, 29]. Within each of the eight stages, both GEP activities correlated positively with their respective fixed gene-set expression scores after adjustment for detected-gene coverage (Supplementary Fig. S24c; Methods). ORA of genes with positive profile loadings in *Q* emphasized striated muscle contraction for GEP06 and actomyosin and smooth-contractile processes for GEP43 (Supplementary Fig. S22d,e).

Across mouse organogenesis, the common profile space recovered GEPs that recurred across anatomical contexts, changed in activity across stages, formed graded patterns within tissues and remained transcriptomically distinct despite spatial overlap. These findings show how one shared decomposition can resolve multiple modes of developmental program deployment.

## Discussion

SORT provides a unified computational framework for jointly recovering shared and context-specific transcriptional programs across multi-section spatial atlases while preserving tissue spatial structure. By factorizing multiple sections into a common transcriptional profile space, it resolves programs from sample-specific and group-shared patterns to region-associated signatures, with graph total variation to maintain spatial coherence and orthogonality to reduce redundancy. In doing so, the framework systematically distinguishes programs by their level of sharing and spatial deployment, enabling each level of program to be interpreted in its own biological context.

The simulations validated the capability of SORT against known ground-truth programs. It achieved its largest advantage for weaker subtype-related programs when patient-level variation dominated, and ablation analyses linked orthogonality to nonredundant recovery and graph total variation to boundary preservation, whereas activity sparsity had a smaller effect on performance. Across the three biological atlases, SORT demonstrated its ability to recover biologically interpretable programs in PDAC, the cross-species cerebellum and mouse organogenesis. In PDAC, SORT separated a classical secretory program from basal-like programs with keratinization, mesenchymal and hypoxia-related features, and linked several programs to survival across four external cohorts using fixed bulk projections. In the cerebellum, recurrent layer organization was distinguished from activity differences among the sampled animals. During organogenesis, the common profile space linked chondrogenic activity across changing anatomical compartments, resolved changes across stages in the epidermis and graded neural patterns, and distinguished sarcomeric and smooth-contractile programs with local spatial overlap. Gene-level associations and positive-profile enrichment provided complementary support for selected annotations.

SORT also scaled to large input sizes in this study under the tested configuration. Supplementary Fig. S18 shows the observed runtime across six input sizes from 51,997 to 1,546,637 spatial locations. The complete 1.55-million-location cerebellar atlas required 25.0 minutes end to end on one NVIDIA A30 GPU.

The common profile space in SORT also provides a foundation for addressing several more complex biological questions. A natural extension is to model sample or condition covariates directly on GEP activities. This would allow formal comparison of how a shared transcriptomic program is deployed across disease states, treatments or developmental stages while retaining a common profile identity. Another is adapting the framework to count-based data, where likelihoods such as negative binomial or Poisson could better handle sparse or shallowly sequenced assays. Spatial graphs could also be extended to incorporate histological images, additional molecular modalities, or weak biological priors to guide regularization. Biological replication across species, lineage-traced developmental data and prospective clinical cohorts would extend SORT to population-level, lineage-specific and prognostic inference. Together, these extensions would preserve the core idea of SORT while broadening its applicability to richer data types and more challenging biological questions.

## Methods

### SORT: spatially regularized transcriptomic decomposition with orthogonal constraints

SORT jointly analyzes multiple spatial transcriptomic sections by decomposing them into shared transcriptomic profiles and section-specific spatial activity scores that describe program deployment within each section and variation across sections. For section *s* = 1*,…, S*, *X*^(*s*)^ ∈ R*^ns×p^* contains the expression of *p* genes at *n_s_* spatial locations. Dataset-specific filtering, normalization and feature selection are described in the corresponding analysis subsections and summarized in Supplementary Table 3.

For each spatial section *s*, SORT approximates its expression matrix as

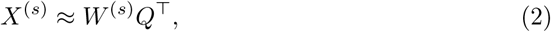

where *Q* = (*q*_0_*, Q*_sig_) ∈ R*^p×K^* contains the shared transcriptomic profiles. Here, *q*_0_ is the background profile, and the remaining *K*_sig_ = *K* − 1 columns of *Q*_sig_ are signal profiles. Correspondingly, *W^(s)^ = (w_0_^(s)^, W^(s)^_sig_) ∈ R^ns×K^_+_* contains the spatial activity scores for these profiles within section *s*, where *w^(s)^_0_* and the columns of *W^(s)^_sig_* encode the activities of the background and signal profiles, respectively. Each signal GEP *k* comprises transcriptomic profile *q_k_*and its activity maps *{w^(s)^_k_}^S^_s=1_* across sections.

SORT jointly factorizes the expression matrices of all sections by solving the following optimization problem:

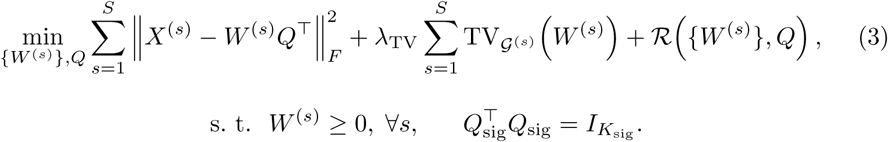

Here, TV*_G_*_(_*_s_*_)_ denotes the graph total variation penalty, which promotes spatial coherence of activity maps within each section while allowing sharp transitions at tissue boundaries. The regularizer R(·, ·) is defined as

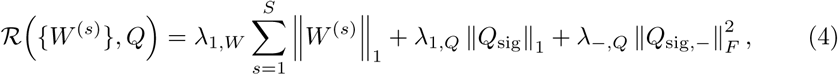

where ||·||_1_ and ||·||*_F_* denote the element-wise *ℓ*_1_ and Frobenius norms, respectively, and the *ℓ*_1_ penalties promote sparsity in all the activity scores and the transcriptomic profiles of the signal GEPs. *Q*_sig,_*_−_* = min(*Q*_sig_, 0) applies an element-wise penalty to negative entries in *Q*_sig_, shrinking negative signal-profile values without imposing a nonnegativity constraint. The hard constraint *Q^T^_sig_ Q_sig_ = I_Ksig_* enforces column-wise orthogonality among the signal profiles, reducing redundancy and promoting diversity among the learned transcriptional programs [30].

#### Spatial graph total variation

For each section, we constructed a symmetric *k*-nearest-neighbor graph using the Euclidean distances between spatial coordinates, defining the adjacency structure over which the graph total variation (TV) penalty is evaluated. Let *A* denote the binary connectivity matrix, *D* its degree matrix, and define

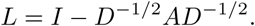

For *i* ≠ *j*, the TV-specific adjacency weight is given by *A^TV^_ij_ = max(−L_ij_, 0)*. For section *s*, let *E^(s)^ = {(i,j): i<j, A^(s)^_ij_ = 1}* denote its set of undirected edges. To normalize the edge weights across sections, we defined the static weight for each edge (*i, j*) ∈ *E*^(*s*)^ as

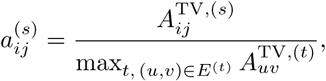

where the denominator is taken as the single maximum over the combined block-diagonal graph across all sections. Finally, the componentwise TV penalty is given by

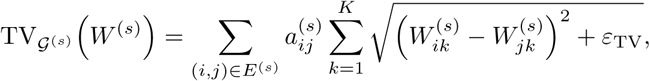

where *ε*_TV_ *>* 0 is a small constant that stabilizes the approximation of the absolute difference [31].

#### Alternating optimization

To minimize the objective in equation (3), SORT alternates between the section activity score matrices *W* ^(*s*)^ and the shared transcriptomic profile matrix *Q* = (*q*_0_*, Q*_sig_). With *Q* fixed, each activity score matrix *W* ^(*s*)^ is updated by solving

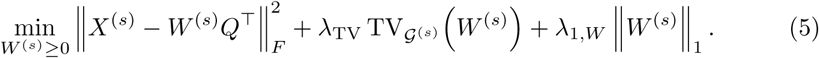

The nonnegativity of *W* ^(*s*)^ is maintained by multiplicative updates. At each update, the graph TV term is replaced by an iteratively reweighted least-squares (IRLS) quadratic majorizer calculated from the current edgewise activity differences [32].

With the activity score matrices fixed, the background profile *q*_0_ is estimated via unconstrained least squares by solving

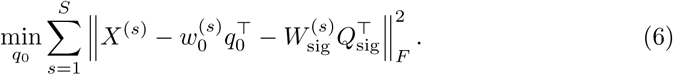

The signal profiles *Q*_sig_ are updated in two stages. Stage 1 computes an unpenalized orthogonal Procrustes solution, which serves as an initialization. Stage 2 then incorporates sparsity and the penalty on negative values, and optimizes the resulting objective on the Stiefel manifold. To formulate the signal profile update subproblems, the data matrices and activity score matrices are stacked by rows:

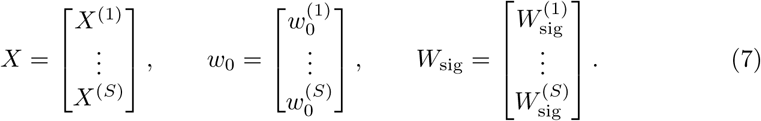

Let *R_0_ = X − w_0_q_0_^T^* denote expression after subtracting the fitted background component. Stage 1 solves

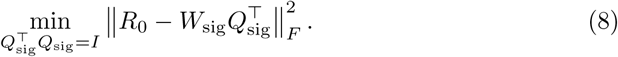

If *R^T^_0_W_sig_ = UΣV^T^*, the orthogonal Procrustes solution is *Q*_sig_ = *UV ^⊤^*.

Stage 2 instead optimizes the full signal profile subproblem

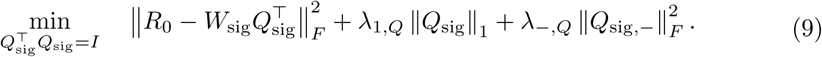

The signal profile gradient is projected onto the tangent space of the Stiefel manifold, and each Adam update is followed by a polar retraction that returns the profile matrix to the manifold [33]. This maintains *Q^T^_sig_ Q_sig_ = I*.

Supplementary Note 1 provides the IRLS derivation, tangent projection, retraction, initialization, update schedule, prespecified epoch numbers, component-specific sparsity weighting and implementation details. Supplementary Algorithm 1 summarizes the complete update sequence. Parameter values for each analysis are listed in Supplementary Tables 3–4.

#### Guidance for the rank selection

We recommend using the elbow *k*_0_ of a principal component analysis (PCA) log-eigenvalue scree curve to guide the factorization rank. The curve is calculated from a balanced row sketch of the same preprocessed expression matrix *X* used for SORT. A practical default for the rank *K* is *K* ≈ 2*k*_0_, with larger values providing finer resolution for more heterogeneous datasets and smaller values giving coarser representations of simpler datasets. Supplementary Fig. S25 shows the scree curves and *k*_0_ estimates for the three biological atlases and the 20 simulated datasets. These diagnostics provide a reference scale for selecting *K* in practice.

### Simulation benchmark

#### Simulation design

We simulated eight patient sections on 36 × 36 grids, giving *N* = 10,368 spatial locations and *J* = 8,000 genes. Baseline counts were generated with splatter::splatSimulate using parameters estimated from a reference PDAC dataset [34], an outlier-gene probability of 0.005 and a log library size shift of log(2.5). We then used seqgendiff::thin diff [35] to inject 13 signal GEPs. Their spatial activities were represented by *W*_true_ ∈ R*^N×^*^13^ and their gene effects by *Q*_true_ ∈ R*^J×^*^13^. The product *W_true_Q^T^_true_* specified the location-by-gene signal used to perturb the Splat-generated baseline counts.

The injected programs represented three levels of shared specificity. A designated focus region contained eight patient-specific programs and three subtype-related programs. Two region-related programs occupied areas outside this region. Their nonoverlapping marker sets contained 500, 100 and 75 genes, respectively, with mean log_2_ fold changes of log_2_(3.0), log_2_(3.25) and log_2_(3.0). The region layouts were annular or wedge-sector. We generated 20 independent replicates, comprising nine annular and 11 wedge-sector layouts. Supplementary Fig. S26 shows both layout families.

#### Data preprocessing and model fitting

Genes expressed in fewer than 32 locations were removed, after which 5,000 highly variable genes were selected for model fitting. Counts were normalized with scran size factors using quickCluster and calculateSumFactors and then log-transformed [36]. The exact transformation is given in Supplementary Note 2.

All methods were fitted at the same nominal rank of 24 where applicable. For SORT, this comprised one background component and 23 signal components. As the simulated ground truth contained 13 signal GEPs, the fitted rank was overspecified relative to the recovery targets, allowing unmatched or redundant components rather than enforcing a one-to-one decomposition. Baseline inputs and shared simulation settings are listed in Supplementary Tables 3–6.

#### Evaluation metrics

Recovery was evaluated using spatial activity maps. For each injected GEP, locations from all eight sections were pooled, and the true and inferred activity vectors were min–max scaled over these pooled locations. The inferred component with the highest signed Pearson correlation coefficient (PCC) was selected independently for each true program, and the corresponding PCC was recorded. The procedure evaluated each injected pattern without imposing a one-to-one assignment.

Boundary metrics used horizontal and vertical edges connecting adjacent locations within each 36 × 36 section grid. True boundaries were the top 10% of nonzero edge gradients in the ground truth. To compute the edge area under the precision–recall curve (AUPRC), true boundary edges were treated as positives and the corresponding inferred activity gradients were used to rank all edges. The blur ratio divided the sum of inferred gradients on neighboring nonboundary edges by the sum on true boundary edges. Higher edge AUPRC and lower blur ratio indicate more faithful boundaries. Supplementary Note 2 gives the complete definitions. Moran’s *I* summarized similarity between neighboring locations. It was interpreted together with edge AUPRC and blur ratio because spatial autocorrelation alone does not distinguish coherent recovery from oversmoothing.

Recovery was computed for each of the 13 injected programs, whereas boundary metrics were averaged across programs within each simulation replicate. Full formulas and numerical settings are provided in Supplementary Note 2 and Supplementary Table 7.

#### Sensitivity and ablation

All sensitivity and ablation analyses used the same 10 matched simulation seeds across the comparisons. We evaluated a local grid around the main simulation setting, with *λ*_TV_ ∈ {0.3, 0.6, 1.0} and *λ*_1*,W*_ ∈ {0.3, 0.5, 0.7}, while keeping the remaining settings fixed.

To isolate the contribution of graph TV and *W* sparsity, we also evaluated three ablation settings: no graph TV (*λ*_TV_ = 0*, λ*_1*,W*_ = 0.5), no *W* sparsity (*λ*_TV_ = 1.0*, λ*_1*,W*_ = 0) and removal of both terms (*λ*_TV_ = 0*, λ*_1*,W*_ = 0). The remaining optimization and constraint settings were unchanged. Performance was evaluated using the best matched Pearson correlation with the true programs, boundary AUPRC and blur ratio.

For the orthogonality design ablation, full SORT used the Stiefel constraint and its symmetric NMF and orthogonal Procrustes initialization. The comparison retained unit Euclidean norms but removed pairwise orthogonality. It used standard NMF with a random nonnegative start and inverse rescaling of *W* to preserve the initial reconstruction. The transcriptomic profile update in Stage 1 used residual least squares followed by independent column normalization. In stage 2, each profile gradient was projected onto the tangent space of its unit sphere before independent normalization. The comparison therefore evaluated the orthogonality constraint together with its associated initialization.

Inputs and all remaining settings were matched, including 5,000 highly variable genes, within-section graphs with eight neighbors, rank *K* = 24, *λ*_TV_ = 1.0, *λ*_1*,W*_ = 0.5, *λ*_1*,Q*_ = 300, *λ_−,Q_* = 1.0 and *ε*_TV_ = 0.01. Both variants used graph TV updates every five epochs during stage 2, 40 profile optimizer steps per epoch and 100 epochs per stage. We recorded recovery, boundary metrics, relative reconstruction error, mean absolute cosine similarity between distinct signal profiles and duplicate best matches. Comparisons were paired by seed. Exact two-sided Wilcoxon signed rank tests for class-specific recovery were exploratory, with Benjamini–Hochberg adjustment across the three classes.

### PDAC analyses

We analyzed 108 Visium sections from treatment-näıve primary tumors of 39 patients in the published PDAC atlas [4]. The original study described a continuum of classical and basal-like tumor states with accompanying differences in their surrounding microenvironments. Patients could contribute multiple sections, and sections differed in their enrichment for the published basal-like and classical signatures. The atlas therefore provided a setting for evaluating the recovery of shared tumor-state GEPs and their subsequent spatial and clinical interpretation.

SORT used expression normalized to 10^4^ counts per retained location followed by log(1 + *x*) transformation. Spatial graphs were constructed separately within each section using *k* = 24 nearest neighbors. SORT was fitted with *λ*_TV_ = 0.3, *λ*_1*,W*_ = 0.3 and *λ*_1*,Q*_ = 300.

### Subtype GEP recovery benchmark

To benchmark recovery of the basal-like and classical programs, UCell scores were first computed from signatures reported by the original PDAC atlas [4, 15] to serve as continuous reference signals. Atlas-wide 95th-percentile thresholds were applied to these scores to define high-scoring spatial locations for each subtype. Sections containing at least 10% high-scoring locations were assigned to basal-enriched, classical-enriched or mixed reference strata before decomposition. To ensure that the basal-like reference could be evaluated in every subset, one of the two sections (samples 42 and 43) from patient C2 D6 was included as a fixed basal anchor. Supplementary Fig. S27 shows the atlas-wide reference score distributions, fractions of high-scoring locations and spatial overlays used to assess the available anchor and comparison pools before model fitting.

A total of 20 stratified benchmark subsets were constructed, each containing four basal-enriched, four classical-enriched, and two mixed sections. The two sections from patient C2 D6 were each allocated to 10 subsets under a balanced, randomly ordered anchor schedule. The remaining sections were selected to meet the fixed subtype quotas by sampling eligible non-anchor patients without replacement. When a patient had multiple eligible sections, one was selected at random. Each subset thus comprised ten sections from ten distinct patients.

#### Subset-specific preprocessing and fitting

Genes detected in at least max(10, 0.01*N*_subset_) locations were eligible, where *N*_subset_ is the number of locations in that subset. SORT and NMF used 3,000 highly variable genes selected independently within each subset using the cell ranger procedure, whereas the two STAMP configurations used 6,000 genes and their method-specific count inputs. All four methods were applied at rank *K* = 20. Supplementary Note 3 and Supplementary Tables 3–5 provide the complete sampling procedure and method-specific inputs.

To assess agreement with the published subtype references, we selected, for each subtype reference, the inferred component with the highest correlation. For each inferred component *k* and subtype *t*, let *w_k_* denote its activity score vector and *u_t_*the corresponding UCell reference score vector. The matching was defined as

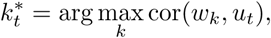

with recovery score cor(*w_k^*^t_, u_t_*). SORT was compared against the best-performing baseline using paired two-sided Wilcoxon signed-rank tests across the 20 subsets (Supplementary Note 3).

### Analysis of the complete PDAC atlas

For this analysis, we retained locations with at least 200 detected genes. Genes detected in at least 1% of retained locations and with a raw mean expression of at least 0.01 were considered for feature selection. A total of 10,000 highly variable genes were selected with the batch-aware seurat v3 procedure. SORT was then fitted to 352,102 retained locations with rank *K* = 40, comprising one background and 39 signal components.

#### Quantifying gene and signature associations with recovered GEPs

We used the published PDAC and tumor microenvironment signatures provided with the original atlas [4]. Gene symbols were standardized, and only genes among the 10,000 used in the fit were retained. Signatures with at least three such genes were included in the analysis. To quantify gene associations with the fitted activity maps, we fitted one multivariable ordinary least-squares (OLS) model for each gene. For gene *g*, normalized expression *X_·g_* was the response, and the 39 signal activity columns in *W*_sig_, together with an intercept, were used as predictors. Coefficient *B_gk_* therefore measures the association of gene *g* with activity map *k* after accounting for the other fitted activities. For the gene-level display, 74 nonredundant representative genes were selected from the published signatures, and their coefficients were z-scored across the 39 signal activity maps and clipped to [−2, 2]. For signature *m* and the signal activity map *k*, the projection score was the signed mean of *B_gk_*across all overlapping genes in this signature, followed by z-scoring across the GEPs included in each display. Supplementary Note 4 records the source fields, symbol harmonization, fixed representative-gene panel and projection calculations.

#### Spatial colocalization analysis

For each displayed section and each pair of GEPs, let *A* and *B* denote the locations with activity scores at or above their respective section-specific 80th percentiles. Let *r* denote the Pearson correlation between the two unthresholded activity vectors across all locations in that section. The descriptive colocalization score was defined as

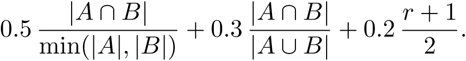

The first term measures containment of the intersection within the smaller set, the second term measures symmetric overlap, and the third captures the covariation. The weighting thus prioritized spatial containment while retaining complementary information from overlap and correlation. The overlap coefficient was omitted when min(|*A*|, |*B*|) = 0, and the correlation term was omitted when either activity score vector had zero variance. The remaining weights were then rescaled to sum to one.

The same section-specific thresholds classified each location as high for the first GEP only, high for the second GEP only, high for both or high for neither. Supplementary Note 4 describes candidate-section selection and the complete calculation.

#### Bulk cohort projection and survival analysis

To relate the recovered programs to patient outcomes, we constructed fixed gene sets from the signal GEP profiles in *Q*. Genes with positive profile values were ranked using a composite of three criteria: the profile weight *q_gk_*, the ratio between the variance of GEP *k*’s fitted contribution and the total fitted variance of gene *g* (*R*_var*,gk*_), and the signed correlation between GEP *k*’s activity and the reconstructed expression of gene *g* (*C*_recon*,gk*_). Let *W_c_* be the activity matrix of the 39 signal GEPs after centering each column, let *w_k_* denote its *k*th column and let *q_g_*contain the profile weights of gene *g*. The fitted expression of gene *g* was *x*^^^*_g_* = *W_c_q_g_* = Σ*_j_ q_gj_w_j_*, and the contribution of GEP *k* was *q_gk_w_k_*. We defined

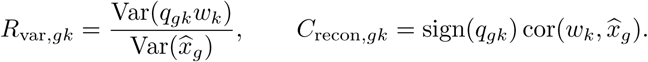

Thus, *R*_var*,gk*_ compares the variance of the individual GEP contribution with the total fitted variance of gene *g*, whereas *C*_recon*,gk*_ measures how closely the fitted expression of gene *g* follows activity *k*, accounting for the sign of its profile weight. Genes with positive profile values were ranked by the sum of their descending ranks across the three criteria. If more than 10 genes were at or above the 90th percentile for all three quantities, ranking was restricted to that set. Otherwise, all genes with positive profile values were ranked. The top 50 genes formed the fixed projection set for each signal GEP. Supplementary Note 4 provides the complete ranking rule.

We used these fixed gene sets to calculate bulk projection scores in the TCGA, CPTAC, QCMG and PanCuRx PDAC cohorts [37–40], which provided both bulk expression and overall survival data. For each GEP and patient, the bulk projection score was calculated by averaging the normalized expression values of genes that overlapped between that GEP’s fixed gene set and the cohort’s expression matrix. These GEP-specific patient-level scores were then z-scored across patients within each cohort. Associations between bulk projection scores and overall survival were estimated with Cox proportional hazards models [41], adjusting for available age, sex and stage covariates. Log hazard ratios and standard errors from each cohort were combined by inverse variance meta-analysis under a fixed-effect model. Meta-analysis *P* values were adjusted across the 39 signal GEPs using the Benjamini–Hochberg procedure to control the false discovery rate (FDR). Heterogeneity between cohorts was summarized by *I*^2^, the estimated proportion of variation among cohort effects attributable to heterogeneity rather than sampling error.

For Kaplan–Meier visualization, we displayed the four GEPs with the smallest FDR values after adjustment across all 39 GEPs. The curves pooled the four cohorts after standardizing projection scores within each cohort and used the global median as the cutoff. High was defined strictly above the median (High *n* = 195, Low *n* = 196, 391 patients and 260 events). The Kaplan–Meier curves complemented the continuous Cox models and meta-analysis, and their log-rank *P* values were interpreted descriptively. Supplementary Note 4 provides the complete display procedure. Cohort and event counts are provided in Supplementary Table 1.

### Cross-species cerebellar analysis

The published Stereo-seq atlas maps the cerebellar cortex of macaque, marmoset and mouse and provides layer and cortical-depth information for cross-species comparison [5]. We selected three adjacent sections from one animal of each species, yielding nine sections and 1,546,637 spatial locations in total. This subset was used to compare the spatial activity distribution and activity magnitude of fitted GEPs among the selected specimens. Gene symbols were converted to uppercase, and exact symbol matches present in all nine sections were retained without orthology mapping. Each section was normalized to a total of 10^4^ counts per location and log-transformed. SORT was fitted with 4,000 highly variable genes at rank *K* = 25.

#### Cortical depth profiling and activity magnitude comparison

The published atlas provided a cortical depth coordinate that placed the Purkinje cell layer at zero. Negative values extended toward the granule layer, and positive values toward the molecular layer and surface. To compare spatial activities across sections and species, we divided the interval between the 0.5th and 99.5th percentiles of the pooled cortical depth coordinates into 160 equally spaced bins. For each section and GEP, bins containing fewer than 20 locations were treated as missing. The remaining profile was smoothed using a moving average over nine bins, with each bin weighted by its number of locations. Missing bins received zero weight, and windows at either end were shortened. We then averaged the three section-level spatial activity curves for each species, weighting each bin by its total number of locations across sections. Each resulting spatial activity profile was z-scored across bins with available values. Spatial activity similarity was summarized by the mean of the three Pearson correlations from pairwise comparisons among species.

To compare activity magnitude, we returned to the unstandardized, smoothed section activity profiles, and computed their arithmetic means over the common cortical depth interval [−1, 1]. All sections contained the same 119 finite bin centers in this interval. Section-level mean activities were then averaged within species. Relative variation in magnitude was summarized by the coefficient of variation (CV) of these three species means:

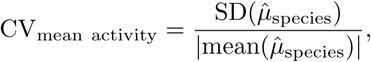

where *µ*^_species_ is the species mean of the three section-level mean activities and SD denotes the population standard deviation across the three species means.

#### Validation of GEP activity using prioritized gene detection

For GEP02 and GEP14, a fixed pool of 60 positive-weight genes was defined using transcriptomic profile weight, the component-to-total fitted variance ratio (*R*_var*,gk*_) and the reconstructed expression–activity correlation (*C*_recon*,gk*_). The 20 genes with the largest profile weights within each GEP-specific pool were retained.

To assess whether the activity differences for GEP02 and GEP14 were accompanied by concordant gene-detection differences, we examined their prioritized genes within corresponding spatial domains in both the fitting and additional animal sets. The analysis used 18 sections: nine from the original SORT fit and nine in three adjacent sections from one additional animal of each species. Standardized gene symbols were intersected across all sections to define a common normalization and projection universe. This feature-matching procedure aligned measured genes across sections without establishing one-to-one orthology.

In the fitting animals, the target domain for each GEP comprised locations at or above the 90th percentile of its strictly positive learned activity values within each section. For each additional animal, we held *Q* fixed and masked the 20 tested genes before projection. For location *i*, let *x_i_*denote its normalized expression vector and *q^(−G)^_k_* the masked profile of GEP *k*. The projection score was defined as the nonnegative least-squares coefficient

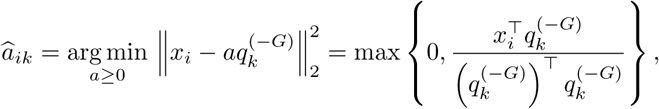

which locates corresponding spatial domains without using the tested genes. In the additional animals, the corresponding domain comprised locations at or above the 90th percentile of strictly positive projection scores within each section. Detection fractions for the masked genes were calculated as the proportion of locations with nonzero expression within each domain, averaged across the three sections from each animal, and contrasted as marmoset minus the mean of macaque and mouse.

To assess the robustness of these results, we compared the detection fractions observed for the 20 selected genes against 10,000 control gene sets matched for detection prevalence and expression level. Calibration was performed separately for the animals used for fitting and the additional animals. One-to-one orthology was then evaluated through the Ensembl release 116 REST homology endpoint and used only as a sensitivity filter [42, 43]. A separate control comparison contrasted 12 genes from background GEP00 with 12 from signal GEP01 within their shared granule layer domain in the original sections. Supplementary Note 5 provides detailed descriptions of the support thresholds, matched-set construction, orthology criteria, and reciprocal-contrast procedure.

#### Computational scalability benchmark

We benchmarked six cross-species input sizes, ranging from 51,997 to 1,546,637 spatial locations, with 4,000 highly variable genes. The rank *K* = 25, model settings, software environment and NVIDIA A30 graphics processing unit (GPU) were held fixed. Fractional inputs were centered, spatially contiguous rectangular crops of one representative section per species. Larger inputs used complete sections. We recorded end-to-end and model-fitting times. Each input size was run once, the benchmark thus describes the empirical runtime over the tested range.

### Mouse embryogenesis analysis

The mouse organogenesis Stereo-seq atlas maps transcriptional and anatomical changes across embryonic development [6]. We analyzed eight cross-sectional stages from E9.5 to E16.5 and used the anatomical annotations provided with the atlas to interpret the GEPs. The cross-sectional series enabled comparison of temporal deployment across stages and graded activity within developing anatomical compartments. Locations with at least 50 detected genes were retained, and genes detected in at least 0.1% of locations were kept. Counts were normalized to 10^4^ per location and log-transformed. The final joint analysis included 520,389 spatial locations and 8,000 genes, with retained stage sizes ranging from 5,902 to 121,752 locations. SORT was fitted jointly at rank *K* = 60, comprising one background component and 59 signal GEPs (Supplementary Tables 3–4).

#### Temporal and anatomical activity analyses

For each signal GEP, mean spatial activity score was calculated across all retained locations within each stage and divided by that GEP’s maximum stage mean for display. Profiles were arranged from early peaks through intermediate maxima to later increases for visualization.

To characterize the spatial deployment of selected GEPs, we summarized their activities within specific anatomical annotations. GEP25 activity was assessed in the sclerotome, head mesenchyme, cartilage primordium, and cartilage; GEP31 activity was assessed in the surface ectoderm, epidermis, and mucosal epithelium; and GEP02 activity was assessed in the brain. Stage-by-annotation strata with fewer than 50 locations were excluded from the analysis. To interpret the biological functions of these GEPs, we compiled literature-informed gene sets containing genes with established roles in the corresponding tissue or developmental process. Supplementary Table 2 records their membership, biological rationale, and literature sources. For each stage and anatomical annotation, we computed a gene-set expression score by z-scoring log-transformed expression for each gene and averaging across the gene set. Spearman correlation between GEP activity score and this expression score measured their association.

To assess sensitivity to variation in detected-gene coverage, spatial activity, expression score and detected-gene count were rank transformed. Ranked activity and expression score were separately regressed on ranked detected-gene count. The Pearson correlation between the residuals was reported as the coverage-adjusted rank correlation, where coverage denotes the number of detected genes per location. Supplementary Note 6 provides the temporal display ordering, evaluated contexts and GEP02 sensitivity analyses.

#### GEP06 and GEP43 profile similarity and spatial overlap

The positive part of transcriptomic profile *k* was defined as *Q^+^_k_ = max(Q_k_, 0)*. GEP06 and GEP43 profile similarity was summarized by the cosine similarity between *Q^+^_06_* and *Q^+^_43_*. We also calculated the Jaccard index of their top 50 genes with positive weight in *Q*.

For spatial display, activity score was scaled as *W_k_* = min{*W_k_/q*_0.99_(*W_k_*), 1}, where the 99th percentile was calculated across all retained locations. Colocalization scores and high-activity Jaccard indices were calculated as described for PDAC, using the 90th-percentile unscaled activity threshold for each GEP within each stage. Gene-set concordance was calculated separately within each stage across all locations, comparing each GEP’s activity with its own fixed gene-set score using the coverage-adjustment procedure above. Supplementary Note 6 gives the calculation details.

### Functional enrichment of genes with positive profile weights

For the PDAC and mouse embryogenesis analyses, genes were ranked within each signal GEP by the transcriptomic profile weights in *Q*. The primary gene set comprised genes with positive weights in the top 2% of the complete ranking for that GEP. This gave 200 genes in PDAC and 160 genes in mouse embryogenesis. Top 1% and top 5% sets were additionally evaluated as sensitivity analyses to assess whether the pathway enrichment results were robust to the choice of gene-set size cutoff. The enrichment universe for each dataset was defined as all genes included in the corresponding SORT fit.

We performed one-sided over-representation analysis (ORA) using Hallmark and Reactome gene sets from MSigDB v7.5.1 for PDAC. For mouse embryogenesis, we used Gene Ontology Biological Process terms from org.Mm.eg.db v3.18.0. Gene sets containing 10–500 genes from the enrichment universe were tested. Hypergeometric *P* values were adjusted by the Benjamini–Hochberg method across the complete family of signal-GEP–pathway tests, separately for each dataset, collection and profile-size cutoff. A pathway was considered eligible for display when its primary top 2% result had global FDR *<* 0.05 and the top 1% or top 5% analysis supported the same pathway. Displayed pathways were nonredundant eligible terms relevant to the GEPs examined in the Results.

ORA quantified pathway over-representation among strongly positive genes in each fitted transcriptomic profile and used *Q* only. The PDAC gene and signature annotations instead used per-gene regression coefficients obtained with fitted spatial activities *W* as predictors, followed by signature projections calculated from those coefficients. The enrichment analysis therefore provides transcriptomic profile-level functional context rather than a test of pathway activation, differential expression or cell identity. Supplementary Note 7 gives the complete gene-set construction, multiple-testing family and display criteria.

### Competing methods

In the simulation benchmark, SORT was compared with PCA [7], nonnegative matrix factorization (NMF) [8], GraphPCA [10] (https://github.com/YANG-ERA/GraphPCA-Turbo), NNMF [12] (https://github.com/ragnhildlaursen/NNMF) and STAMP [14] (https://github.com/JinmiaoChenLab/scTM). PCA and NMF were run using scikit-learn [44].

In the PDAC benchmark, SORT was compared with NMF and two STAMP configurations. STAMP-noCov omitted sample covariates, whereas STAMP included patient identity as a covariate.

The comparison methods used the same datasets or selected sections and the same nominal rank where applicable. Official implementations, preprocessing, graph settings and covariates are documented in Supplementary Tables 3–5. Exact method-specific configurations and random seeds are documented in the analysis code.

## Computational implementation

SORT was implemented in Python 3.8.20 with NumPy 1.22.4 [45], SciPy 1.10.1 [46], scikit-learn 1.3.2 [44], PyTorch 2.4.1 [47] and CuPy 12.3.0 [48]. PyTorch implemented Stiefel updates and CuPy accelerated sparse graph operations.

## Statistics and reproducibility

Paired two-sided Wilcoxon signed-rank tests compared methods across matched simulation replicates and PDAC resampling iterations. Survival analyses used covariate-adjusted Cox models, fixed-effect inverse variance meta-analysis and Benjamini–Hochberg correction. Kaplan–Meier log-rank tests were descriptive. Matched-set upper-tail probabilities for GEP02 and GEP14 were calculated from 10,000 expression- and detection-matched control sets and were not treated as population-level species tests.

Unless an inferential test is stated, correlations, coefficients of variation, temporal summaries, gene-set associations, sensitivity analyses and computational benchmarks are descriptive. A simulation replicate was one generated dataset. A PDAC iteration was one benchmark subset containing one section per patient. Locations within a section were not treated as independent biological replicates. PDAC sections were nested within patients, and cerebellar sections were nested within animals.

Simulation recovery used 20 datasets, sensitivity and ablation analyses used 10 matched seeds, and the PDAC subtype benchmark used 20 paired iterations. No statistical method was used to predetermine sample size. Public atlas sizes and the prespecified simulation and resampling designs determined the analyzed sample sizes. Supplementary Table 7 summarizes analysis units and sample accounting.

## Data availability

All datasets analyzed in this study are available through the repositories associated with their original studies, subject to the corresponding access conditions. The PDAC Visium dataset is available from the Gene Expression Omnibus under accession GSE282302 (https://www.ncbi.nlm.nih.gov/geo/query/acc.cgi?acc=GSE282302). Mouse whole-embryo Stereo-seq data across developmental time points are available from the MOSTA database (https://db.cngb.org/stomics/mosta/) [6]. The cross-species cerebellar Stereo-seq dataset is available through the CNGBdb STOmics database [5] (http://db.cngb.org/stomics/cbmsta). Processed TCGA, CPTAC, QCMG and PanCuRx bulk expression and clinical data were obtained from the Zenodo companion repository of Liu et al. [30, 49] (https://doi.org/10.5281/zenodo.8271036). The corresponding original cohort studies are TCGA [37], CPTAC [38], QCMG [39] and PanCuRx [40]. Access and redistribution remain subject to the source repositories and cohort restrictions.

## Code availability

The SORT software package is available at https://github.com/zerovain/SORT under the MIT license, with documentation and tutorials at https://sort-tutorial.readthedocs.io.

## Author contributions

Conceptualization: SZ and LZ; methodology: LZ; software: LZ; validation: LZ; formal analysis: LZ; investigation: LZ; data curation: LZ; writing—original draft preparation: LZ; writing—review and editing: all authors; visualization: LZ; supervision: LL, SZ; project administration: SZ; funding acquisition: SZ. All authors have read and agreed to the published version of the manuscript.

