## Supplementary materials for "SORT disentangles multilevel gene programs across heterogeneous spatial transcriptomic atlases"

### Supplementary Information for SORT

#### Supplementary Note 1. SORT objective and optimization details

For spatial section $s = 1, \ldots, S$, SORT takes a normalized expression matrix $X^{(s)} \in \mathbb{R}^{n_s \times p}$, where rows are spatial locations and columns are genes. The model fits

$$X^{(s)} \approx W^{(s)} Q^\top,$$

where

$$Q = (q_0, Q_{\text{sig}}) \in \mathbb{R}^{p \times K}, \qquad W^{(s)} = (w_0^{(s)}, W_{\text{sig}}^{(s)}) \in \mathbb{R}_+^{n_s \times K}.$$

The pair $(w_0^{(s)}, q_0)$ is the background component, and the remaining columns form the signal profiles and their section activities:

$$Q_{\text{sig}} \in \mathbb{R}^{p \times K_{\text{sig}}}, \qquad K_{\text{sig}} = K - 1.$$

The Stiefel constraint is imposed on $Q_{\text{sig}}$, not on the full $Q$ matrix:

$$Q_{\text{sig}}^\top Q_{\text{sig}} = I_{K_{\text{sig}}}.$$

The constrained objective used in the reported analyses can be written as

$$\begin{aligned} \min_{\{W^{(s)}\}, Q} \quad & \sum_{s=1}^{S} \left\| X^{(s)} - W^{(s)} Q^\top \right\|_F^2 + \lambda_{\text{TV}} \sum_{s=1}^{S} \text{TV}_{\mathcal{G}^{(s)}}(W^{(s)}) + \mathcal{R}(\{W^{(s)}\}, Q) \\ \text{s.t.} \quad & W^{(s)} \geq 0, \qquad Q_{\text{sig}}^\top Q_{\text{sig}} = I_{K_{\text{sig}}}. \end{aligned}$$

The auxiliary regularizer $\mathcal{R}(\cdot, \cdot)$ contains $W$ sparsity, $Q$ sparsity and negative profile shrinkage:

$$\mathcal{R}(\{W^{(s)}\}, Q) = \lambda_{1,W} \sum_s \|W^{(s)}\|_1 + \lambda_{1,Q} \|Q_{\text{sig}}\|_1 + \lambda_{-,Q} \sum_{g,k} \{\max(0, -Q_{\text{sig},gk})\}^2,$$

Dataset-specific values of $\lambda_{1,W}$ and $\lambda_{1,Q}$ are listed in Supplementary Table 7. The objective notation $\lambda_{\text{TV}}$ corresponds to the implementation parameter $\alpha$.

The reported $\lambda_{1,W}$ and $\lambda_{1,Q}$ values are base sparsity weights. Component-specific $W$ weights were fixed after initialization by scaling the base value in proportion to the initial column $L_2$ norm. Signal-profile weights used the same scaling when the ratio between the largest and smallest initial signal-profile norms exceeded 1.5. Otherwise, all signal profiles used the base $\lambda_{1,Q}$ value. The background profile was not subject to $Q$-sparsity regularization. Adaptive signal-profile weights were used only for the cerebellum analysis.

For each spatial section, graph total variation (TV) regularization uses an edge-incidence operator $D_{\mathcal{G}}^{(s)}$, which maps spatial activities to edge-wise activity differences. Section-wise operators may be assembled block-diagonally for computation, so no graph edge connects different tissue sections.

The following formulas describe the active optimized implementation used for all reported analyses. The separable graph TV penalty is

$$\mathrm{TV}_{\mathcal{G}}(W) = \sum_{(i,j)\in E} a_{ij} \sum_{k=1}^{K} \sqrt{(W_{ik} - W_{jk})^2 + \varepsilon_{\mathrm{TV}}}.$$

Thus, with the reported default $\varepsilon_{\mathrm{TV}} = 0.01$, the active expression is $\sqrt{(W_{ik} - W_{jk})^2 + 0.01}$. Reported analyses first constructed a symmetric binary k-nearest-neighbor adjacency matrix $A$ within each section. The normalized graph Laplacian $I - D^{-1/2}AD^{-1/2}$ was then used to obtain static TV edge weights proportional to $A_{ij}/\sqrt{d_i d_j}$, where $d_i$ is the graph degree. After section-wise graphs were assembled, the extracted weights were divided by one maximum calculated over the combined block-diagonal graph supplied to the fit. Thus, $a_{ij}$ reflects degree-normalized connectivity rather than physical distance; physical distances were not used as TV weights in the reported analyses. The sum extends over all fitted activity columns. The background $W$ column was TV-regularized together with the signal columns; its exclusion applied only to the Stiefel constraint on $Q_{\mathrm{sig}}$.

**IRLS approximation for graph TV.** For edge $e = (i, j)$ and component $k$, let $d_{e,k} = W_{ik} - W_{jk}$ denote the activity difference, and write $a_e = a_{ij}$ for the static edge weight. At iteration $t$, concavity of the square root gives the quadratic majorization

$$\sqrt{d_{e,k}^2 + \varepsilon_{\mathrm{TV}}} \leq \frac{d_{e,k}^2 + \varepsilon_{\mathrm{TV}}}{2\sqrt{(d_{e,k}^{(t)})^2 + \varepsilon_{\mathrm{TV}}}} + \frac{\sqrt{(d_{e,k}^{(t)})^2 + \varepsilon_{\mathrm{TV}}}}{2}.$$

After removing terms independent of the updated $W$, the weighted quadratic contribution is proportional to

$$\frac{a_e}{2\sqrt{(d_{e,k}^{(t)})^2 + \varepsilon_{\mathrm{TV}}}} d_{e,k}^2.$$

The factor 1/2 is absorbed into the global graph TV weight, giving

$$\omega_{e,k}^{(t)} = \frac{a_e}{\sqrt{(d_{e,k}^{(t)})^2 + \varepsilon_{\mathrm{TV}}}}.$$

Writing $E$ for the edge-incidence matrix, the corresponding reweighted term is

$$E^\top \left(\omega^{(t)} \odot EW\right).$$

This weighting penalizes small local activity differences more strongly than large jumps. The bound is tight at the current iterate, and the resulting surrogate term is incorporated into the multiplicative $W$ updates within the alternating non-convex optimization.

**Alternating optimization.** Optimization proceeds by updating the section activity matrices $W^{(s)}$ and the shared profile matrix $Q = (q_0, Q_{\mathrm{sig}})$ in turn. Within the profile update, $q_0$ is estimated by unconstrained least squares, whereas $Q_{\mathrm{sig}}$ is updated in two stages. Automatic initialization used symmetric nonnegative matrix factorization (NMF) to obtain nonnegative activities and singular value decomposition to initialize the shared profiles orthogonally.

With $Q$ fixed, each activity matrix solves

$$\min_{W^{(s)}\geq 0} \left\| X^{(s)} - W^{(s)}Q^\top \right\|_F^2 + \lambda_{\mathrm{TV}}\, \mathrm{TV}_{\mathcal{G}^{(s)}}(W^{(s)}) + \lambda_{1,W}\|W^{(s)}\|_1. \tag{1}$$

The nonnegativity of $W^{(s)}$ is maintained by multiplicative updates. Graph TV was inactive in stage 1 and applied in stage 2. The graph TV term is replaced at the current iterate by the IRLS quadratic majorizer derived above. For implementation, if $W_{\text{diff}} = EW$, the dynamic edge weights are

$$\widetilde{w}_{e,k} = \frac{a_e}{\sqrt{(W_{\text{diff},e,k})^2 + \varepsilon_{\text{TV}}}}, \tag{2}$$

and the reweighted graph contribution is evaluated from

$$L_{\text{TV}} W = E^\top (\widetilde{w} \odot EW), \qquad D_{\text{TV}} = |E|^\top \widetilde{w}. \tag{3}$$

With the activity matrices and $Q_{\text{sig}}$ fixed, the background profile solves

$$\min_{q_0} \sum_s \left\| X^{(s)} - w_0^{(s)} q_0^\top - W_{\text{sig}}^{(s)} Q_{\text{sig}}^\top \right\|_F^2, \tag{4}$$

without an orthogonality or sign constraint.

For the signal profile updates, stack sections by rows and define $R_0 = X - w_0 q_0^\top$. Stage 1 omits the signal profile sparsity and negative value penalties and solves

$$\min_{Q_{\text{sig}}^\top Q_{\text{sig}} = I} \left\| R_0 - W_{\text{sig}} Q_{\text{sig}}^\top \right\|_F^2. \tag{5}$$

If $R_0^\top W_{\text{sig}} = U \Sigma V^\top$, its orthogonal Procrustes solution is $Q_{\text{sig}} = UV^\top$. Stage 2 instead solves

$$\min_{Q_{\text{sig}}^\top Q_{\text{sig}} = I} \left\| R_0 - W_{\text{sig}} Q_{\text{sig}}^\top \right\|_F^2 + \lambda_{1,Q} \|Q_{\text{sig}}\|_1 + \lambda_{-,Q} \|Q_{\text{sig},-}\|_F^2. \tag{6}$$

For a Euclidean gradient $G$, Riemannian Adam uses the tangent projection

$$\Pi_Q(G) = G - Q\ \text{sym}(Q^\top G), \qquad \text{sym}(M) = \tfrac{1}{2}(M + M^\top), \tag{7}$$

followed by the polar retraction

$$\text{Retr}_Q(\Delta) = (Q + \Delta) \left[ (Q + \Delta)^\top (Q + \Delta) \right]^{-1/2}. \tag{8}$$

This keeps $Q_{\text{sig}}^\top Q_{\text{sig}} = I$ after every signal profile update.

**Supplementary Algorithm 1** Two-stage alternating optimization in SORT

**Require:** Expression matrices $\{X^{(s)}\}$, spatial graphs $\{\mathcal{G}^{(s)}\}$, rank $K$, regularization parameters and stage-specific epoch numbers $E_1, E_2$

Initialize $W^{(s)} \geq 0$, $q_0$ and orthonormal $Q_{\text{sig}}$
**for** stage $h = 1, 2$ **do**
  **for** epoch $e = 1, \ldots, E_h$ **do**
    **for** section $s = 1, \ldots, S$ **do**
      Update $W^{(s)}$ using equation (1), setting $\lambda_{\text{TV}} = 0$ in stage 1 and evaluating its graph TV term with equations (2)–(3) in stage 2
    **end for**
    Update $q_0$ using equation (4) according to the stage schedule
    **if** $h = 1$ and a signal profile update is scheduled **then**
      Update $Q_{\text{sig}}$ using the orthogonal Procrustes solution to equation (5)
    **else if** $h = 2$ **then**
      Optimize equation (6) using the tangent projection and retraction in equations (7) and (8)
    **end if**
  **end for**
**end for**
**return** $\{W^{(s)}\}$, $q_0$ and $Q_{\text{sig}}$

Supplementary Algorithm 1 summarizes the complete update schedule and links each step to its corresponding optimization subproblem.

#### Supplementary Note 2. Simulation benchmark, sensitivity analysis and metrics

**Simulation design.** The simulation benchmark adapted a Splat/binomial-thinning design from single-cell measurements to spatial locations. Each replicate contained eight spatial sections, each represented by a $36 \times 36$ grid, for $N = 10{,}368$ spatial locations and $J = 8{,}000$ genes before analysis filtering. A baseline count matrix was generated with `splatter::splatSimulate`, with Splat parameters fitted to a reference PDAC dataset. The Splat-generated baseline counts provided library-size variation, gene-level variability and background expression structure before structured spatial programs were injected.

Structured signals were injected with `seqgendiff::thin_diff` [1] using a spot-by-program design matrix $L_{\text{true}}$ and a gene-by-program coefficient matrix $F_{\text{true}}$. In the manuscript notation, these matrices are denoted by $W_{\text{true}} = L_{\text{true}}$ and $Q_{\text{true}} = F_{\text{true}}$, respectively, and $W_{\text{true}} Q_{\text{true}}^{\top}$ specifies the location-by-gene signal supplied to the thinning procedure. The structured ground truth contained no separate background component: baseline and background expression variation was already represented by the Splat-generated reference-derived count matrix, whereas $W_{\text{true}}$ and $Q_{\text{true}}$ encoded only the 13 injected non-background GEPs used for recovery evaluation.

The 13 structured programs comprised eight patient-specific programs, three shared subtype-related programs and two non-focus region-related programs. Patient-specific, subtype-related and non-focus region-related programs used 500, 100 and 75 non-overlapping marker genes, respectively. Mean $\log_2$ fold changes were approximately 1.585, 1.700 and 1.585 for the three program classes. All true signatures and marker perturbations were nonnegative, matching the assumptions of the

NMF-based baselines. Spatial layouts used geometrically defined focus and non-focus patterns, including wedge-sector and annular layouts. Counts were normalized with scran size factors using `quickCluster` and `calculateSumFactors`, then log-transformed. Let $\widetilde{x}_{ij}$ denote the simulated count after the structured perturbation, $\hat{s}_i$ denote the scran size factor for spatial location $i$, and $m_{\hat{s}}$ denote the median size factor. The normalized expression value used for model fitting was

$$y_{ij} = \log\left(10\left(\frac{m_{\hat{s}}\widetilde{x}_{ij}}{\hat{s}_i} + 0.1\right)\right).$$

All six comparison methods were fitted at the same nominal rank of 24 where applicable. For SORT, this comprised one background component and 23 signal components, whereas the recovery target contained 13 injected non-background programs. We treated this as an over-specified benchmark setting, which allowed residual components, signal splitting and redundancy to remain observable.

**GEP recovery (PCC).** For true program $k$, let $\ell_k \in \mathbb{R}^N$ denote its column of $W_{\text{true}}$, and let $\widehat{w}_c$ denote inferred component $c$. After independently min–max scaling the ground-truth and inferred activity vectors across all simulated spatial locations pooled across sections, the recovery of program $k$ was

$$R_k = \max_{c=1,\ldots,K} \text{cor}_{\text{Pearson}}(\ell_k, \widehat{w}_c).$$

Correlations were signed; absolute-value matching was not used. Matching was performed independently for each true program, so an inferred component could match more than one true program. Overall recovery for a replicate was the arithmetic mean of $R_k$ across the 13 structured programs.

**Boundary AUPRC and blur ratio.** Boundary metrics used undirected horizontal and vertical grid edges within each section; no edges connected different sections. For an edge $e = (i, j)$, ground-truth and inferred edge-gradient magnitudes were

$$g_{e,k} = |\ell_{ik} - \ell_{jk}|, \qquad \widehat{g}_{e,k} = |\widehat{w}_{i,c(k)} - \widehat{w}_{j,c(k)}|,$$

where $c(k)$ is the component selected by $R_k$, and both activity vectors were min–max normalized before edge gradients were calculated. The true-boundary set $\mathcal{B}_k$ contained edges at or above the 90th percentile of nonzero $g_{e,k}$. Boundary AUPRC was the average precision obtained by using $\widehat{g}_{e,k}$ to classify $\mathbf{1}\{e \in \mathcal{B}_k\}$.

Let $\mathcal{N}_k$ contain non-boundary edges incident to at least one node belonging to an edge in $\mathcal{B}_k$. The blur ratio was

$$\text{Blur}_k = \frac{\sum_{e\in\mathcal{N}_k} \widehat{g}_{e,k}}{\max\left(\sum_{e\in\mathcal{B}_k} \widehat{g}_{e,k}, 10^{-12}\right)}.$$

Lower values indicate less edge-gradient spillover immediately adjacent to true boundaries. Boundary AUPRC and blur ratio were averaged across the 13 structured programs within each replicate.

**Spatial autocorrelation.** For the selected, min–max-normalized activity vector $\widehat{w}$, let $z_i = \widehat{w}_i - \overline{\widehat{w}}$. With $E$ denoting the within-section undirected grid edges, edge-based Moran's $I$ was

$$I = \frac{N}{2|E|} \frac{2\sum_{(i,j)\in E} z_i z_j}{\sum_i z_i^2}.$$

Moran's $I$ was computed per structured program and averaged within replicate. It was interpreted as spatial coherence rather than boundary accuracy and was considered jointly with boundary AUPRC and blur ratio.

**Parameter sensitivity and ablation.** All sensitivity and ablation analyses used the same 10 matched simulation seeds. Supplementary Table 9 summarizes the tested parameter values and the number of simulation datasets used for each analysis.

**Orthogonality design ablation.** We compared full SORT with a nonorthogonal variant. Full SORT constrained the signal profiles to the Stiefel manifold and used symmetric NMF and orthogonal Procrustes initialization. The comparison retained a unit Euclidean norm for each profile but left pairwise inner products unrestricted. Its initialization used standard NMF with a random nonnegative start, coordinate descent and Frobenius loss. The profiles were normalized independently, with inverse rescaling of the corresponding $W$ columns to preserve reconstruction.

In stage 1, residual least squares was followed by independent column normalization. In stage 2, each profile gradient was projected onto the tangent space of its unit sphere before independent normalization. All other settings matched full SORT. Shared settings are summarized in Supplementary Tables 4 and 6.

Recovery and boundary metrics followed the definitions above. Relative reconstruction error was the residual Frobenius norm divided by that of the expression matrix. Mean absolute cosine similarity among signal profiles verified the fitted geometry. Duplicate assignments counted ground truth programs whose independently best matched component had already been selected. Comparisons were paired by seed. Exact two-sided Wilcoxon signed rank tests were exploratory, with Benjamini–Hochberg adjustment across the patient, subtype and region recovery comparisons.

### Supplementary Note 3. PDAC subtype-reference benchmark

The PDAC subtype-reference benchmark used 20 controlled resampling iterations stratified by reference subtype, with one section per patient. Basal and classical UCell scores were computed from subtype signature genes reported in the original PDAC atlas. These scores served as biological reference signals for method comparison. Atlas-wide 95th-percentile (q95) thresholds were calculated separately for the basal and classical scores. A section was considered basal-positive or classical-positive if at least 10% of in-tissue spots exceeded the corresponding q95 threshold. Basal-only positive sections were labeled basal-enriched, classical-only positive sections were labeled classical-enriched, and sections positive for both basal and classical were labeled mixed. These eligibility strata were defined entirely from reference UCell scores before fitting any compared method.

**Reference diagnostics.** Raw section-average UCell scores were used to assess reference coverage within each signature. Atlas-wide diagnostics identified samples 42 and 43 from patient C2_D6 as the two strongest basal-reference sections and confirmed spatially coherent basal-high regions (Supplementary Fig. S27a–c). The classical-enriched pool contained 12 sections from eight patients. These diagnostics defined benchmark subsets with evaluable subtype reference signals and were not used during model fitting.

To keep the basal reference axis evaluable in every subset, one of these sections was included as a basal anchor. Each iteration contained four basal-enriched, four classical-enriched and two mixed sections. The two basal anchors were each assigned to 10 iterations. Eligible non-anchor patients were sampled without replacement within these subtype quotas, with one eligible section selected for each patient. Both mixed-stratum patients entered every iteration. Each iteration therefore contained 10 sections from 10 patients, with one section per patient. The 20 iterations were controlled resamples of the same benchmark.

Without additional location filtering, genes detected in at least $\max(10, 0.01N_{\text{subset}})$ subset locations were eligible for feature selection, where $N_{\text{subset}}$ is the number of locations in that subset. Counts were normalized to $10^4$ per location and $\log(1+x)$-transformed for SORT and NMF. Within each subset, 3,000 highly variable genes were selected independently for SORT and NMF using the `cell_ranger` procedure. STAMP-noCov and STAMP instead used their method-specific count inputs with 6,000 selected genes. SORT used a separate $k = 24$ spatial graph within each section and rank $K = 20$, with analysis-specific settings summarized in Supplementary Table 7.

SORT, NMF, STAMP-noCov and STAMP were applied to identical selected sections. STAMP-noCov omitted the patient covariate used by STAMP. For each method and subtype, the matched component was selected using only its Pearson correlation with the target reference score:

$$c_t^* = \arg\max_c \ \mathrm{cor}(w_c, u_t),$$

where $w_c$ is the inferred activity of component $c$ and $u_t$ is the target basal or classical UCell score. Component matching was performed after model fitting and did not affect estimation. Subtype recovery was reported as the Pearson correlation $\mathrm{cor}(w_{c_t^*}, u_t)$ between the matched component and the target subtype UCell score. The comparison method with the highest median recovery correlation was identified separately for the basal and classical references across the 20 paired iterations.

As a separate post-fit specificity diagnostic, we compared the target correlation with correlation to the alternative subtype reference:

$$d_t = \mathrm{cor}(w_{c_t^*}, u_t) - \left|\mathrm{cor}(w_{c_t^*}, u_{\bar{t}})\right|,$$

where $u_{\bar{t}}$ is the alternative basal or classical UCell score. The matched component $c_t^*$ remained fixed from target-only matching. This diagnostic was not used for component selection or model fitting.

#### Supplementary Note 4. Complete PDAC atlas annotation, colocalization and survival

**Complete atlas decomposition.** The full PDAC decomposition used 108 Visium FFPE sections from 39 patients. After the filtering and feature selection described in Methods, SORT was fitted to 352,102 locations and 10,000 genes at rank $K = 40$, with component 0 treated as background and 39 signal components.

**GEP–gene association.** Normalized log-expression values were modeled gene-wise using ordinary least squares with an intercept and the fixed SORT signal activities $W_{\mathrm{sig}}$ as predictors. The $W_{\mathrm{sig}}$ predictors were used on their fitted scale and were not standardized before regression. The resulting coefficient matrix $B$ was used as a post hoc GEP–gene association matrix. The coefficients depend on the fitted activity scale and were used for annotation and visualization, not for causal interpretation or differential-expression testing.

Gene categories and membership for main Fig. 4a and the signatures projected in main Fig. 4b and Supplementary Fig. S9 were obtained from the published PDAC and tumor microenvironment signatures compiled in sheet 4 of Table S1 from the original atlas. Gene symbols were trimmed, uppercased, deduplicated and harmonized. For Fig. 4a, they were intersected with the 10,000 genes used in the SORT fit. To accommodate the available figure width, main Fig. 4a displays 74 nonredundant representative genes from these source signatures. The associated categories describe tumor subtype, epithelial region, stromal, vascular and immune-associated biology and were used only for post-fit annotation. Heatmap values were calculated by z-scoring $B_{gk}$ across all 39 signal GEPs for each gene and were visually clipped to $[-2, 2]$. The heatmap therefore shows relative gene–GEP association patterns across the fitted signal activities.

**Signature projection.** For each signature, we averaged the signed post hoc association coefficients $B_{gk}$ across genes represented in the full gene-level association matrix. The main Fig. 4b heatmap was row-wise standardized across its 14 displayed signal GEPs, whereas Supplementary Fig. S9 was row-wise standardized across all 39 signal GEPs.

**Colocalization.** For each displayed section and pair of GEPs, the 80th percentile was calculated separately for each activity vector within that section. These thresholds defined the sets $A$ and $B$ of locations with high activity. The overlap coefficient was $|A \cap B| / \min(|A|, |B|)$, and the Jaccard index was $|A \cap B|/|A \cup B|$. The colocalization score combined overlap coefficient, Jaccard index and rescaled Pearson correlation as

$$0.5 \times \text{overlap} + 0.3 \times \text{Jaccard} + 0.2 \times \frac{r+1}{2},$$

where $r$ is the Pearson correlation between the two unthresholded activity vectors across all locations in that section. The colocalization scores reported in the Results for main Fig. 4c were calculated from these section-specific 80th-percentile masks. The overlap-coefficient term was excluded when $\min(|A|, |B|) = 0$. The correlation term was excluded when either activity vector had zero variance, after which the remaining weights were rescaled to sum to one. The same masks defined the first-only, second-only, both-high and neither display categories.

Candidate sections were screened using analogous atlas-wide 90th-percentile masks. Retrospective permutation tests assessed the displayed section-level scores by holding the first GEP and section labels fixed and permuting the second GEP's location labels within each section. Empirical $P$ values used $(\#\{\text{null} \geq \text{observed}\} + 1)/(N + 1)$, with $N = 500$ permutations. Supplementary Fig. S11 separately used global 90th-percentile masks to compare section-level activity correlation with co-high Jaccard overlap for GEP16 and GEP32.

**Survival projection.** For bulk survival analysis, representative SORT gene lists were defined before bulk survival modeling. For each signal GEP, genes were first restricted to those with positive SORT profile values.

The support scores were calculated from the 39 signal components after excluding background GEP00. Let $W_c$ be the column-centered signal-activity matrix and $q_g$ the vector of signal profile values for gene $g$. Let $w_k$ denote the centered activity vector of signal GEP $k$, and let $\widehat{x}_g = W_c q_g = \sum_j q_{gj} w_j$ denote the centered fitted expression of gene $g$ reconstructed from all signal GEPs. The fitted contribution of GEP $k$ to this expression is $q_{gk} w_k$. We defined

$$R_{\text{var},gk} = \frac{\text{Var}(q_{gk} w_k)}{\text{Var}(\widehat{x}_g)}, \qquad C_{\text{recon},gk} = \text{sign}(q_{gk}) \, \text{cor}(w_k, \widehat{x}_g).$$

$R_{\text{var},gk}$ compares the variance of the individual GEP contribution with the gene's total fitted variance. $C_{\text{recon},gk}$ measures how closely the fitted expression of the gene follows activity $k$, accounting for the direction of its profile value.

Genes were then ranked by the sum of their descending ranks for the profile value, $R_{\text{var}}$ and $C_{\text{recon}}$. Within the positive-profile genes of each component, genes at or above the 90th percentile for all three quantities formed the high-support set. When more than 10 genes passed, selection was restricted to this set; otherwise all positive-profile genes were ranked. The fixed top-50 gene list used by the downstream analysis was retained as the projection input for each component.

Representative SORT gene lists were projected into TCGA, CPTAC, QCMG and PanCuRx PDAC cohorts. These cohorts were not used for SORT fitting, spatial GEP annotation or representative gene-list definition before projection. Cohort sizes, events and available covariates are summarized in Supplementary Table 2. For each GEP and cohort, normalized expression values were averaged across genes present in both the representative GEP gene list and the cohort to obtain one bulk projection score per patient. These GEP-specific scores were then z-scored across patients within the same cohort.

**Survival association across cohorts.** For each of the 39 signal GEPs, we fitted a Cox proportional hazards model relating the bulk projection score to overall survival within each cohort and adjusting for available age, sex and stage [2]. Log hazard ratios and standard errors from the four cohorts were combined by inverse variance meta-analysis under a fixed-effect model. Meta-analysis $P$ values were adjusted by the Benjamini–Hochberg procedure across the 39 GEPs.

**Descriptive Kaplan–Meier display.** Kaplan–Meier curves pooled the four cohorts after standardizing scores within each cohort. The four GEPs with the smallest FDR values were displayed using a global median split, which was also applied to all 17 GEPs with all-39 FDR $< 0.05$ in Supplementary Fig. S14. Log-rank $P$ values were used descriptively.

#### Supplementary Note 5. Cross-species cerebellum analysis

**Input and cross-species activity summaries.** The original cross-species cerebellar SORT fit used three analyzed sections from one animal of each species: macaque, marmoset and mouse. Gene symbols were converted to uppercase and intersected directly across the nine sections. No one-to-one orthology mapping was applied during SORT fitting. Each section was normalized to a target sum of $10^4$, log-transformed and analyzed using 4,000 highly variable genes at rank $K = 25$.

GEP activity was summarized along the normalized distance-to-layer coordinate supplied by the original atlas. We used 160 equally spaced bins spanning the pooled finite-distance 0.5th–99.5th percentile interval. Bins with fewer than 20 locations within a section were treated as missing. Remaining values were smoothed with a nine-bin moving average weighted by the number of locations. Missing bins received zero weight, and edge windows were shortened. The three smoothed section curves were averaged within species using bin-level location counts as weights. Each species profile was z-scored across available cortical-depth bins. Spatial-profile similarity was the mean Pearson correlation across the three species pairs.

Activity magnitude was summarized from the unstandardized smoothed profiles. We calculated the arithmetic mean of the finite values whose bin centers fell within $[-1, 1]$. All section profiles contained the same 119 finite bin centers in this interval. The three section-level values were averaged within each species. Variation in activity magnitude was then summarized descriptively by the population coefficient of variation (CV) of the three species means. No interpolation, extrapolation or additional count weighting was applied at this stage.

**Fixed GEP02 and GEP14 gene sets.** After conversion to uppercase symbols, we defined a support-aware candidate pool of 60 positive-weight genes for each fitted component using transcriptomic-profile weight, component-attributable reconstructed variance $R_{\text{var}}$ and reconstructed expression–activity correlation $C_{\text{recon}}$. For GEP02 and GEP14, the 20 genes with the largest transcriptomic-profile weights within this pool were retained. Candidate pools, gene membership and order were fixed from the original SORT fit before cross-species detection effects were examined.

**Additional animals and fixed-profile projection.** The extended analysis added three adjacent sections from one additional animal of each species. The standardized uppercase symbols were intersected across all 18 sections to define a common normalization and projection universe. Counts at each location were normalized to a total of $10^4$ over this fixed universe and log-transformed. Genes from the SORT fit that were present in every section were retained. These steps aligned measured features across sections but did not assert one-to-one orthology.

The learned $W$ activity was used for domains in the animals included in the original SORT fit. SORT was not refitted to the additional animals. Instead, the fitted $Q$ matrix was held fixed and used to calculate nonnegative projection scores. For each tested gene set, its 20 genes were removed from every projected $Q$ column before projection. For location $i$, let $x_i$ denote its normalized expression vector and let $q_k^{(-G)}$ denote the profile after masking the tested genes. The nonnegative projection score was $\widehat{a}_{ik} = \max\{0, x_i^\top q_k^{(-G)}/[(q_k^{(-G)})^\top q_k^{(-G)}]\}$. These gene-masked projection coefficients located corresponding domains in the additional animals without using the evaluated genes or refitting SORT.

Within each section, fitting-animal domains contained locations at or above the 90th percentile of strictly positive learned activities, whereas additional-animal domains used the corresponding percentile of strictly positive masked projection scores. Detection fraction was calculated separately for each gene and section as the proportion of domain locations with nonzero expression. For each animal set, the three section values were averaged within species. The gene-level effect was the marmoset mean minus the mean of the macaque and mouse means. The gene-set effect was the mean of the 20 gene-level effects. Heatmap values were standardized across the 18 sections separately for each gene.

**Expression-matched control sets.** Matched calibration was performed separately for the animals used for SORT fitting and the additional animals. Global detection prevalence and mean expression among detected locations were converted to percentile ranks. For each tested gene, controls were drawn without replacement from the 100 nearest eligible genes in this two-dimensional rank space. Genes with the strongest profile-weight or combined-support rankings for either GEP were excluded from the control pool. Each of 10,000 control sets contained one match per tested gene. Outcome effects were not used for matching. The null statistic was the mean detection effect of the 20 selected controls in the same target-GEP domain. The empirical upper-tail probability was $(1+b)/(10{,}000+1)$, where $b$ denotes matched sets with a mean at least as large as the observed mean.

**Ensembl one-to-one orthology sensitivity.** Orthology was assessed after the primary analysis as an external annotation audit and was not used in SORT fitting or primary gene selection. Each selected standardized uppercase symbol was submitted using *Homo sapiens* as the anchor species and queried through the Ensembl release 116 REST homology endpoint against *Macaca mulatta*, *Callithrix jacchus* and *Mus musculus* [3, 4]. A gene passed the strict sensitivity filter only when all three queries succeeded and each returned exactly one relationship of type `ortholog_one2one`. Sixteen GEP02 genes and 18 GEP14 genes passed this filter. Their mean detection effects were recalculated from the primary domain-level effects; SORT was not refitted and spatial domains were not redefined.

**Reciprocal GEP00–GEP01 contrast.** The GEP01 analysis retained the original nine sections used for SORT fitting and the fixed GEP00–GEP01 component pair. GEP00 denotes the fitted background component. Candidate genes with positive profile weights were required to pass the same $Q$, $R_{\text{var}}$ and $C_{\text{recon}}$ support criterion. Eligible genes were ranked by the combined-support order, and 12 genes were fixed from each component before detection testing. The raw activity sum of GEP00 and GEP01 defined a shared granule-layer domain. One threshold was calculated as the 90th percentile of strictly positive composite activity pooled across the nine sections. Locations at or above this threshold were retained within each section. If fewer than 150 locations were selected and positive activity existed, the top $\min(150, n_{\text{positive}})$ positive locations were used.

Detection fraction was calculated for each gene and section. Effects were the macaque mean minus the mean of marmoset and mouse. A one-sided exact test compared the three macaque section values with the six comparison-section values over all 84 assignments of three target labels. Benjamini–Hochberg correction was applied across the fixed 24-gene family. These section-level statistics and the reciprocal group means provide descriptive support within the sampled animals rather than population-level species inference.

**Empirical scalability.** Computational scaling was assessed at six balanced input sizes from 0.1 to 3 section-equivalents per species, corresponding to 51,997–1,546,637 spatial locations. The 0.1-, 0.2- and 0.5-equivalent inputs used centered, spatially contiguous rectangular crops of one representative section per species. Larger inputs used one, two or three complete sections per species. The 4,000-gene feature set, $K = 25$, model settings, software environment and NVIDIA A30 graphics processing unit (GPU) were held fixed. End-to-end time was measured around the complete run, and model-fitting time was recorded internally. Each input size was run once, providing single-run empirical runtime under fixed settings across the tested input sizes.

#### Supplementary Note 6. Mouse embryogenesis analysis

**Input and alignment.** The mouse embryogenesis fit contained 520,389 aligned spatial locations and 8,000 genes included in the SORT fit. SORT was fitted jointly at rank $K = 60$, with component 0 treated as background and GEP01–GEP59 treated as signal GEPs.

**Temporal summaries and spatial display.** For each signal GEP, mean activity was calculated across all retained locations within each stage and divided by that GEP's maximum stage mean. For display, normalized stage profiles were ordered from early peaks to late increases. The groups shown in Supplementary Fig. S19 are descriptive and do not define biological classes. Within-group order was determined by average-linkage clustering using correlation distance. The main dynamic panel contained the 25 GEPs with the largest across-stage ranges. GEP02 and GEP25 were additionally included because they are examined in subsequent panels and were absent from the range-based set. Supplementary Fig. S19 shows all 59 signal GEPs.

For each component, one color scale spanning its pooled 0.5th–99.5th activity percentiles was used across all stages. Anatomical annotations were obtained from the source atlas and were not used to fit SORT. Stage-by-annotation strata containing fewer than 50 locations were treated as unsupported.

**Literature-informed gene-set concordance.** Literature-informed gene sets containing genes with established roles in the corresponding tissue or developmental process were fixed before the concordance summaries and are documented with their biological rationale and citations in Supplementary Table 5. For GEP25, GEP31 and GEP02, stored log-transformed expression was standardized separately for each gene within each stage-by-annotation stratum and averaged across genes to form the gene-set score. Raw concordance was the Spearman correlation between continuous activity and gene-set score. For the detected-gene-coverage sensitivity, activity, gene-set score and detected-gene count were rank-transformed; ranked activity and ranked gene-set score were each regressed on ranked detected-gene count, and the Pearson correlation between the residual vectors was reported as the coverage-adjusted rank correlation.

**GEP-specific analyses.** GEP25 concordance was evaluated in nine stage-by-annotation contexts spanning E11.5 Sclerotome and Head mesenchyme, E12.5 Cartilage primordium and later Cartilage or Cartilage primordium. Activity-gradient summaries divided ranked GEP25 activity into ten equal-count bins and reported the median and first to third quartiles of the gene-set score in each bin. GEP31 was evaluated separately within Surface ectoderm, Epidermis and Mucosal epithelium; these annotations were not pooled.

For GEP02, continuous concordance was evaluated within Brain at each stage. Threshold sensitivity compared the median neural-associated gene-set score between the within-stage Brain activity groups defined by 20th/80th percentile, 25th/75th percentile or tertile groups. For spatial sensitivity, each stage-specific Brain bounding box was divided into $6\times6$, $8\times8$ or $10\times10$ normalized grids. Blocks with fewer than 20 locations were omitted, and the reported block-level Spearman correlation compared median GEP02 activity with the median neural-associated gene-set score across the retained blocks.

For GEP06 and GEP43, positive transcriptomic profiles were defined as $Q_k^+ = \max(Q_k, 0)$. Profile similarity was summarized by cosine similarity and by the Jaccard index of the top 50 positive-weight genes.

Gene-set concordance was evaluated across all locations within each of the eight stages, using the fixed sets in Supplementary Table 5. The GEP06 set comprised *Myh3*, *Tnnt3*, *Actn2* and *Des*, and the GEP43 set comprised *Acta2*, *Myl9*, *Actg2* and *Tagln*. Within each stage, log-transformed expression was standardized separately for each gene and averaged across the corresponding set. Each GEP's unscaled activity was compared with its own gene-set score using the raw and coverage-adjusted rank correlations described above. All 520,389 locations were included across the eight stages.

For the continuous activity maps, each GEP was scaled as $\widetilde{W_k} = \min\{W_k/q_{0.99}(W_k), 1\}$, with the 99th percentile calculated over all locations. For the overlap maps, high activity was defined independently for each GEP within each stage as an unscaled activity value at or above the 90th percentile across the whole section. Locations were classified as high for GEP06 only, GEP43 only, both or neither.

Colocalization scores and high-activity Jaccard indices were calculated over each whole section using the definitions and weights given in the PDAC spatial colocalization analysis.

These post-fit summaries were descriptive, and no location-level $P$ values were calculated.

#### Supplementary Note 7. Positive-profile over-representation analysis

**Gene-set definition and enrichment universe.** For each signal GEP in the PDAC and mouse embryogenesis fits, genes were ranked by their fitted transcriptomic-profile loading in $Q$. The primary set contained positive-loading genes in the top 2% of the complete ranking of genes included in the corresponding SORT fit (200 genes for PDAC and 160 genes for mouse embryogenesis). Top 1% and top 5% sets were analyzed as sensitivity sets (100 and 500 genes for PDAC; 80 and 400 genes for mouse embryogenesis). The enrichment universe was the complete set of genes included in the corresponding SORT fit. Hallmark and Reactome gene sets from MSigDB v7.5.1 were used for PDAC, and Gene Ontology Biological Process terms from `org.Mm.eg.db` v3.18.0 were used for mouse embryogenesis. Only gene sets containing 10–500 genes from the fit were tested.

**Statistical analysis and display.**

We tested each positive profile set for over-representation using a one-sided hypergeometric test. Benjamini–Hochberg correction was applied across the complete family of signal-GEP–pathway tests, separately for each dataset, pathway collection and profile-size cutoff. Tests with zero overlap or nonsignificant results remained in the correction family. Fold enrichment was the observed overlap divided by its expectation under random sampling from the complete set of genes included in the fit. A pathway was eligible for display when the primary top 2% test had global FDR $< 0.05$ and the same pathway was also supported at top 1% or top 5%. Supplementary Figs. S10 and S22 show nonredundant terms selected from these eligible results for the GEPs discussed in the main text. Gray crosses mark cutoff-specific results with global FDR $\geq 0.05$.

For the cerebellar catalog, the enrichment universe comprised the 4,000 genes included in the SORT fit. The reference tables summarize significant terms from the primary top 2% analysis, whereas Supplementary Figs. S10 and S22 additionally show sensitivity to the gene-set cutoff.

**Relationship to the spatial-activity annotation.** ORA used genes with positive $Q$ loadings and complemented the $W$-based gene and signature annotations used for PDAC.

#### Supplementary Figures

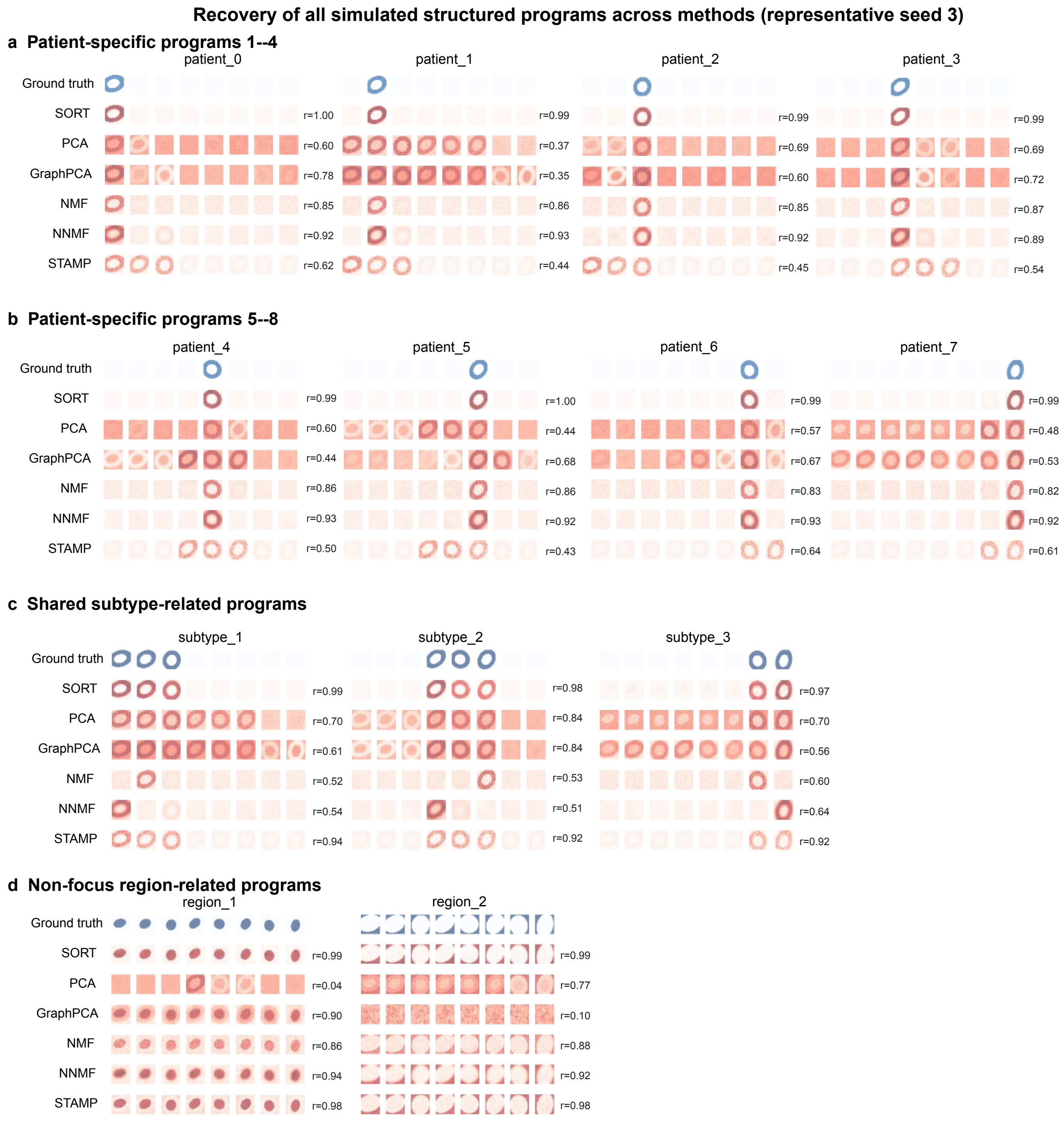


Supplementary Figure S1: **Recovery of all simulated ground-truth gene expression programs across methods.** The 13 ground-truth GEPs from seed 3 are shown with their best-matched components from SORT, PCA, GraphPCA, NMF, NNMF and STAMP. **(a,b)** Eight patient-specific programs. **(c)** Three subtype-related programs. **(d)** Two non-focus region-related programs. Matches were selected by signed Pearson correlation across pooled locations and sections. The displayed $r$ is the matched correlation. Maps are scaled for display.

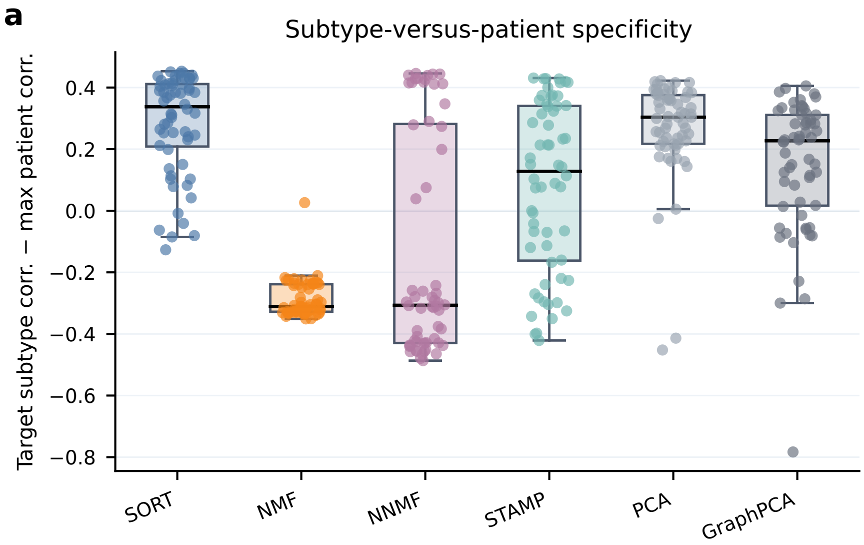


Supplementary Figure S2: **Specificity of recovered subtype programs relative to patient-specific programs.** For each inferred subtype gene expression program (GEP) already selected as the best match to a ground-truth subtype GEP in Main Fig. 2e, the plotted value is its correlation with the matching ground-truth subtype GEP minus its maximum correlation with any patient-specific ground-truth GEP. No additional component matching was performed. Positive values indicate that the recovered component is more specific to the intended subtype program than to any patient-specific program.

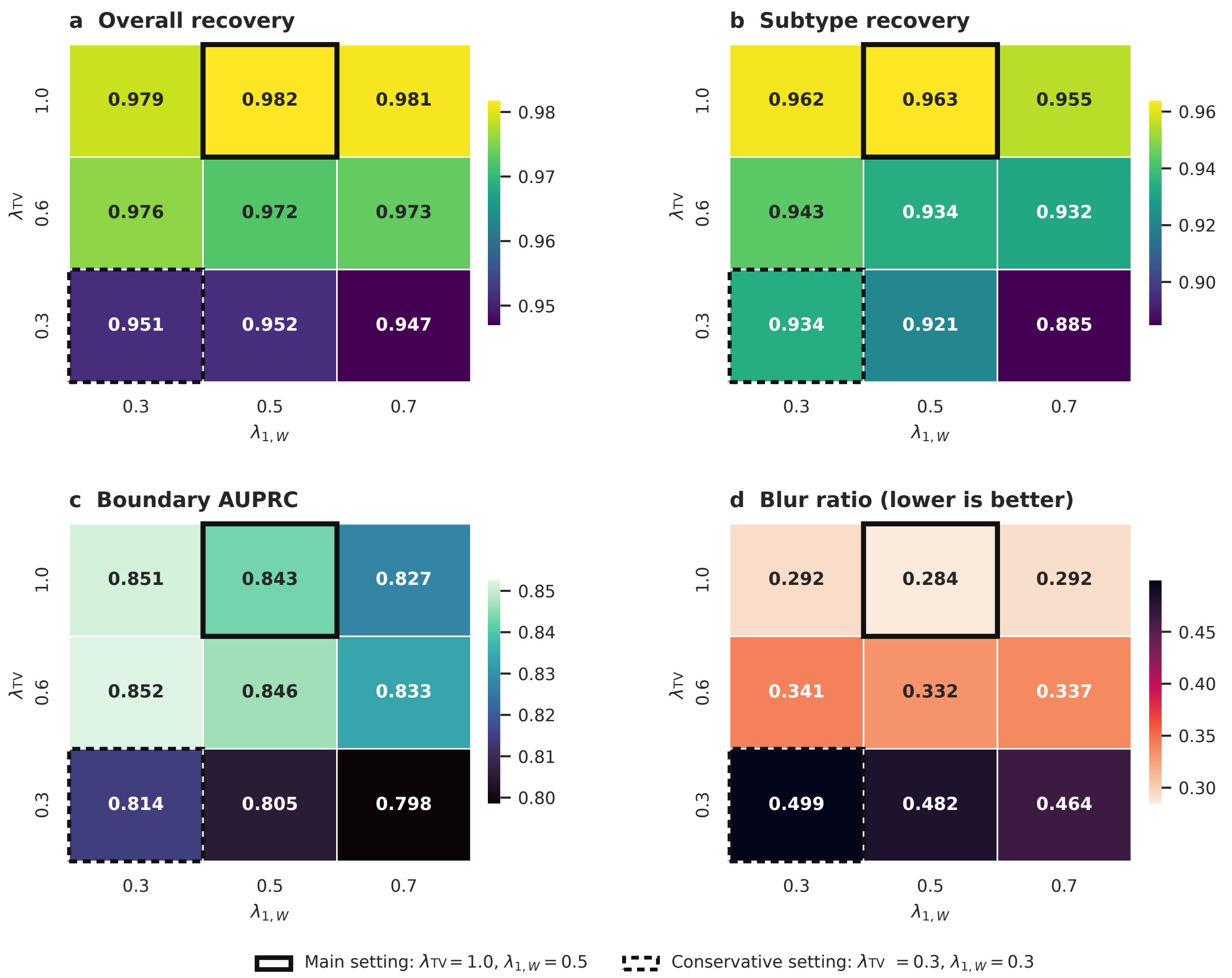


Supplementary Figure S3: **Sensitivity of simulation performance to graph total variation and spatial-activity sparsity.** Heatmaps show median performance across 10 matched simulation seeds for $\lambda_{\text{TV}} \in \{0.3, 0.6, 1.0\}$ and $\lambda_{1,W} \in \{0.3, 0.5, 0.7\}$. Other settings match the main simulation benchmark. **(a)** Overall gene expression program (GEP) recovery. **(b)** Subtype-related GEP recovery. **(c)** Boundary area under the precision–recall curve (AUPRC). **(d)** Blur ratio. Solid and dashed outlines mark the main and conservative settings. Higher recovery and AUPRC values are better, whereas lower blur ratios are better.

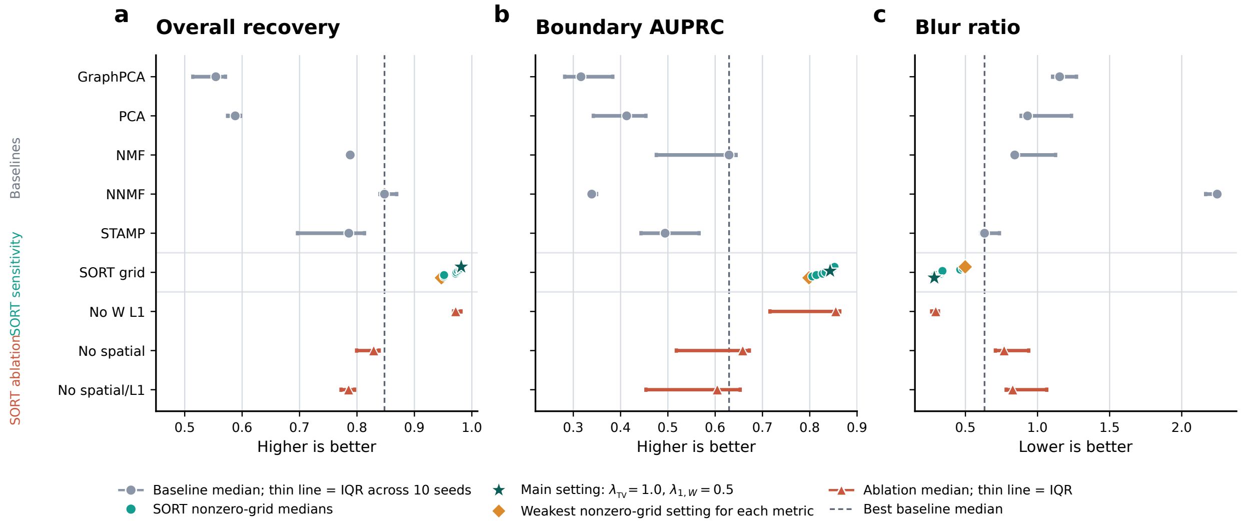


Supplementary Figure S4: **Ablation analysis of SORT regularization in simulation. (a)** Overall gene expression program (GEP) recovery. **(b)** Boundary area under the precision–recall curve (AUPRC). **(c)** Blur ratio. Metrics are compared across baseline methods, the nonzero SORT parameter grid and variants without graph total variation, $W$ sparsity or both terms. Points and thin lines show medians and interquartile ranges across 10 matched seeds. The main SORT setting and the weakest nonzero-grid setting for each metric are marked. Higher recovery and AUPRC values are better, whereas lower blur ratios are better.

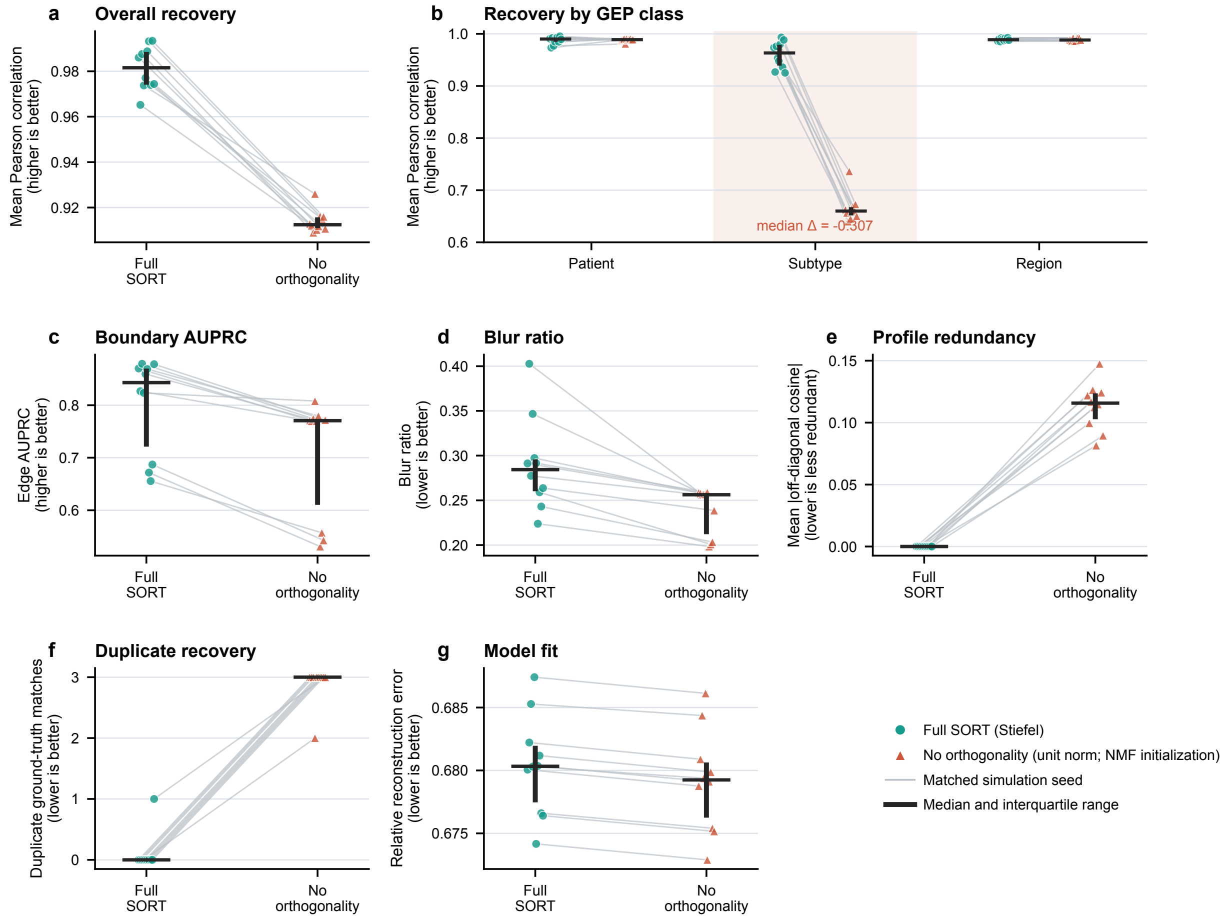


Supplementary Figure S5: **The orthogonality design supports recovery of weaker subtype programs in matched simulations.** Full SORT was compared with a matched variant that retained unit-norm profiles but removed pairwise orthogonality and used standard NMF initialization. **(a)** Mean Pearson correlation across all 13 ground-truth programs. **(b)** Recovery by program class. **(c)** Boundary edge area under the precision–recall curve (AUPRC). **(d)** Blur ratio. **(e)** Mean absolute cosine similarity among signal profiles. **(f)** Duplicate best matches. **(g)** Relative Frobenius reconstruction error. Higher values are better in **(a–c)**, and lower values in **(d–g)**. Points show ten paired simulation datasets. Gray lines connect matched seeds, and black summaries show medians and interquartile ranges.

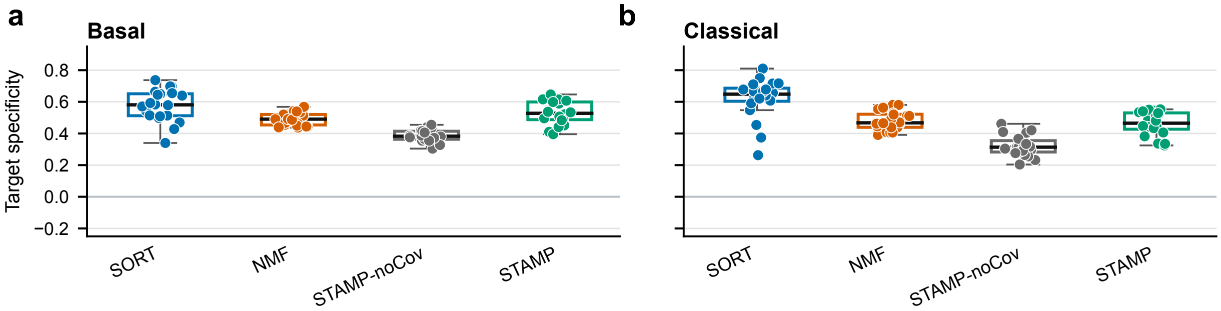


Supplementary Figure S6: **Subtype-reference components show greater target specificity with SORT.** Target specificity across 20 resampling iterations is shown for **(a)** basal and **(b)** classical references. Components were matched using correlation with the target score alone. Specificity was then calculated as the target correlation minus the absolute correlation with the alternative subtype score. Higher values indicate greater separation between the basal and classical references.

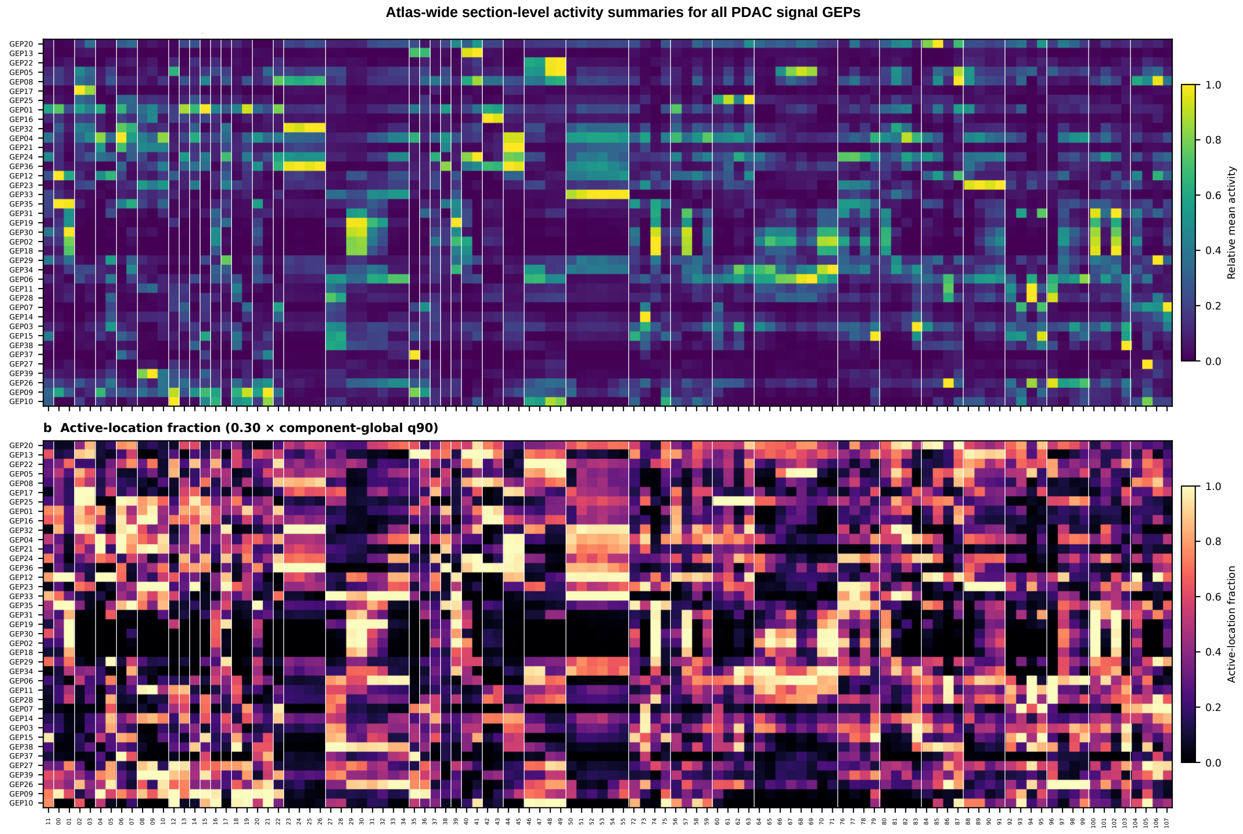


Supplementary Figure S7: **Section-level activity summaries for all pancreatic ductal adenocarcinoma gene expression programs.** Rows show all 39 signal gene expression programs (GEPs), and columns show all 108 Visium sections ordered by patient. **(a)** Mean fitted activity, scaled to the range 0–1 across sections within each GEP. **(b)** Fraction of locations in each section with activity at or above 0.30 times that component's atlas-wide 90th percentile. Vertical lines mark patient boundaries. Rows are ordered by similarity of their section-level mean activities.

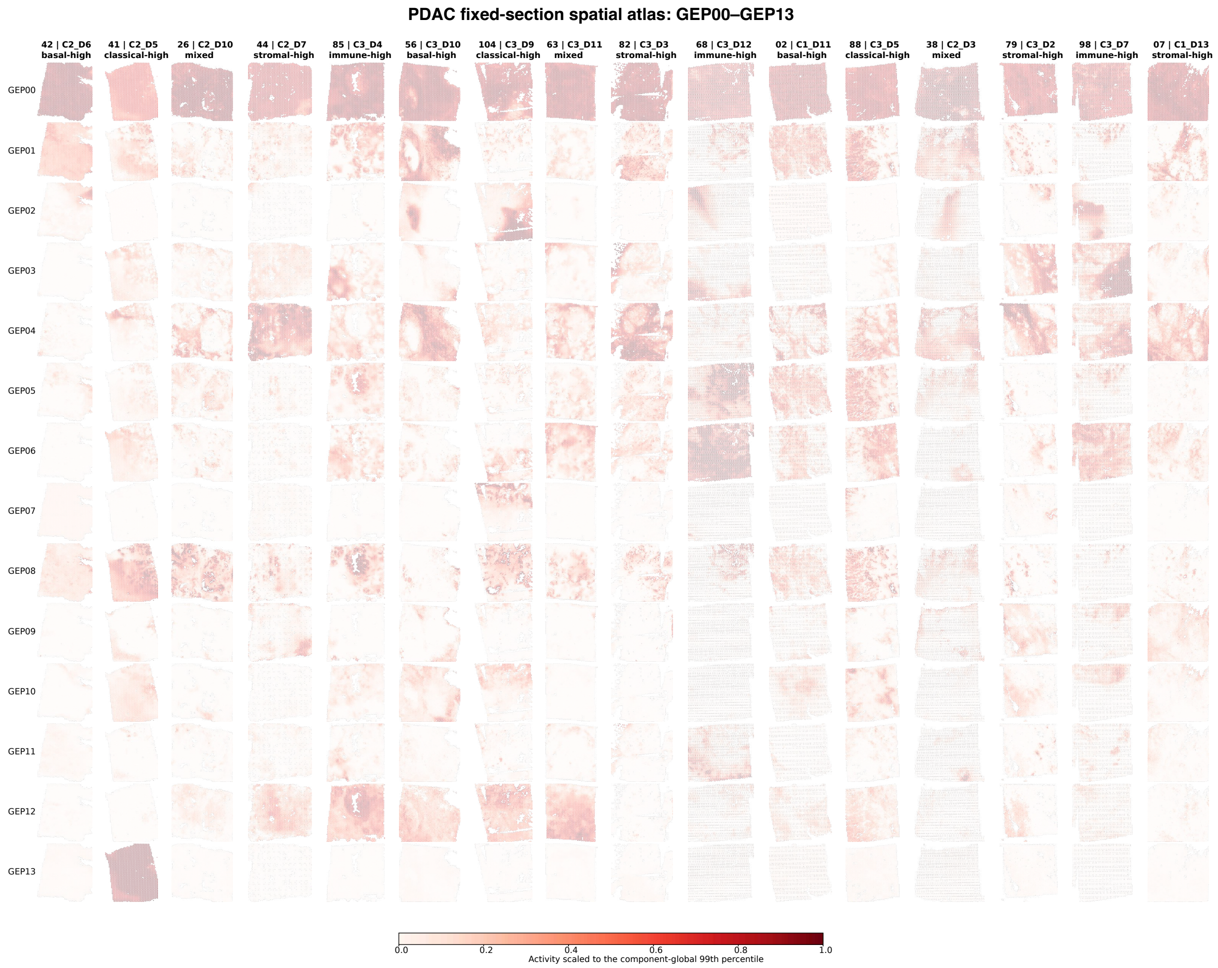


Supplementary Figure S8: **Spatial activity maps for all pancreatic ductal adenocarcinoma SORT components in a fixed section panel.** All 40 fitted components, including GEP00, are shown across 16 Visium sections from 16 patients on three pages. This page shows GEP00–GEP13. The sections were selected independently of SORT activity. Activity is scaled within each component to its global 99th percentile across all 108 sections.

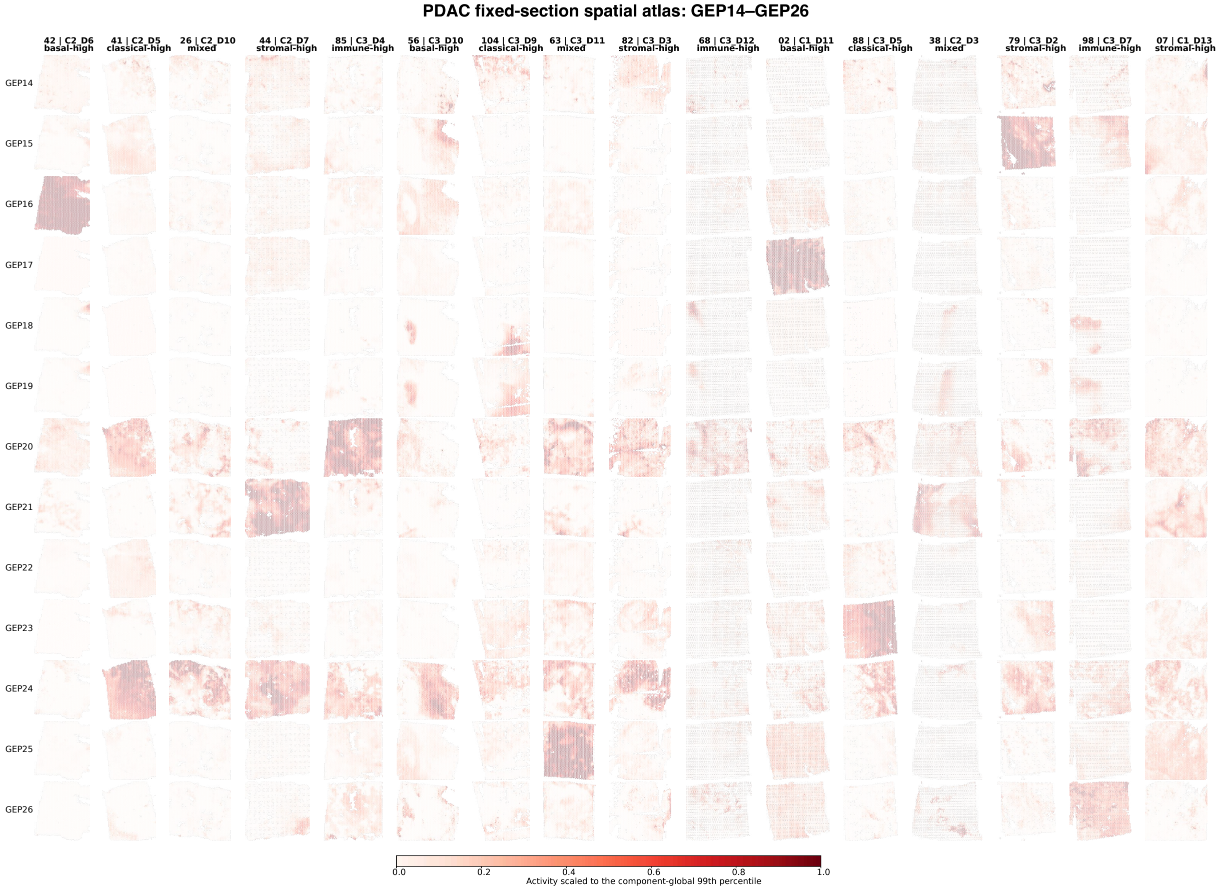


Supplementary Figure S8: **Spatial activity maps for pancreatic ductal adenocarcinoma SORT components in a fixed section panel (continued).** GEP14–GEP26 are shown across the identical 16-section panel using the same component-specific scales.

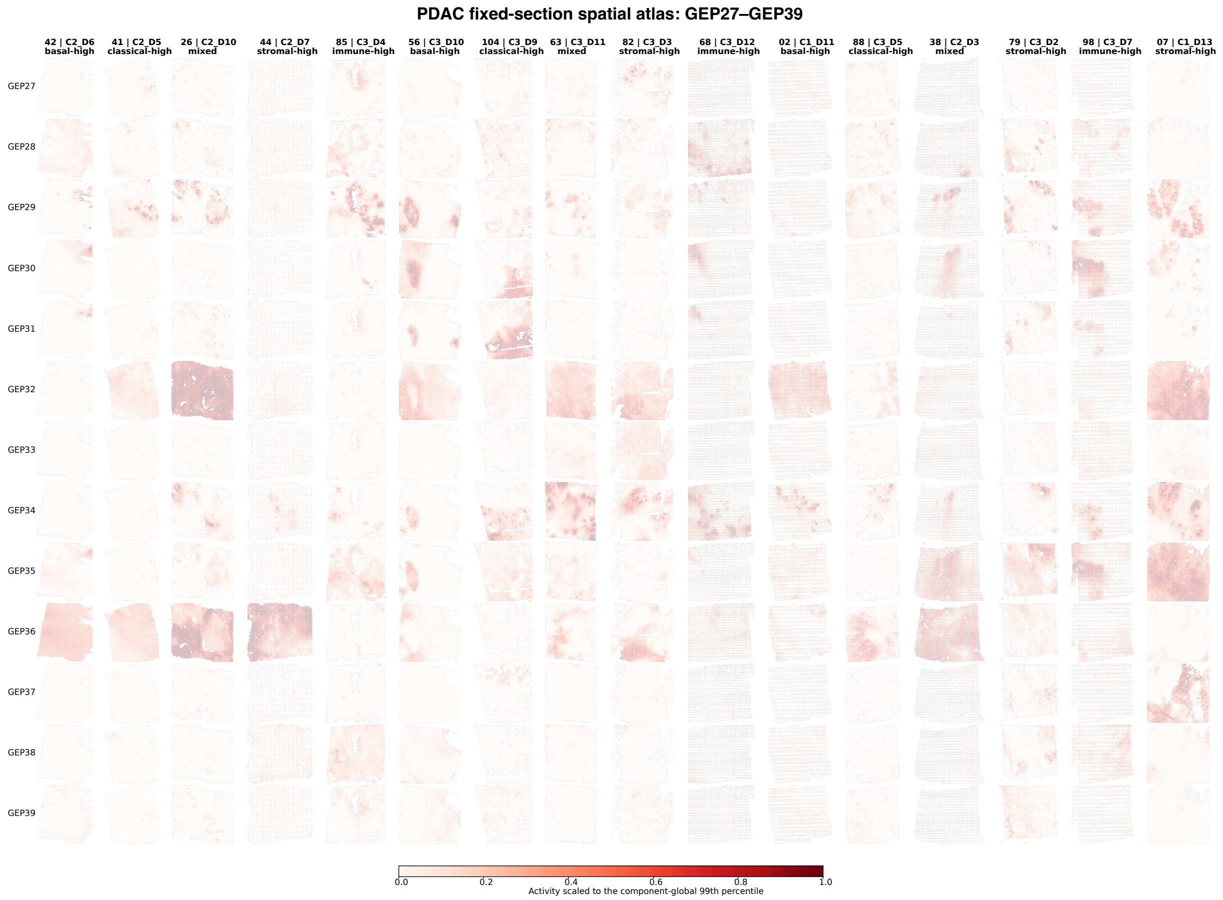


Supplementary Figure S8: **Spatial activity maps for pancreatic ductal adenocarcinoma SORT components in a fixed section panel (continued).** GEP27–GEP39 are shown across the identical 16-section panel using the same component-specific scales.

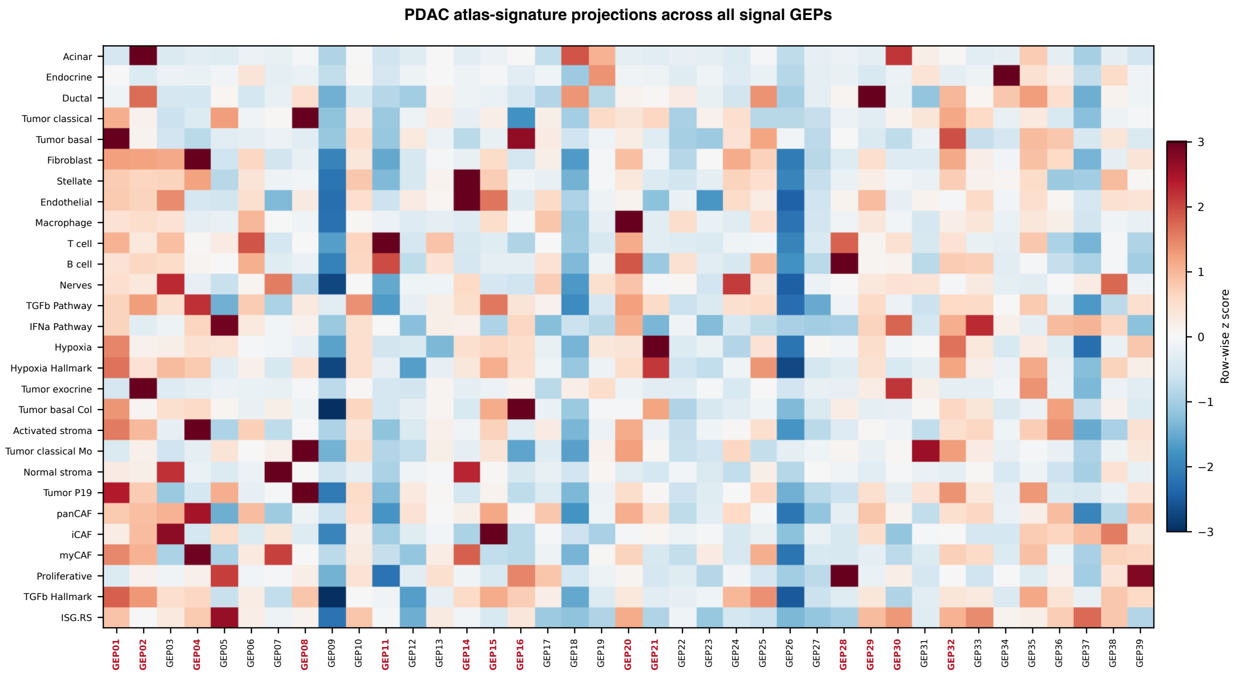


Supplementary Figure S9: **Atlas-signature support for annotation of pancreatic ductal adenocarcinoma programs.** Rows show 28 PDAC and tumor microenvironment signatures, and columns show all 39 signal GEPs. Values are signed mean $B_{gk}$ coefficients across overlapping genes, standardized within each signature across GEPs. Red labels mark GEPs displayed in main Fig. 4.

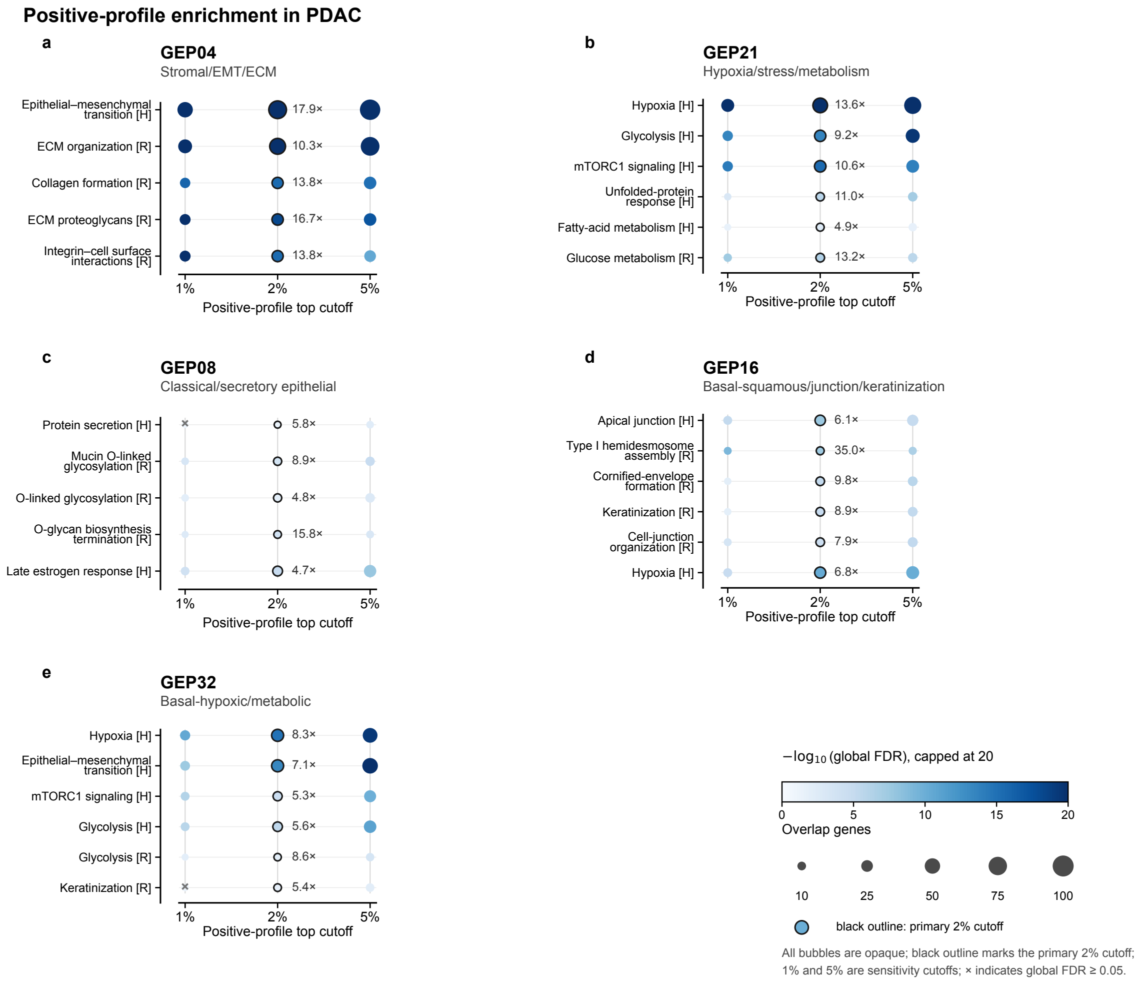


Supplementary Figure S10: **Positive-profile pathway enrichment in pancreatic ductal adenocarcinoma. (a–e)** Results for gene expression programs (GEPs) GEP04, GEP21, GEP08, GEP16 and GEP32. Rows are selected nonredundant Hallmark or Reactome pathways. Columns use the top 1%, 2% and 5% positive-loading genes in $Q$, with the top 2% as the primary analysis. Outlined points are labeled with fold enrichment (FE). Color represents $-\log_{10}$ global false discovery rate (FDR), capped at 20. Point size represents the number of overlapping genes. Gray crosses mark results with global FDR $\geq 0.05$. Benjamini–Hochberg correction was applied across all signal-GEP–pathway tests within each collection and cutoff.

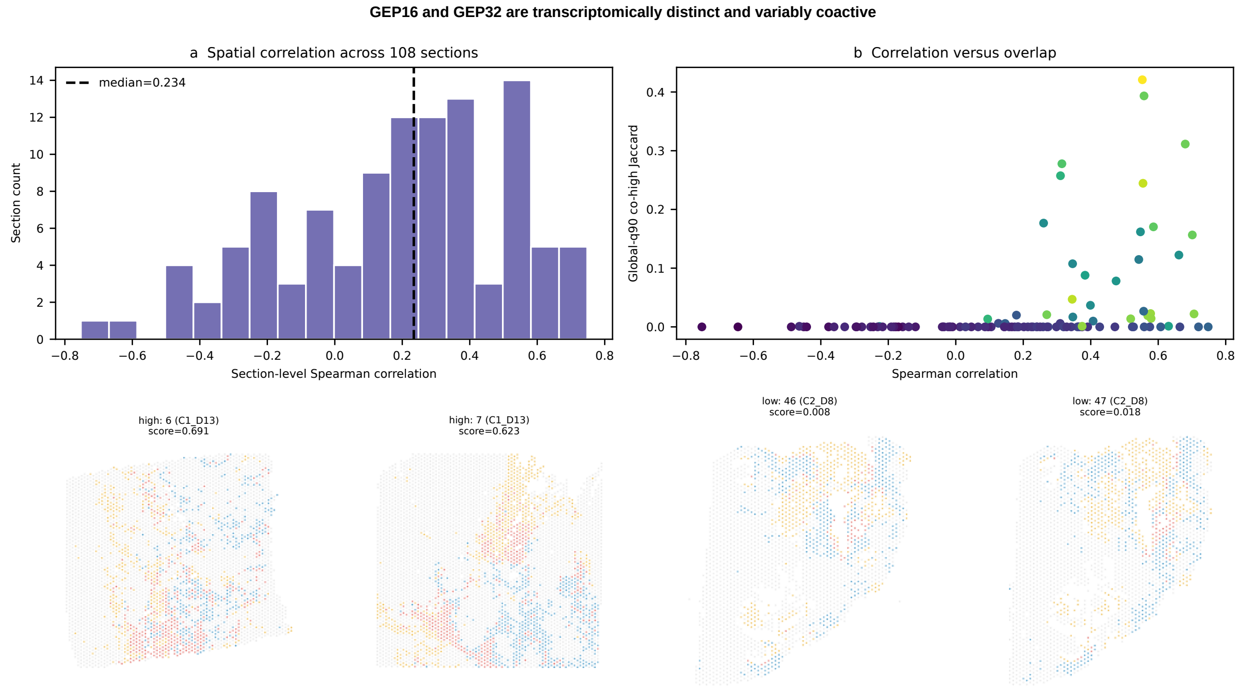


Supplementary Figure S11: **Spatial coactivity of GEP16 and GEP32 across pancreatic ductal adenocarcinoma sections.** **(a)** Distribution of section-level Spearman correlations between their activities across all 108 sections. **(b)** Relationship between spatial correlation and global 90th-percentile co-high Jaccard overlap. The lower maps show representative sections with higher or lower coactivity. Colors denote GEP16 high only (orange), GEP32 high only (blue), both high (red), and neither high (gray).

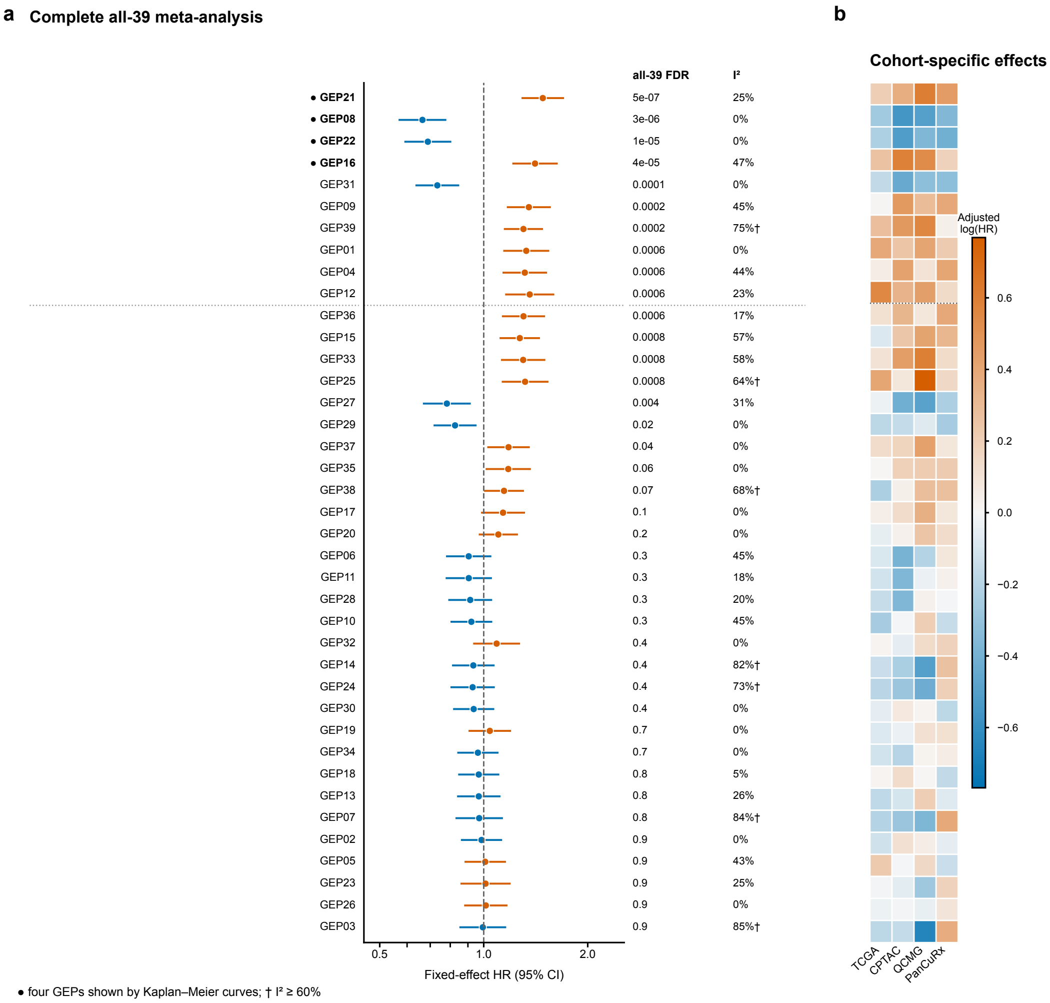


Supplementary Figure S12: **Survival associations for all pancreatic ductal adenocarcinoma signal programs.** All 39 signal gene expression programs (GEPs) are ordered by Benjamini–Hochberg false discovery rate (FDR). The forest plot shows hazard ratios and 95% confidence intervals from inverse variance meta-analysis under a fixed-effect model. Blue and orange indicate protective and adverse associations. Adjacent columns report FDR and $I^2$. Daggers mark $I^2 \geq 60\%$. The heatmap shows covariate-adjusted log hazard ratios in TCGA, CPTAC, QCMG and PanCuRx. Filled bullets mark the four GEPs used for Kaplan–Meier display. The dotted line separates the ten associations shown in main Fig. 4d,e.

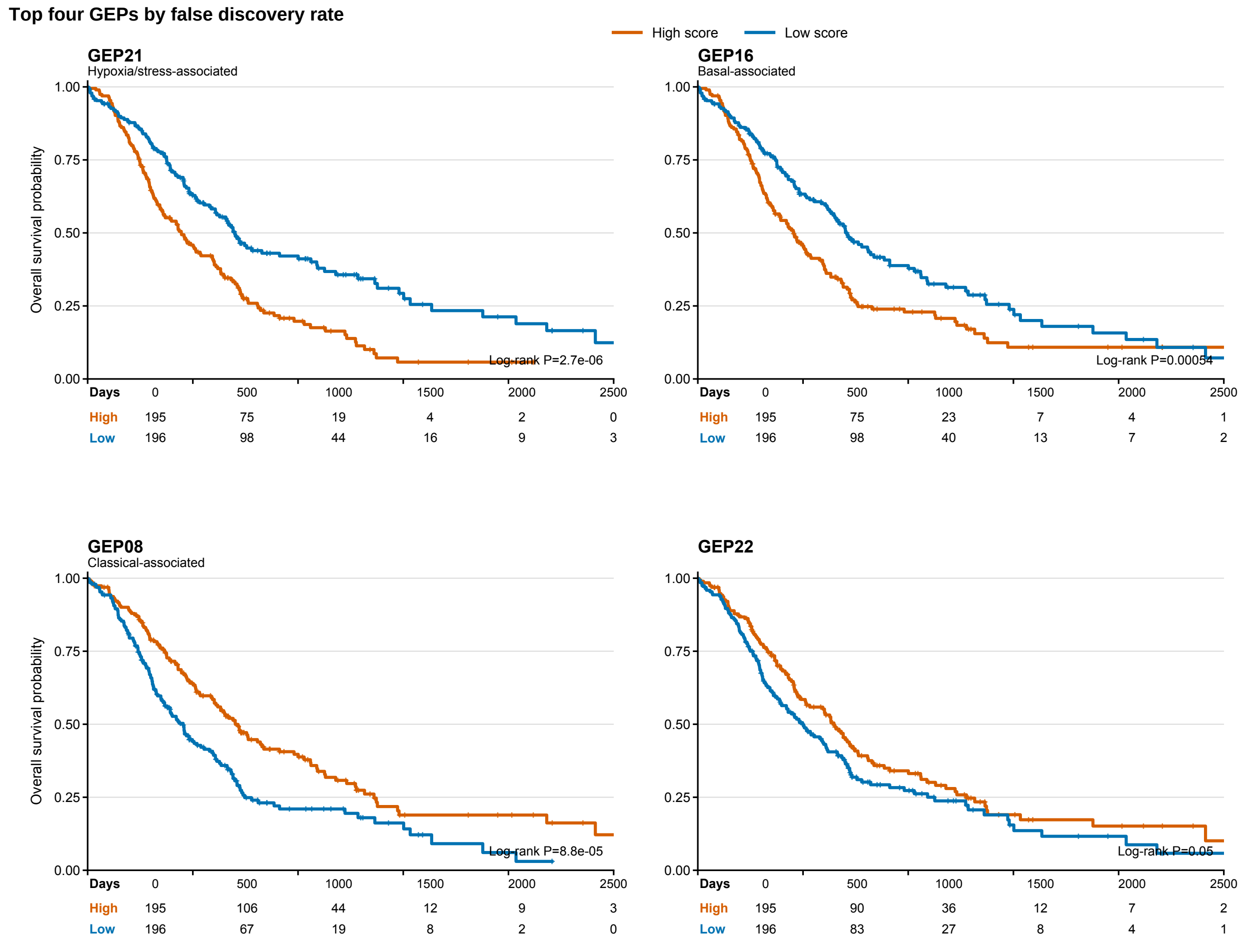


Supplementary Figure S13: **Kaplan–Meier displays for the four programs with the smallest false discovery rates.** Scores were standardized within each cohort before pooling and split at the global median (High, $n = 195$; Low, $n = 196$; 391 patients and 260 events). Censor marks and complete risk tables are shown. Curves complement the continuous Cox models and meta-analysis. Log-rank $P$ values are descriptive.

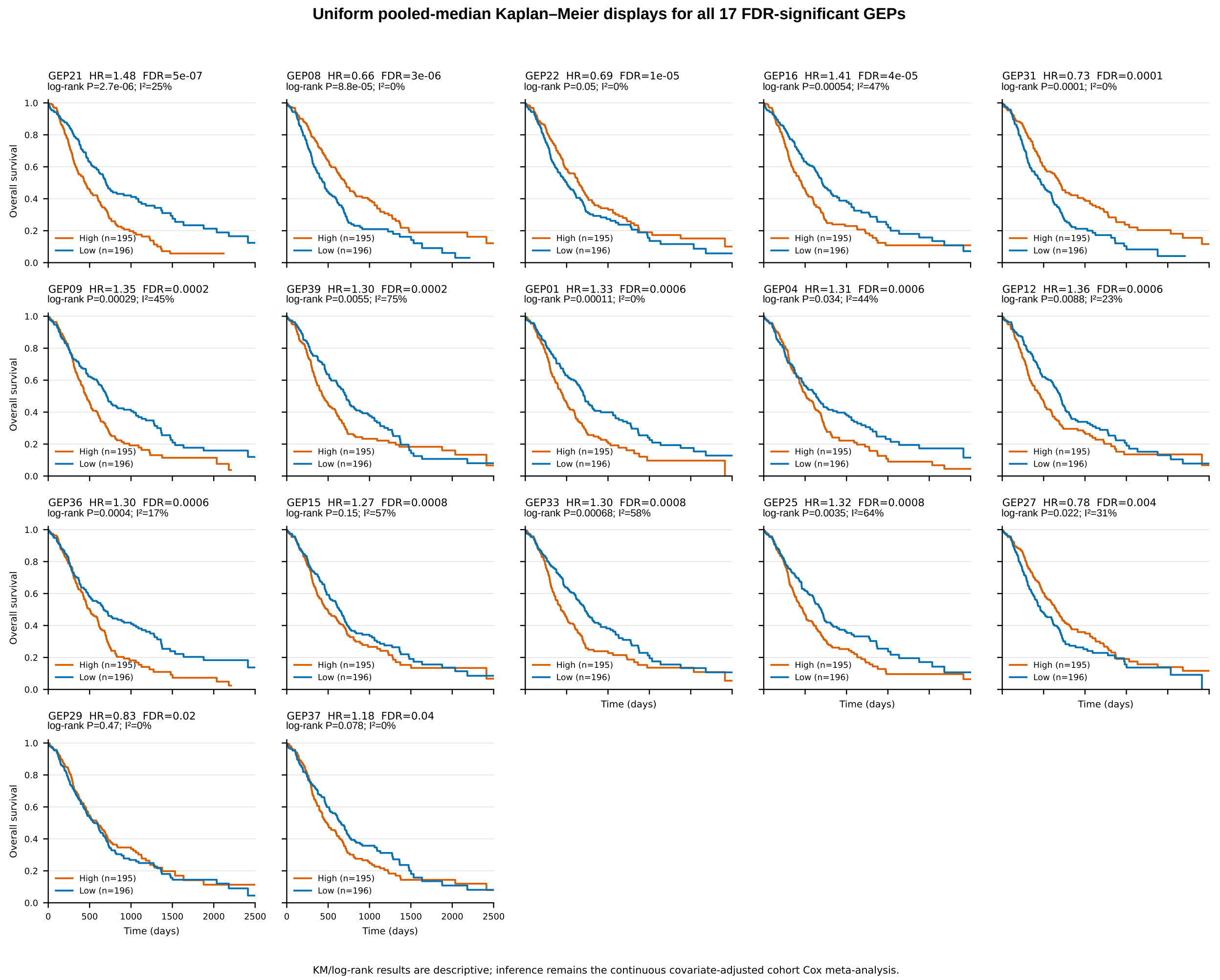


Supplementary Figure S14: **Kaplan–Meier displays for all programs with false discovery rate below 0.05.** The 17 GEPs are shown in FDR order. Scores were standardized within each cohort before pooling and split at the global median. Each panel reports the fixed-effect meta-analysis hazard ratio, FDR, $I^2$ and descriptive log-rank $P$ value.

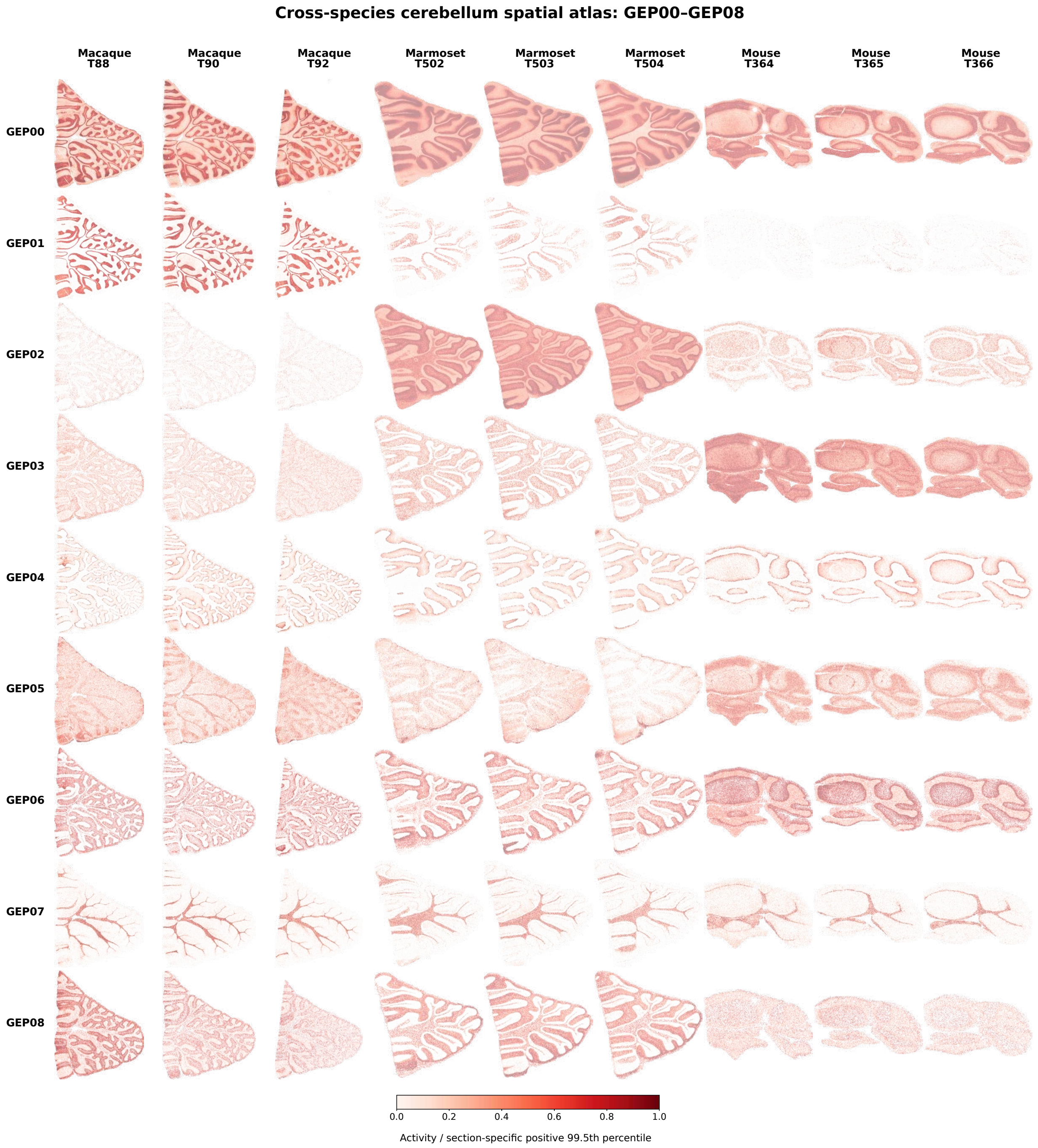


Supplementary Figure S15: **Spatial activity maps for all fitted cerebellar SORT components in selected specimens.** All 25 components are shown across nine Stereo-seq sections, comprising three adjacent sections from one macaque, one marmoset and one mouse, on three pages. This page shows GEP00–GEP08. Positive activities were scaled to the 99.5th percentile within each section and component. Low-activity background locations were subsampled only for rendering.

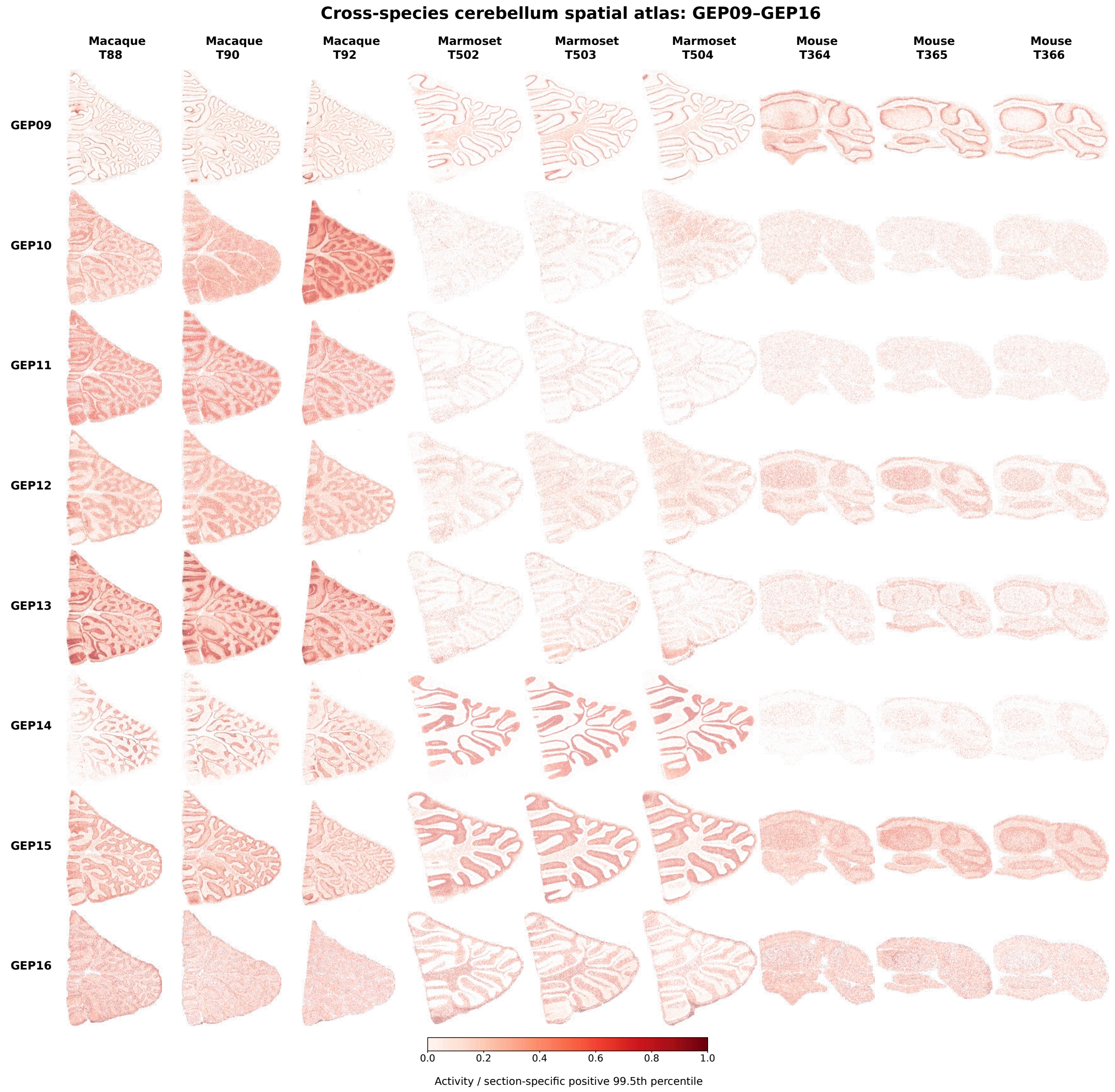


Supplementary Figure S15: **Spatial activity maps for all fitted cerebellar SORT components in selected specimens (continued).** GEP09–GEP16 are shown across the same nine sections using the same rendering procedure.

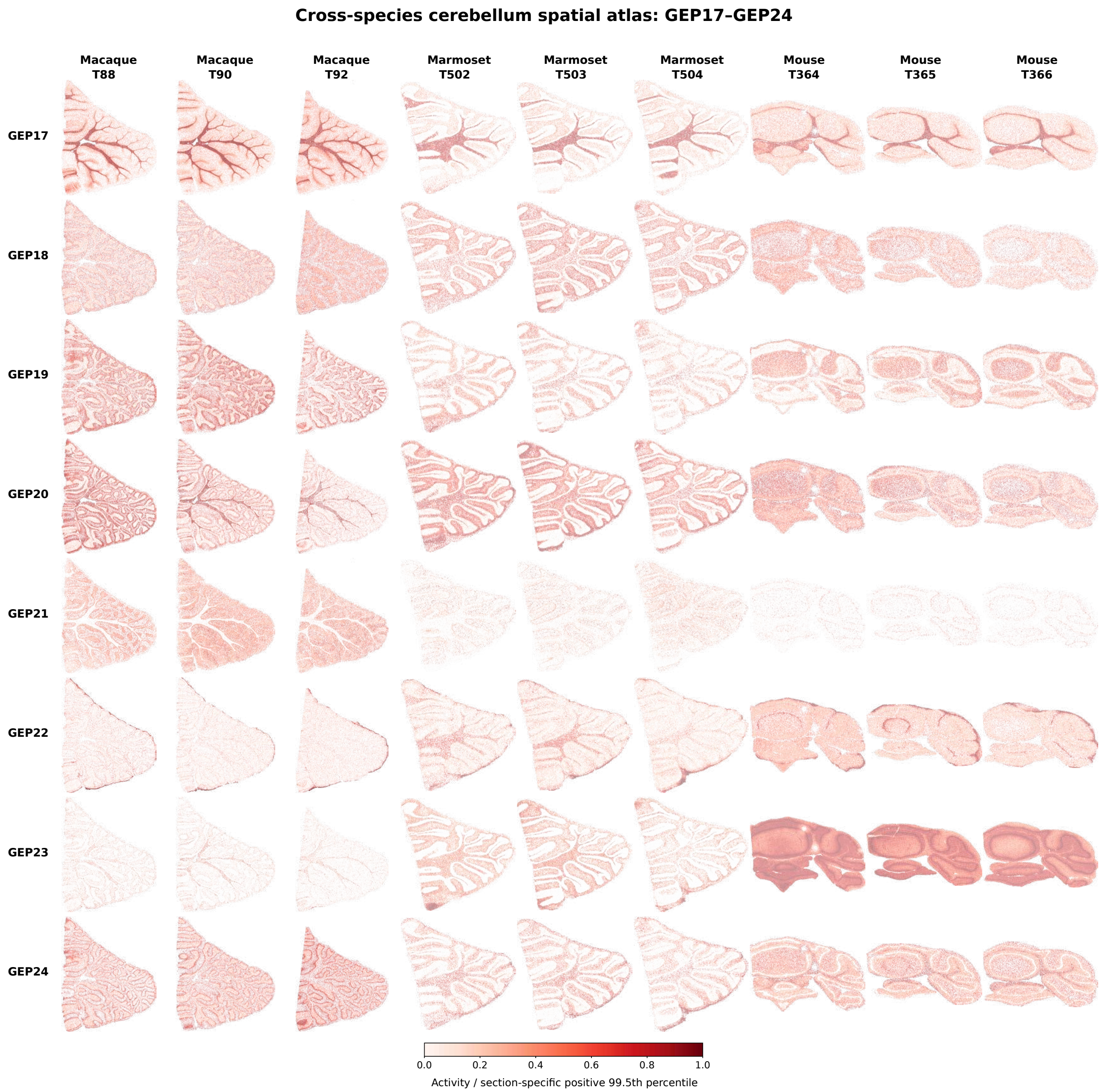


Supplementary Figure S15: **Spatial activity maps for all fitted cerebellar SORT components in selected specimens (continued).** GEP17–GEP24 are shown across the same nine sections using the same rendering procedure.

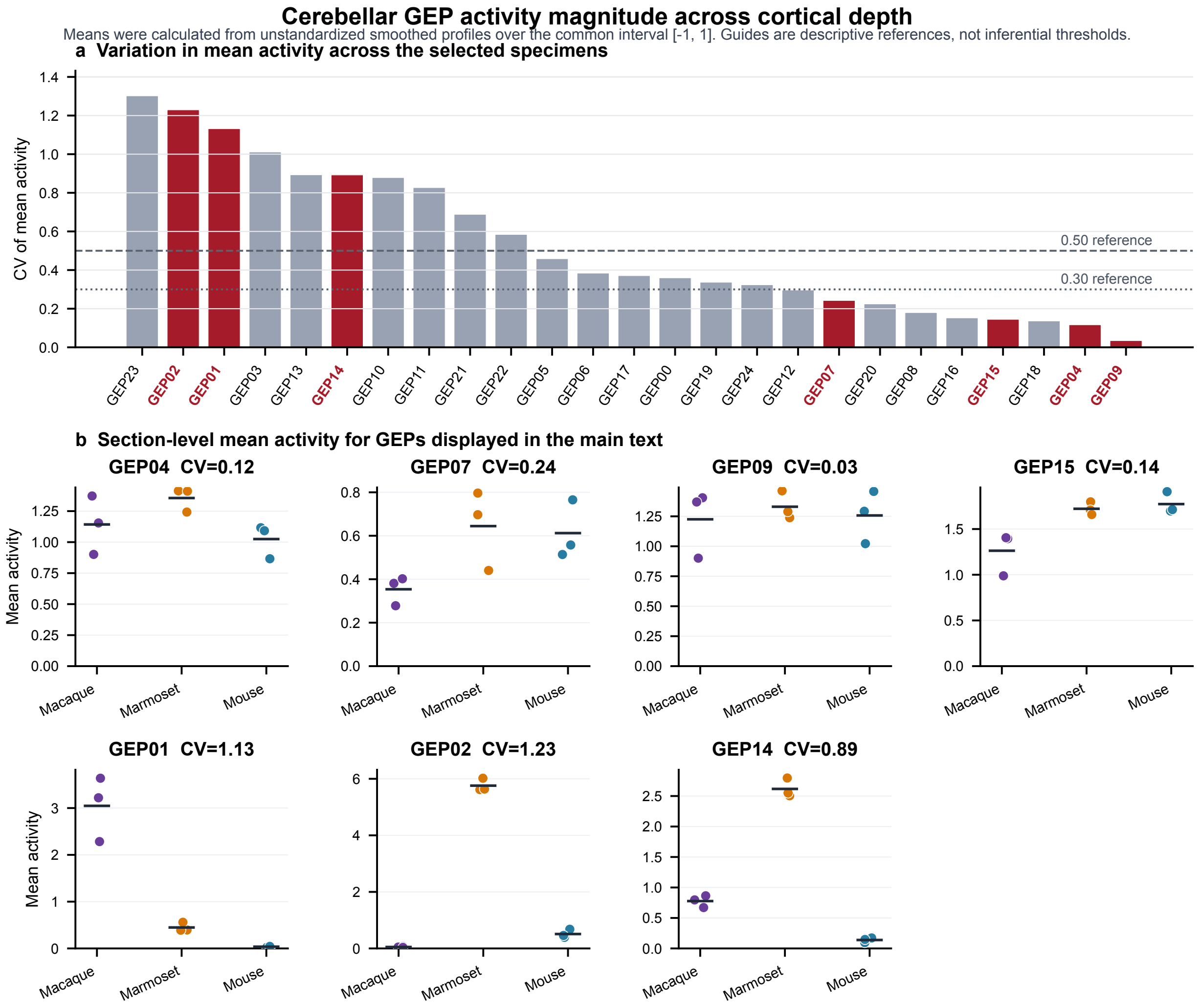


Supplementary Figure S16: **Variation in cerebellar gene expression program activity among selected specimens. (a)** All 25 components ranked by the coefficient of variation (CV) of mean activity. The CV was calculated across the three species means of section activity over the common cortical depth interval $[-1, 1]$. Guides mark CV values of 0.30 and 0.50. Red labels mark the seven signal gene expression programs (GEPs) displayed in main Fig. 5. **(b)** Mean activities for three macaque, three marmoset and three mouse sections show the values underlying the CV for these GEPs. The three sections for each species came from one animal, so the CV summarizes activity magnitude variation among the sampled specimens.

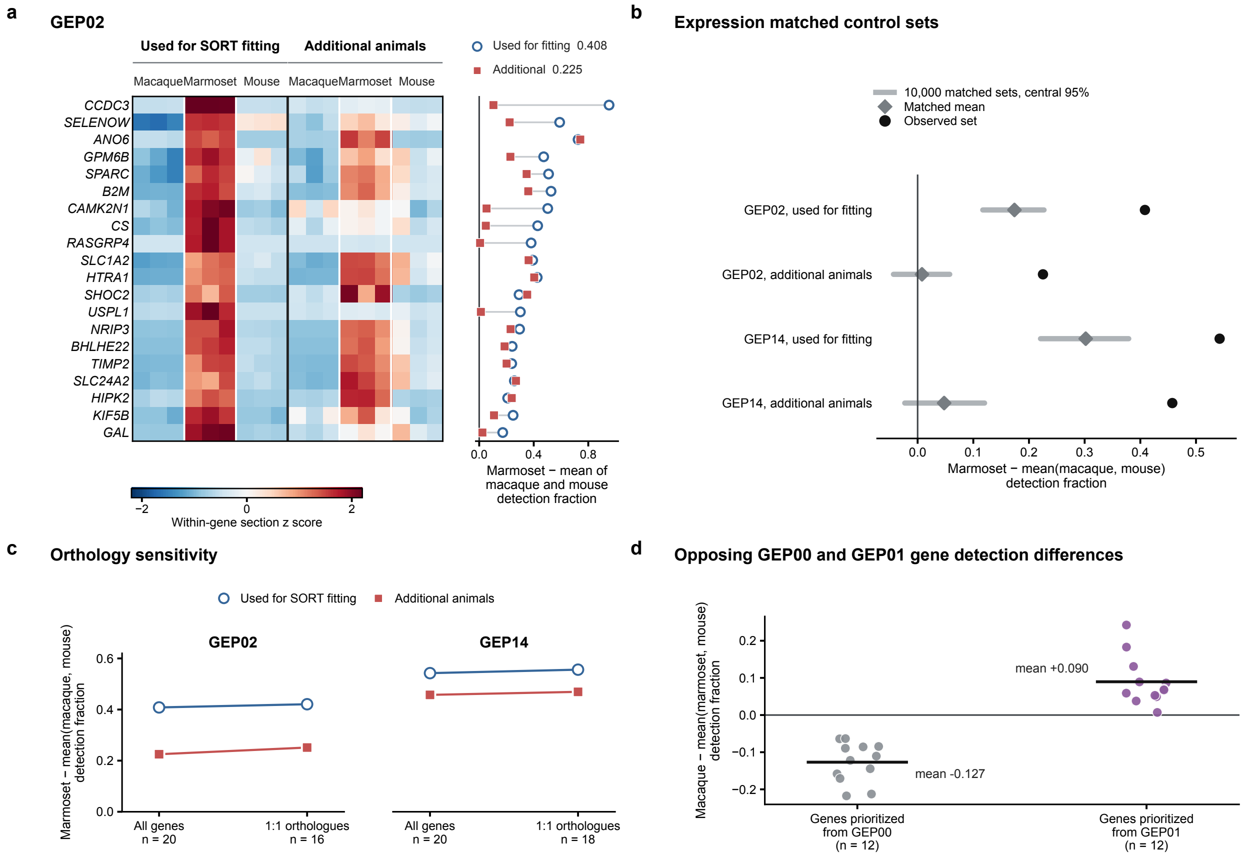


Supplementary Figure S17: **Gene expression support for differences in SORT program deployment among sampled animals.** **(a)** Detection patterns for the 20 prioritized GEP02 genes across three sections from one fitted and one additional animal of each species. Heatmap values are domain detection fractions standardized within each gene. The adjacent plot shows marmoset detection minus the mean detection in macaque and mouse. Open circles and filled squares denote fitted and additional animals. **(b)** Observed gene-set means (black circles) compared with 10,000 control sets matched for gene expression and detection. Gray diamonds and intervals show the matched mean and central 95% interval. **(c)** Mean detection differences for all 20 genes and for confirmed one-to-one orthologues. Lines connect fitted and additional animal sets. **(d)** Detection differences for genes prioritized from GEP00 and GEP01 within their shared granule-layer domain. Points show macaque detection minus the mean in marmoset and mouse. Horizontal bars show group means. For **(a–c)**, corresponding domains in the additional animals were identified by gene-masked projection onto the fixed profiles.

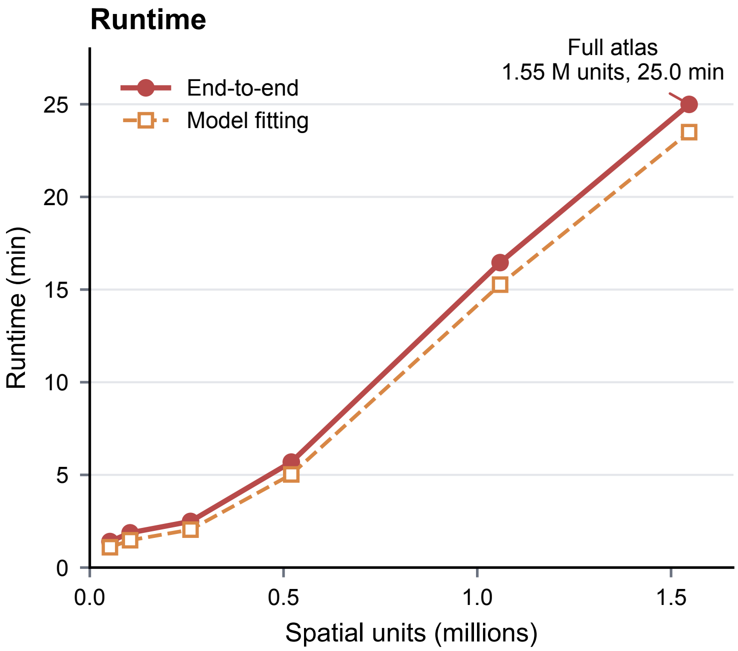


Supplementary Figure S18: **Empirical runtime of SORT on the cross-species cerebellar dataset.** End-to-end wall-clock time and model-fitting time are shown for six balanced inputs containing 51,997–1,546,637 spatial locations. All runs used 4,000 genes, rank $K = 25$ and one NVIDIA A30 graphics processing unit (GPU). Fractional inputs used centered, spatially contiguous crops from one representative section per species. Points show one run per input size under fixed software, hardware and model settings.

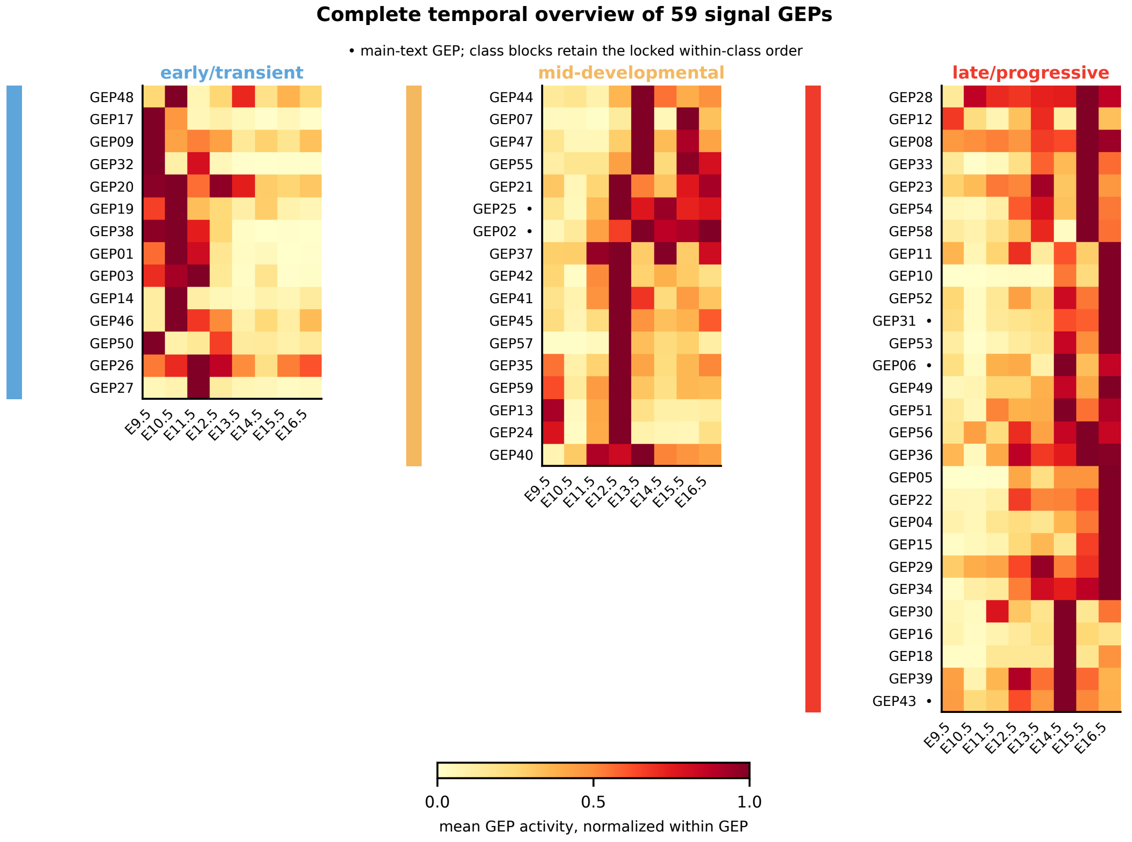


Supplementary Figure S19: **Complete temporal context for all signal programs.** Stage-wise mean activity is shown for gene expression programs (GEPs) GEP01–GEP59 and normalized to the maximum stage mean within each GEP. Rows are arranged in descriptive early/transient, mid-developmental and late/progressive display groups. The 27 GEPs in main Fig. 6b retain the same relative order. Dots mark the five GEPs examined in main Fig. 6. The groups organize the displayed continuum and are not discrete biological categories.

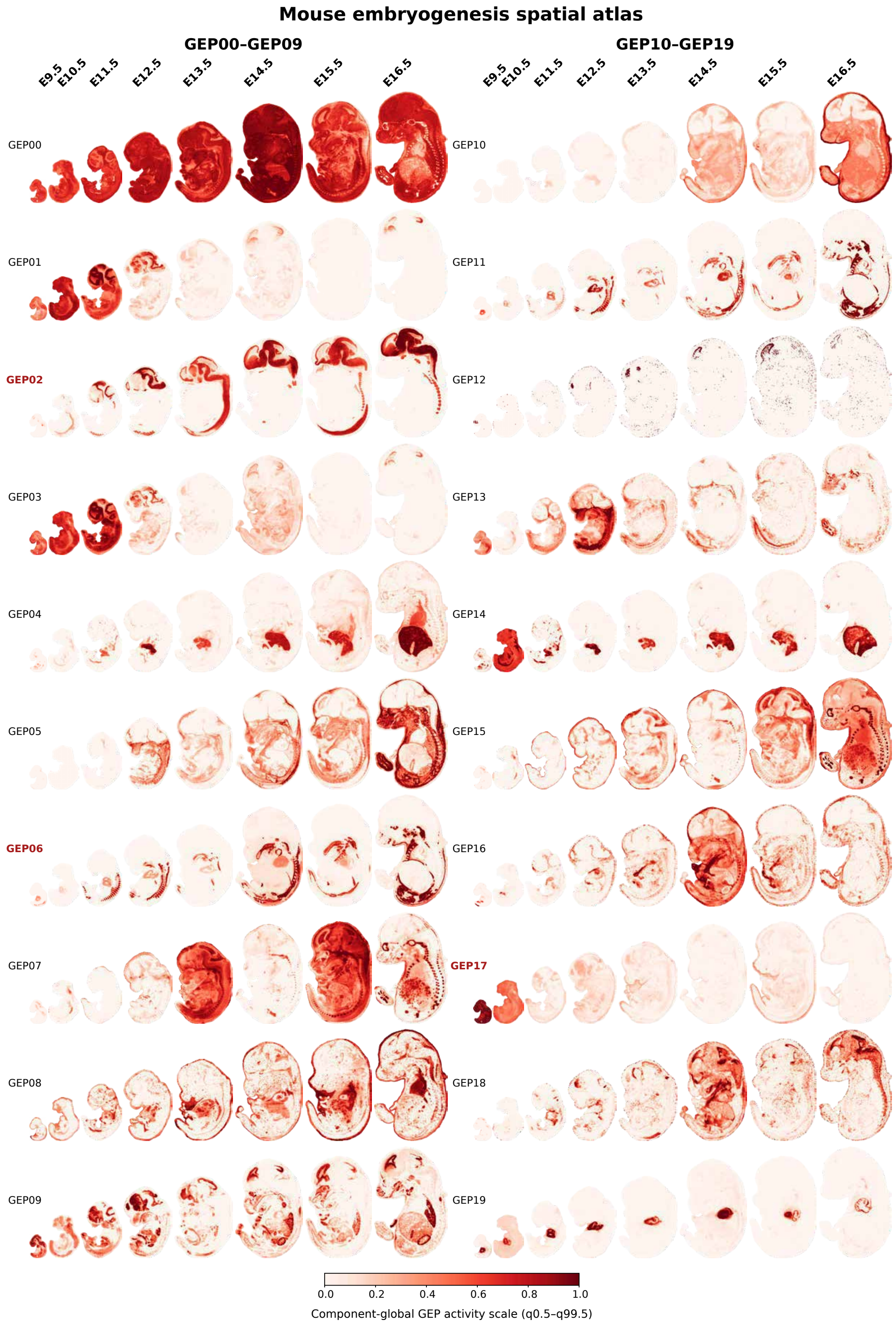


Supplementary Figure S20: **Complete eight-stage spatial catalog of embryogenesis SORT components.** Spatial activity is shown for GEP00 and all 59 signal GEPs across E9.5–E16.5 on three pages. This page shows GEP00–GEP19. Each GEP uses its own color scale across stages.

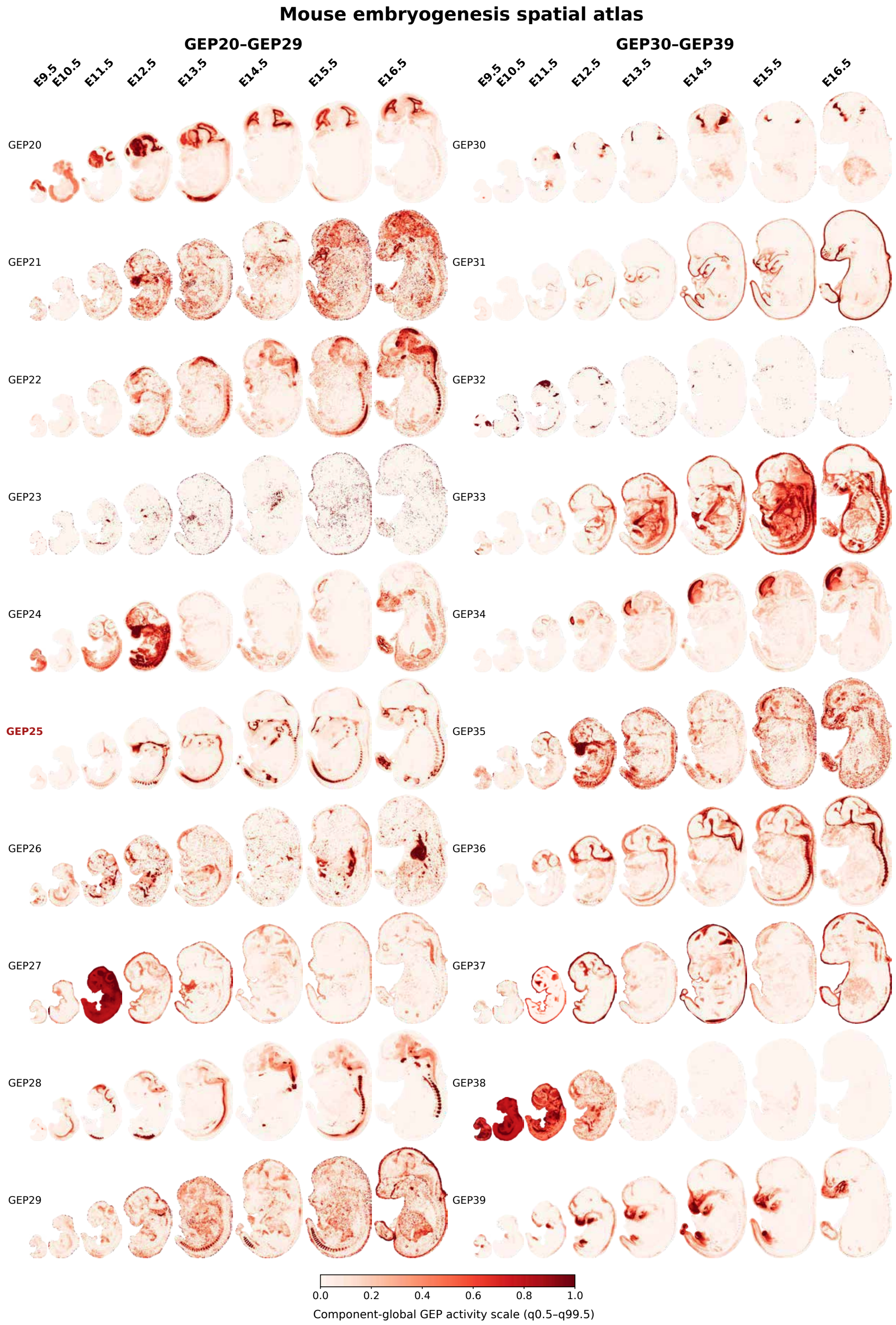


Supplementary Figure S20: **Complete eight-stage spatial catalog of embryogenesis SORT components (continued).** GEP20–GEP39 are shown using the same stage layout and GEP-specific color scales.

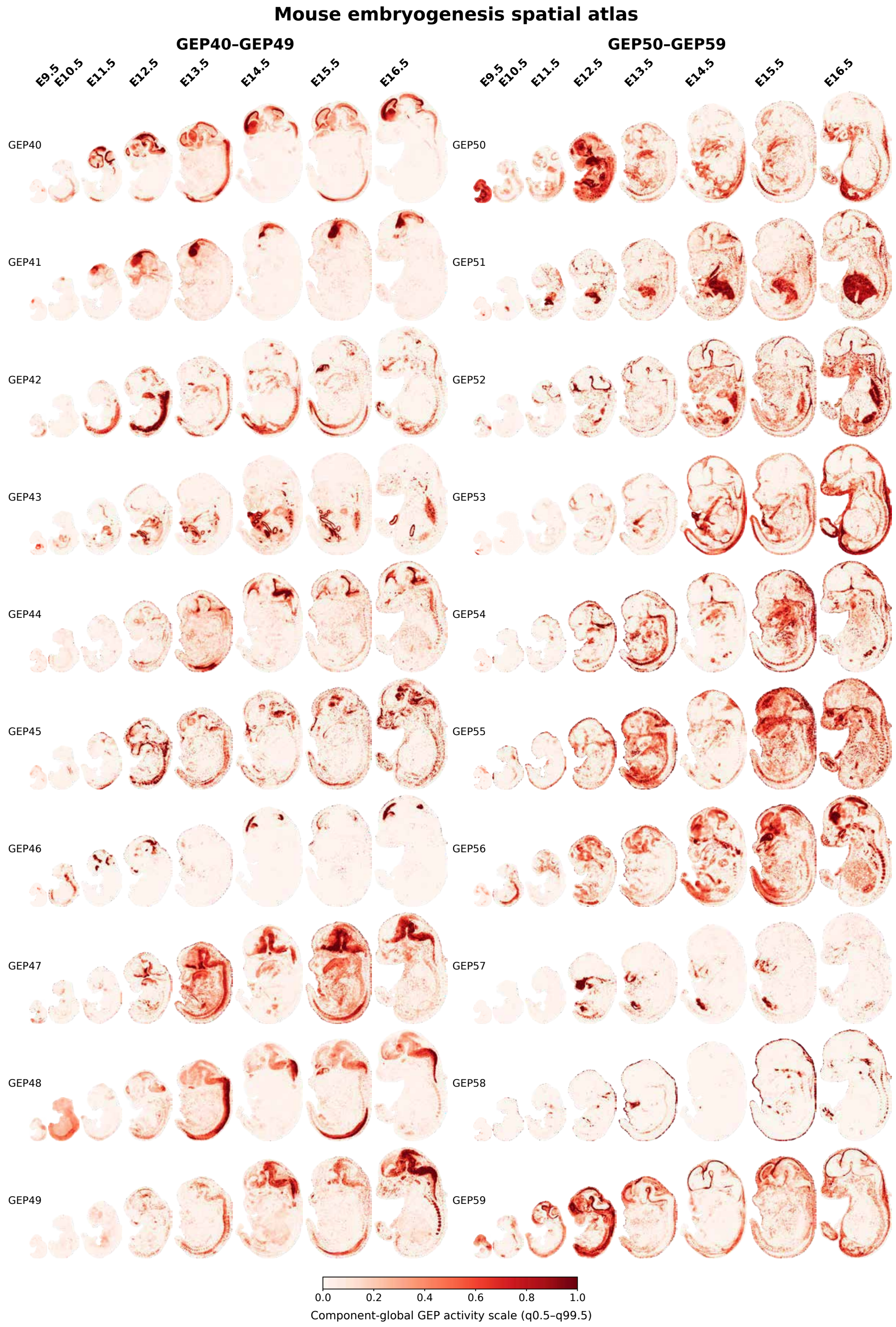


Supplementary Figure S20: **Complete eight-stage spatial catalog of embryogenesis SORT components (continued).** GEP40–GEP59 are shown using the same stage layout and GEP-specific color scales.

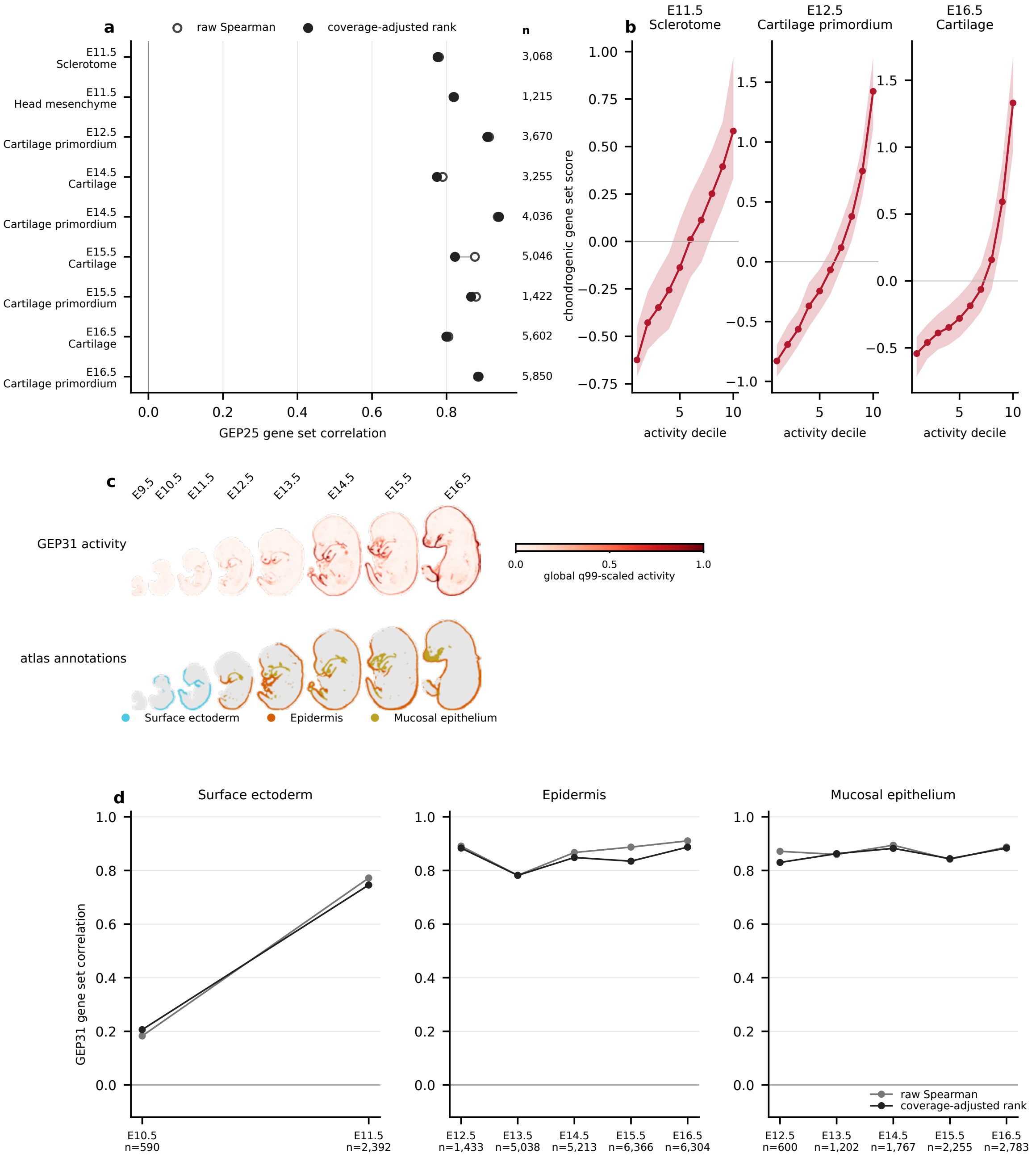


Supplementary Figure S21: **Continuous gene-set support for GEP25 and GEP31.** **(a)** Raw and coverage-adjusted rank correlations between GEP25 activity and a fixed seven-gene chondrogenic gene set across nine anatomical contexts. Coverage denotes the number of detected genes per location, and $n$ is the number of locations. **(b)** Median gene-set score and first to third quartiles across GEP25 activity deciles in representative precursor, primordium and later cartilage contexts. **(c)** Spatial maps of GEP31 activity and the original Surface ectoderm, Epidermis and Mucosal epithelium annotations across eight stages. The three annotations are shown separately. **(d)** Raw and coverage-adjusted rank correlations with the fixed epithelial-associated gene set for every supported stage within each annotation.

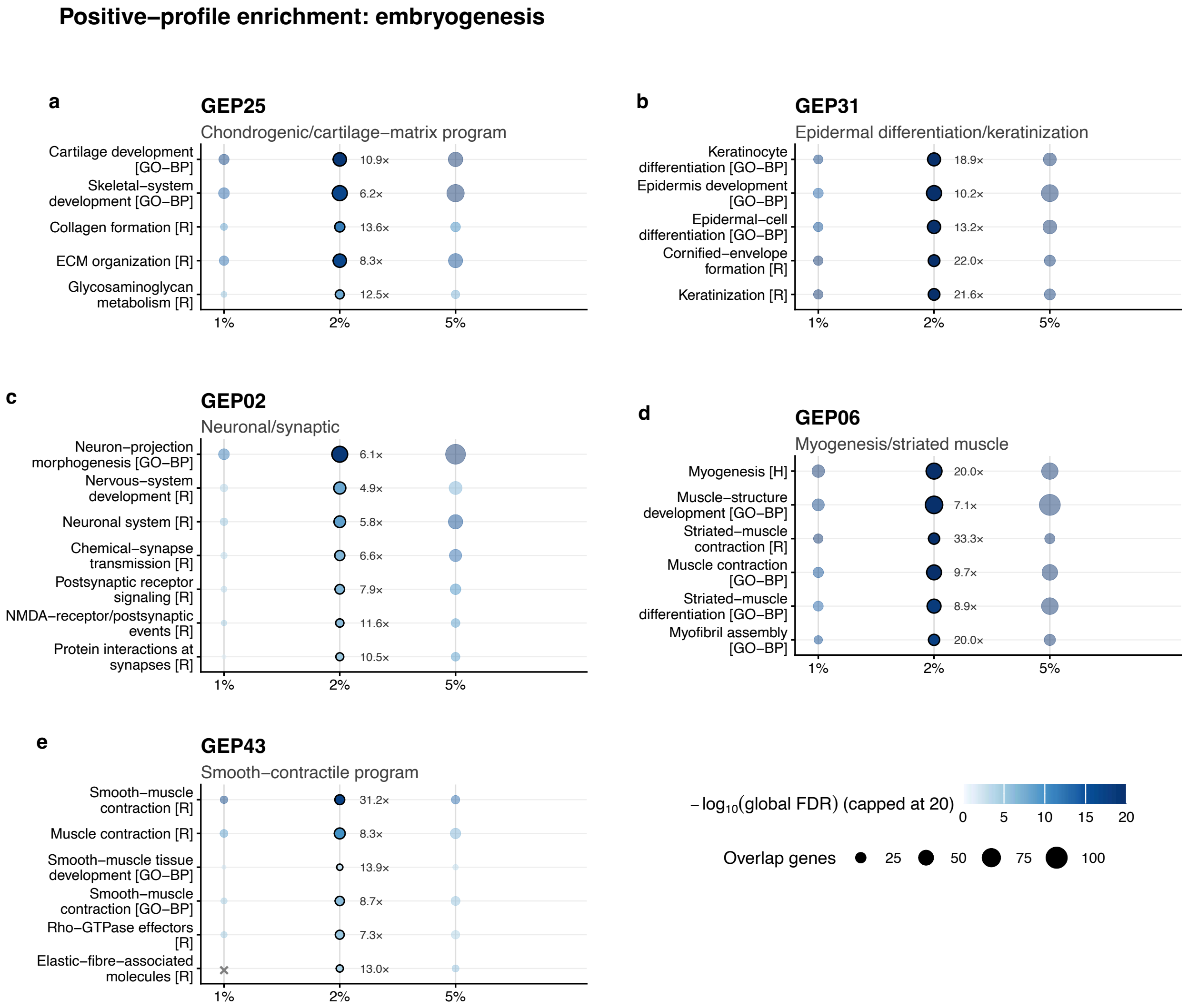


Supplementary Figure S22: **Positive-profile pathway enrichment for selected mouse embryogenesis programs. (a–e)** Results for gene expression programs (GEPs) GEP25, GEP31, GEP02, GEP06 and GEP43. Rows are selected nonredundant Gene Ontology Biological Process terms. Columns use the top 1%, 2% and 5% positive-loading genes in $Q$, with the top 2% as the primary analysis. Outlined points are labeled with fold enrichment (FE). Color represents $-\log_{10}$ global false discovery rate (FDR), capped at 20. Point size represents the number of overlapping genes. Gray crosses mark results with global FDR $\geq 0.05$. Benjamini–Hochberg correction was applied across all signal-GEP–pathway tests within each cutoff.

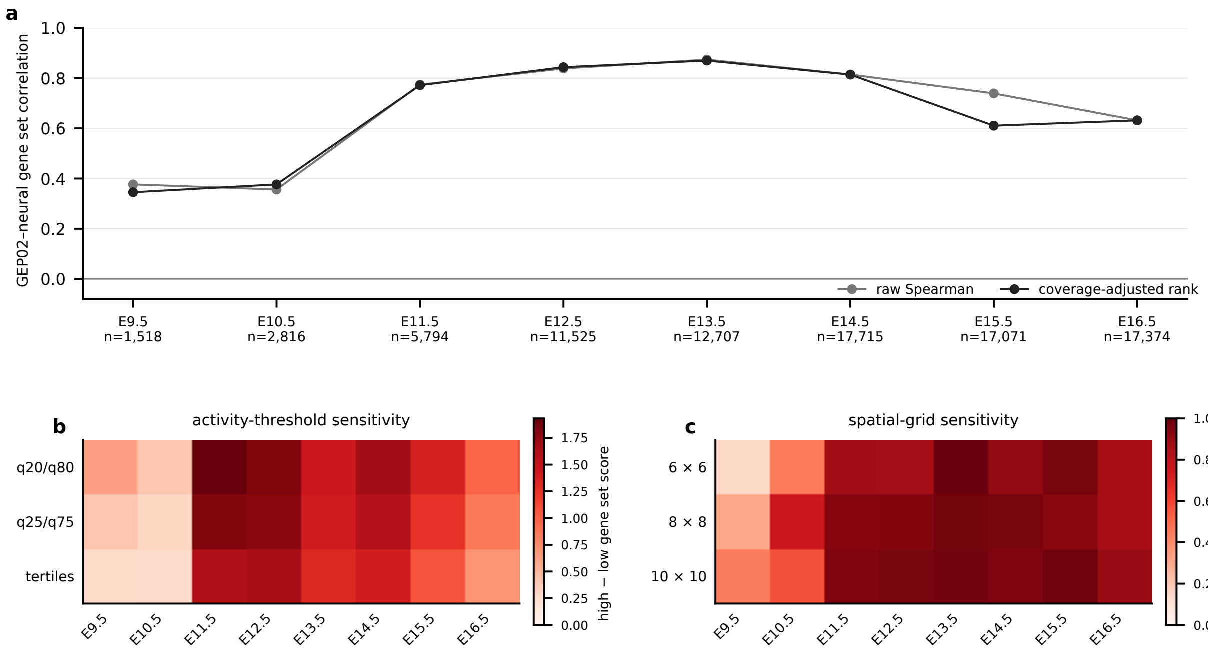


Supplementary Figure S23: **GEP02 gene-set support and sensitivity within Brain. (a)** Raw and coverage-adjusted rank correlations between continuous GEP02 activity and the fixed neural-associated gene-set score at all eight stages. Coverage denotes the number of detected genes per location. **(b)** Difference in median gene-set score between high- and low-activity groups using within-stage 20th/80th percentile, 25th/75th percentile and tertile definitions. **(c)** Spearman correlation between block-median GEP02 activity and gene-set score on $6 \times 6$, $8 \times 8$ and $10 \times 10$ normalized Brain grids. Blocks with fewer than 20 locations are unsupported.

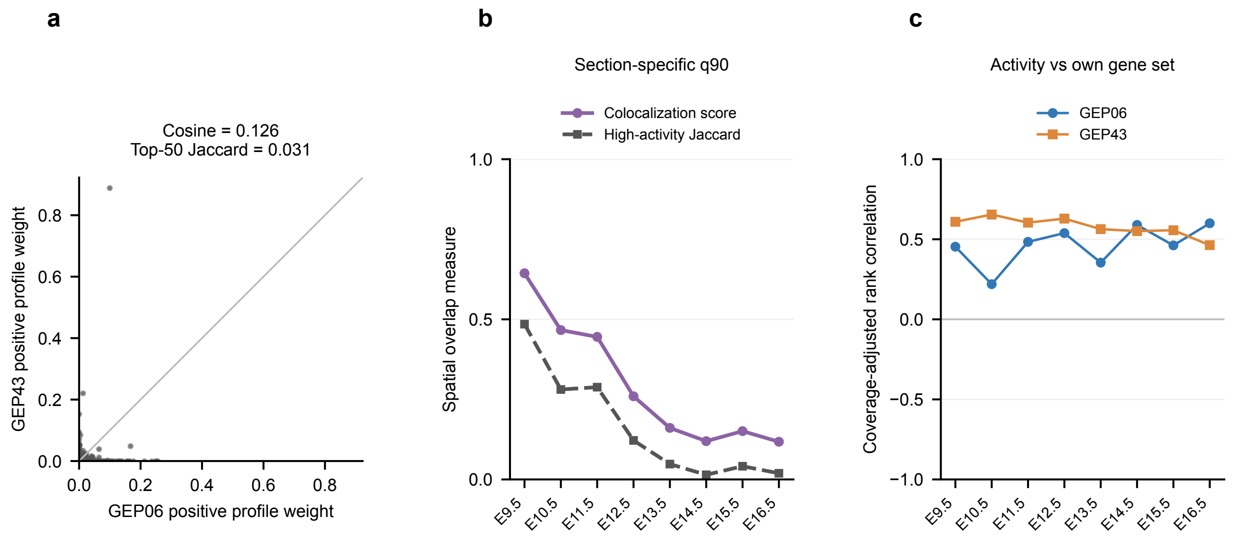


Supplementary Figure S24: **Profile distinction, gene expression concordance and spatial overlap of GEP06 and GEP43 across development.** **(a)** Positive profile weights $Q_k^+ = \max(Q_k, 0)$ for all genes, shown on equal axes with a 1:1 line. Labels report cosine similarity and the Jaccard overlap of the top 50 positive-weight genes. **(b)** Colocalization scores and Jaccard overlap of high-activity locations across stages. High activity was defined separately for each GEP within each whole section using its 90th percentile. **(c)** Coverage-adjusted rank correlations between each GEP's activity and its own fixed gene-set expression score across all locations within each stage. Coverage denotes the number of detected genes per location. Supplementary Table 5 lists the gene sets. Supplementary Note 6 describes the calculations.

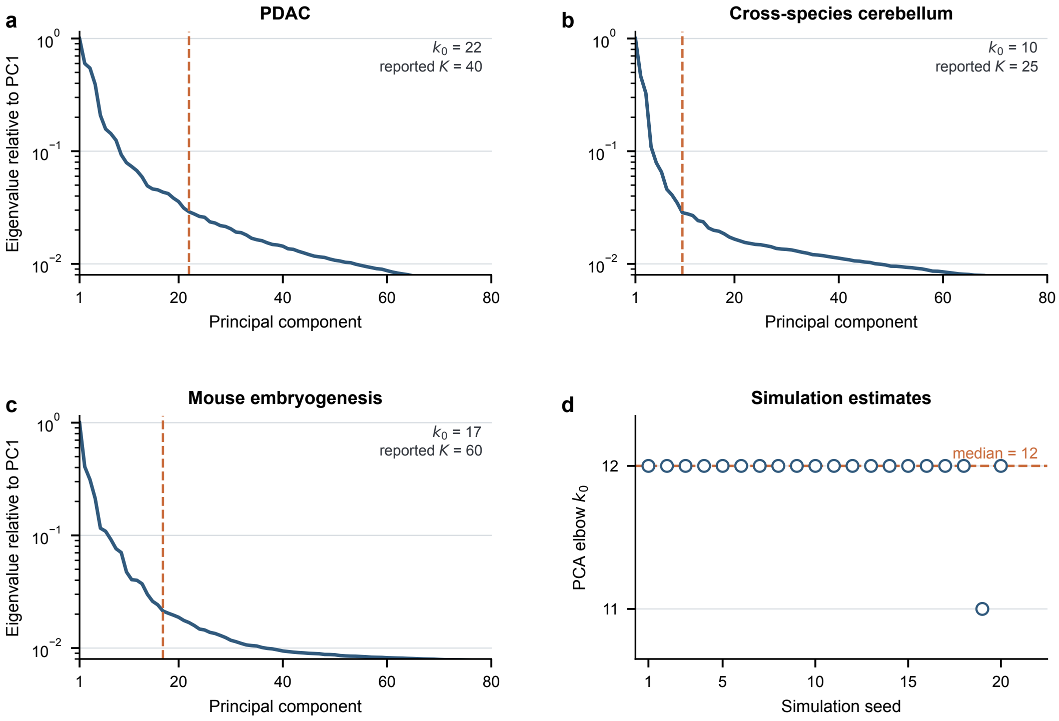


Supplementary Figure S25: **Principal component analysis guidance for the SORT factorization rank. (a–c)** Log eigenvalue scree curves from balanced sketches of the PDAC, cerebellar and mouse embryogenesis datasets, using the genes included in the corresponding SORT fits. Orange dashed lines mark elbow $k_0$, defined by the greatest distance from the line joining the first and eightieth log eigenvalues. The reported SORT rank is shown for comparison. **(d)** Estimates of $k_0$ across the 20 simulation datasets. Points represent simulation seeds, and the dashed line marks the median.

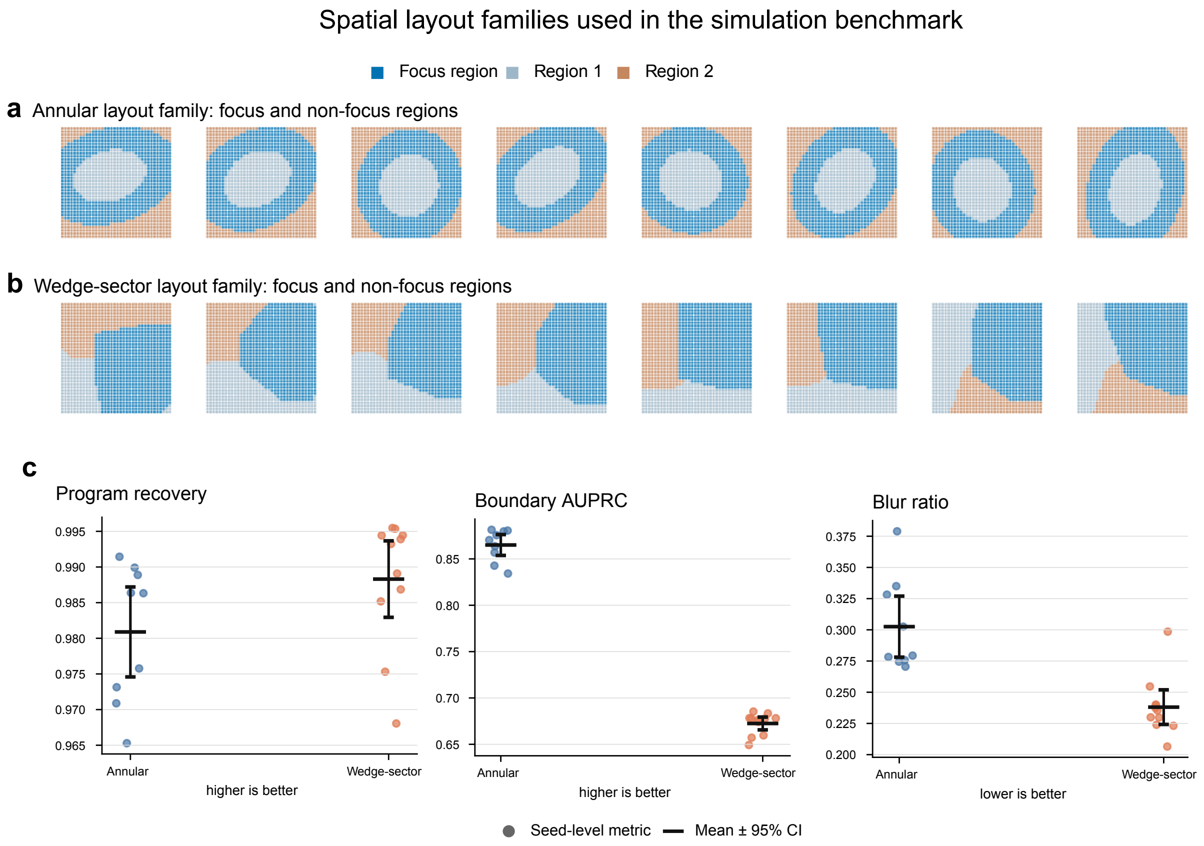


Supplementary Figure S26: **Spatial layouts used in the simulation benchmark. (a,b)** Focus and two non-focus regions for the annular and wedge-sector layout families across 20 simulation seeds. Each row shows all eight sections. **(c)** SORT recovery and spatial fidelity for annular ($n = 9$ seeds) and wedge-sector layouts ($n = 11$ seeds). Black summaries show the mean and approximate 95% confidence interval calculated as 1.96 standard errors. Higher boundary area under the precision–recall curve (AUPRC) and lower blur ratio indicate better boundary recovery.

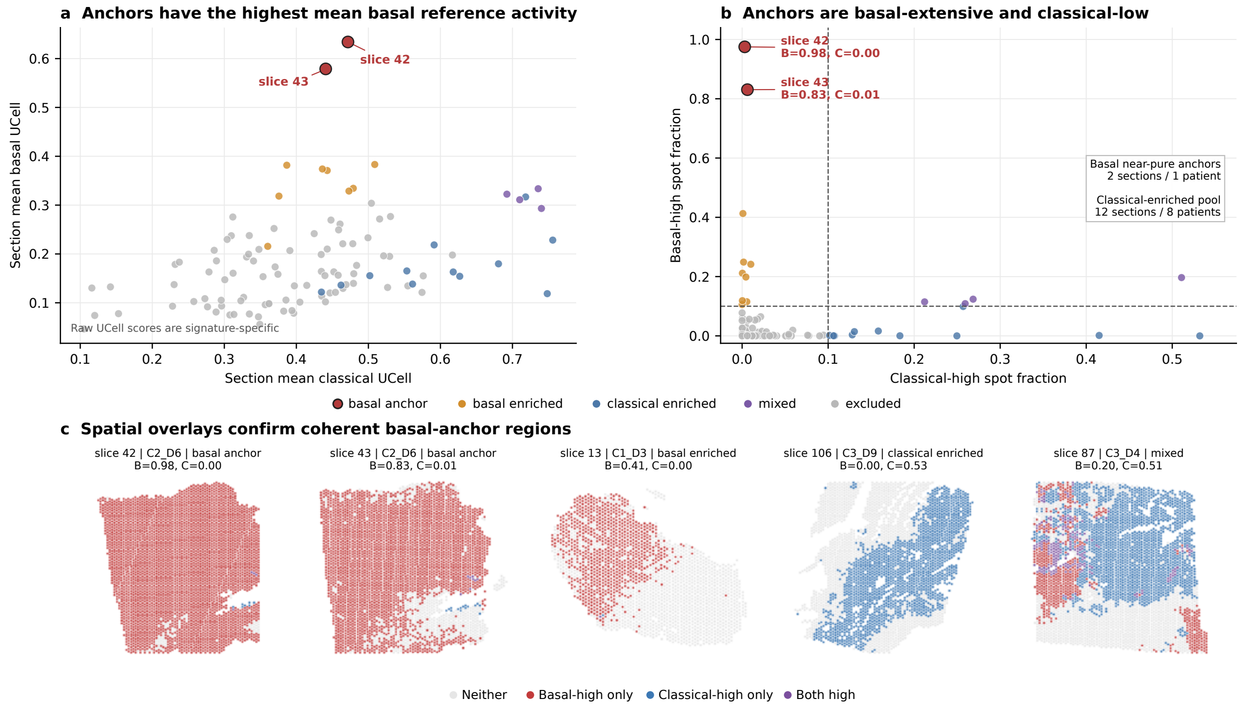


Supplementary Figure S27: **Reference-score diagnostics for the pancreatic ductal adenocarcinoma subtype benchmark.** **(a)** Mean basal and classical UCell scores across all 108 sections. Points represent sections, and colors indicate sampling strata. **(b)** Fractions of basal-high and classical-high locations. Dashed lines mark the section-eligibility threshold, and the inset shows the basal anchors and classical-enriched section pool. **(c)** Spatial overlays for the anchors and representative basal-enriched, classical-enriched and mixed sections. Together, these diagnostics support construction of subsets with evaluable basal and classical reference signals before model fitting.

Supplementary Table 1: Representative enrichment terms for all signal GEPs in PDAC.

| GEP | Top ten positive-weight genes | Representative enriched terms |
|---|---|---|
| **GEP01** | *KRT17*, *LAMC2*, *LCN2*, *TRIM29*, *MMP11*, *KLK6*, *KRT16*, *CLDN4*, *KRT7*, *KRT8* | Extracellular matrix organization$^{R}$<br>Keratinization$^{R}$<br>type I hemidesmosome assembly$^{R}$<br>Coagulation$^{H}$ |
| **GEP02** | *PNLIP*, *CELA3A*, *PRSS1*, *CLPS*, *CPA1*, *CPB1*, *CTRC*, *CEL*, *AMY2A*, *REG1A* | Digestion and absorption$^{R}$<br>Coagulation$^{H}$<br>Epithelial mesenchymal transition$^{H}$ |
| GEP03 | *C7*, *CCN1*, *C3*, *FOS*, *CCDC80*, *EGR1*, *FBLN1*, *CCN2*, *SELENOP*, *PTGDS* | Extracellular matrix organization$^{R}$<br>TNF$\alpha$ signaling via NF-$\kappa$B$^{H}$<br>Apoptosis$^{H}$<br>Hypoxia$^{H}$ |
| **GEP04** | *COL1A1*, *SPARC*, *POSTN*, *COL5A1*, *COL1A2*, *COL3A1*, *FN1*, *ACTA2*, *SFRP2*, *VCAN* | Extracellular matrix organization$^{R}$<br>Collagen formation$^{R}$<br>Integrin cell surface interactions$^{R}$<br>Epithelial mesenchymal transition$^{H}$ |
| GEP05 | *UBD*, *IDO1*, *HLA-A*, *IFI6*, *IGHG1*, *WARS*, *IGLV3-1*, *S100A14*, *PLAAT4*, *C3* | Interferon gamma response$^{H}$<br>Interferon alpha response$^{H}$<br>Antigen presentation folding assembly and peptide loading of class i MHC$^{R}$<br>MYC targets v1$^{H}$ |
| GEP06 | *IGHG1*, *IGLC2*, *IGKV1D-12*, *IGLC3*, *IGHV3-30*, *IGKC*, *IGLC1*, *IGHA1*, *IGHG2*, *IGHV3-35* | Complement cascade$^{R}$<br>Binding and uptake of ligands by scavenger receptors$^{R}$<br>Adaptive immune system$^{R}$<br>FcGR activation$^{R}$ |
| GEP07 | *DES*, *ACTG2*, *MYH11*, *SYNM*, *CNN1*, *TPM2*, *MYLK*, *SYNPO2*, *GREM1*, *TPM1* | Myogenesis$^{H}$<br>Smooth muscle contraction$^{R}$<br>Cell extracellular matrix interactions$^{R}$<br>Cellular response to heat stress$^{R}$ |
| **GEP08** | *TFF1*, *CEACAM6*, *LGALS4*, *GPX2*, *CTSE*, *TFF3*, *MUC1*, *TSPAN8*, *MUC5AC*, *SPINK1* | O-linked glycosylation of mucins$^{R}$<br>Protein secretion$^{H}$<br>Neutrophil degranulation$^{R}$<br>Estrogen response late$^{H}$ |
| GEP09 | *SERF1A*, *COL1A2*, *COL3A1*, *COL1A1*, *UBC*, *TMSB10*, *ACTG1*, *S100A6*, *TPM2*, *FN1* | ECM proteoglycans$^{R}$<br>Collagen degradation$^{R}$<br>Neutrophil degranulation$^{R}$<br>Hypoxia$^{H}$ |
| GEP10 | *IFI6*, *RAN*, *ARGLU1*, *SLC25A5*, *TPM2*, *CD59*, *ANP32E*, *ADAM9*, *S100A4*, *LUC7L3* | Epithelial mesenchymal transition$^{H}$<br>Death receptor signalling$^{R}$<br>Interferon signaling$^{R}$<br>MYC targets v1$^{H}$ |
| **GEP11** | *CCL21*, *SERF1A*, *EEF1A1*, *TRBC2*, *NOP53*, *PTPRC*, *CXCR4*, *PABPC1*, *IL7R*, *DDX5* | Cytokine signaling in immune system$^{R}$<br>Adaptive immune system$^{R}$<br>Translation$^{R}$<br>MYC targets v1$^{H}$ |
| GEP12 | *S100A9*, *SLPI*, *HLA-C*, *S100A8*, *GLUL*, *S100A4*, *ADIRF*, *NACA*, *KRT5*, *PABPC1* | Neutrophil degranulation$^{R}$<br>Translation$^{R}$<br>Epithelial mesenchymal transition$^{H}$ |
| GEP13 | *SERF1A*, *IGKC*, *TMSB10*, *IGHA1*, *EEF2*, *IL32*, *ACTG1*, *HLA-B*, *B2M*, *ENO1* | Epithelial mesenchymal transition$^{H}$<br>Hemostasis$^{R}$<br>Cytokine signaling in immune system$^{R}$<br>TNF$\alpha$ signaling via NF-$\kappa$B$^{H}$ |

Continued on next page

**Supplementary Table 1 (continued)**

| GEP | Top ten positive-weight genes | Representative enriched terms |
|---|---|---|
| **GEP14** | *MCAM*, *ADIRF*, *A2M*, *MYH11*, *ACTA2*, *MUSTN1*, *RGS5*, *TAGLN*, *EPAS1*, *MYL9* | Myogenesis[H]<br>Smooth muscle contraction[R]<br>Extracellular matrix organization[R] |
| GEP15 | *C11ORF96*, *G0S2*, *NR4A3*, *FOSB*, *ADAMTS4*, *NAMPT*, *NR4A1*, *KDM6B*, *LMNA*, *TIMP1* | TNF$\alpha$ signaling via NF-$\kappa$B[H]<br>Extracellular matrix organization[R]<br>Hypoxia[H]<br>Apoptosis[H] |
| **GEP16** | *S100A2*, *PAEP*, *TGM2*, *DCBLD2*, *TCN1*, *LAMA3*, *ITGB4*, *HMGA2*, *MT2A*, *TNFRSF6B* | Epithelial mesenchymal transition[H]<br>type I hemidesmosome assembly[R]<br>Keratinization[R]<br>Hypoxia[H]<br>Glycolysis[H] |
| GEP17 | *PLA2G2A*, *RARRES1*, *CXCL17*, *WFDC2*, *KRT17*, *KRT6B*, *CBLC*, *C3*, *ETS2*, *SLPI* | Estrogen response late[H]<br>Peroxisome[H]<br>Cholesterol homeostasis[H]<br>Interferon signaling[R] |
| GEP18 | *CTRL*, *CELA2B*, *KLK1*, *AMY2B*, *SYCN*, *CELA2A*, *GATM*, *PNLIPRP1*, *PDIA2*, *PRSS3* | Translation[R]<br>Oxidative phosphorylation[H]<br>Digestion[R]<br>Metallothioneins bind metals[R] |
| GEP19 | *PPY*, *HSPA1B*, *HSPA1A*, *AMY2A*, *DNAJB1*, *CHGB*, *PNLIP*, *HSPH1*, *HSP90AA1*, *HSPE1* | Cellular response to heat stress[R]<br>HSF1 dependent transactivation[R]<br>MYC targets v1[H] |
| GEP20 | *APOE*, *LYZ*, *APOC1*, *CTSD*, *TYROBP*, *C1QB*, *IFI30*, *FTL*, *CD68*, *C1QC* | Neutrophil degranulation[R]<br>Complement[H]<br>MHC class ii antigen presentation[R] |
| **GEP21** | *SPP1*, *MT2A*, *ALDOA*, *DDIT4*, *ENO1*, *MT1B*, *SOD2*, *FN1*, *MIF*, *VEGFA* | Hypoxia[H]<br>mTORC1 signaling[H]<br>Glycolysis[H]<br>Cellular response to chemical stress[R] |
| **GEP22** | *HLA-DRB1*, *LYZ*, *HLA-DRB5*, *IGHG4*, *MTRNR2L12*, *CD74*, *SERF1B*, *HLA-DQA1*, *MT1G*, *SLC9A3* | Metallothioneins bind metals[R]<br>Interferon signaling[R]<br>Antigen presentation folding assembly and peptide loading of class i MHC[R] |
| GEP23 | *IGHG1*, *CXCL14*, *GABRP*, *IGHA1*, *CST1*, *IGLC1*, *PTGDS*, *COMP*, *TGM2*, *IGFBP7* | Epithelial mesenchymal transition[H]<br>Myogenesis[H]<br>Apoptosis[H]<br>Fcgamma receptor FcGR dependent phagocytosis[R] |
| GEP24 | *APOD*, *IGFBP5*, *SPON2*, *IGFBP3*, *TSHZ2*, *F2R*, *ANO1*, *C7*, *GEM*, *LAMA2* | Extracellular matrix organization[R]<br>Laminin interactions[R]<br>Signaling by receptor tyrosine kinases[R] |
| GEP25 | *ITGA3*, *KRT17*, *IGHG1*, *HLA-DRB1*, *DEFB1*, *SPINK1*, *HLA-DRB5*, *ANGPTL4*, *SERPINE1*, *MMP7* | Epithelial mesenchymal transition[H]<br>Hypoxia[H]<br>TNF$\alpha$ signaling via NF-$\kappa$B[H]<br>Coagulation[H] |
| GEP26 | *SERF1A*, *IGKC*, *SERF1B*, *ACTB*, *TMSB4X*, *IGHG4*, *EEF1A1*, *TMSB10*, *B2M*, *S100A6* | Epithelial mesenchymal transition[H]<br>Extracellular matrix organization[R]<br>Cytokine signaling in immune system[R]<br>Hemostasis[R] |

Continued on next page

**Supplementary Table 1 (continued)**

| GEP | Top ten positive-weight genes | Representative enriched terms |
|---|---|---|
| GEP27 | *APOA1*, *ANPEP*, *ALDOB*, *RBP2*, *FABP1*, *APOA4*, *SLC5A1*, *CDHR5*, *APOB*, *TMPRSS15* | Biological oxidations[R]<br>Fatty acid metabolism[H]<br>Xenobiotic metabolism[H]<br>Gluconeogenesis[R] |
| GEP28 | *HIST1H1B*, *HIST1H3D*, *HIST1H2AG*, *HIST1H2AJ*, *HIST1H4C*, *HIST1H2BG*, *HIST1H1C*, *HIST1H4D*, *CLU*, *HIST1H3H* | Signaling by the b cell receptor BCR[R]<br>Tcr signaling[R]<br>E2F targets[H]<br>Translation[R] |
| **GEP29** | *LCN2*, *PIGR*, *FXYD2*, *ANXA4*, *CRISP3*, *MUC5B*, *CLU*, *SPP1*, *SLC4A4*, *CRP* | Complement cascade[R]<br>Coagulation[H]<br>Xenobiotic metabolism[H] |
| **GEP30** | *CTRB1*, *CTRB2*, *PRSS2*, *REG1B*, *REG3A*, *MT2A*, *CELA3A*, *IFI6*, *REG3G*, *REG1A* | Interferon alpha response[H]<br>Interferon gamma response[H]<br>Translation[R]<br>Metallothioneins bind metals[R] |
| GEP31 | *LYZ*, *TFF2*, *MUC6*, *PGC*, *GOLM1*, *C6ORF58*, *RNASE1*, *REG1A*, *GP2*, *FMOD* | Antimicrobial peptides[R]<br>Digestion[R]<br>Protein secretion[H]<br>Coagulation[H] |
| **GEP32** | *OLFM4*, *HLA-A*, *KRT17*, *TFF1*, *CXCL14*, *PERP*, *TFF3*, *KRT5*, *HSPB1*, *SLC2A1* | Hypoxia[H]<br>Glycolysis[H]<br>Keratinization[R]<br>Extracellular matrix organization[R] |
| GEP33 | *HLA-A*, *CEACAM6*, *INS*, *IFI27*, *MTRNR2L12*, *SLC34A2*, *PTP4A1*, *COL10A1*, *PFDN5*, *EGR1* | Interferon signaling[R]<br>Complement cascade[R]<br>Neutrophil degranulation[R] |
| GEP34 | *INS*, *GCG*, *TTR*, *CHGB*, *SST*, *SLC30A8*, *CPE*, *SCG5*, *CHGA*, *ERO1B* | Pancreas beta cells[H]<br>Insulin processing[R]<br>Vesicle mediated transport[R]<br>Incretin synthesis secretion and inactivation[R] |
| GEP35 | *REG3A*, *PPY*, *OLFM4*, *PRSS2*, *REG1B*, *HLA-DRB1*, *EGR1*, *IGLC2*, *ZFP36*, *C7* | TNF$\alpha$ signaling via NF-$\kappa$B[H]<br>Epithelial mesenchymal transition[H]<br>Hypoxia[H]<br>Angiogenesis[H] |
| GEP36 | *MMP11*, *COL11A1*, *COL12A1*, *KRT17*, *KRT16*, *CCN1*, *SPP1*, *GREM1*, *ITGA11*, *MMP2* | Extracellular matrix organization[R]<br>Assembly of collagen fibrils and other multimeric structures[R]<br>Integrin cell surface interactions[R] |
| GEP37 | *HLA-C*, *LCN2*, *IFI6*, *CXCL14*, *RNASE1*, *ISG15*, *TCN1*, *PLEC*, *KRT7*, *IFI27* | Epithelial mesenchymal transition[H]<br>Apoptosis[H]<br>Interferon signaling[R]<br>Hypoxia[H] |
| GEP38 | *HBA2*, *HBB*, *HBD*, *HBA1*, *GLUL*, *MT2A*, *IGHA1*, *FOSB*, *APOD*, *ZFP36* | TNF$\alpha$ signaling via NF-$\kappa$B[H]<br>Hypoxia[H]<br>Apoptosis[H] |
| GEP39 | *HMGA2*, *PRSS2*, *YBX3*, *FKBP4*, *ITGB8*, *GAPDH*, *LDHB*, *ANXA2*, *TAGLN2*, *HIST1H1B* | MYC targets v1[H]<br>Cellular response to heat stress[R]<br>Oxidative phosphorylation[H] |

All 39 signal GEPs are listed; background GEP00 is excluded. Bold identifiers denote GEPs discussed in the corresponding Results section. The ten genes with the largest positive $Q$ weights are shown in descending order using standardized gene symbols. Enrichment used the larger set of positive-weight genes in the top 2% of the complete profile ranking and MSigDB v7.5.1 Hallmark and Reactome. Three to five significant terms were selected to summarize enriched themes and reduce repetitive entries. All displayed terms have Benjamini–Hochberg FDR < 0.05, corrected across all signal GEPs and tested terms within each collection. [H]Hallmark; [R]Reactome. Significant enrichment results and FDR values are provided in Supplementary Data 1.

Supplementary Table 2: External PDAC cohorts used for bulk projection and survival analysis.

| Cohort | Patients | Events | Available Cox covariates |
|---|---|---|---|
| TCGA | 102 | 60 | Age, sex, stage |
| CPTAC | 87 | 47 | Age, sex, stage |
| QCMG | 70 | 47 | Age, sex, stage |
| PanCuRx | 132 | 106 | Age, sex, stage |

All cohorts used overall survival. For each patient's bulk sample, normalized expression values were averaged across genes from the fixed GEP gene set that were measured in that cohort. The resulting GEP-specific scores were z-scored across patients within the same cohort. Covariate-adjusted cohort models used the listed variables when available. Cohort-specific log hazard ratios and standard errors were combined by fixed-effect inverse-variance meta-analysis, with Benjamini–Hochberg correction across all 39 signal GEPs. The pooled Kaplan–Meier display included 391 patients and 260 events, with 195 patients above and 196 at or below the global median score.

**Supplementary Table 2 (continued). Survival associations for all 39 pancreatic ductal adenocarcinoma signal programs.**

| Rank | GEP | Meta-analysis HR (95% CI) | $P$ | FDR | $I^2$ (%) | Same direction |
|---|---|---|---|---|---|---|
| 1 | GEP21 | 1.482 (1.295–1.697) | $1.21 \times 10^{-8}$ | $4.74 \times 10^{-7}$ | 25.2 | Yes |
| 2 | GEP08 | 0.665 (0.571–0.774) | $1.63 \times 10^{-7}$ | $3.17 \times 10^{-6}$ | 0.0 | Yes |
| 3 | GEP22 | 0.689 (0.593–0.800) | $1.13 \times 10^{-6}$ | $1.47 \times 10^{-5}$ | 0.0 | Yes |
| 4 | GEP16 | 1.408 (1.218–1.627) | $3.74 \times 10^{-6}$ | $3.65 \times 10^{-5}$ | 47.1 | Yes |
| 5 | GEP31 | 0.734 (0.638–0.844) | $1.38 \times 10^{-5}$ | $1.08 \times 10^{-4}$ | 0.0 | Yes |
| 6 | GEP09 | 1.351 (1.174–1.555) | $2.71 \times 10^{-5}$ | $1.68 \times 10^{-4}$ | 44.8 | Yes |
| 7 | GEP39 | 1.303 (1.151–1.476) | $3.01 \times 10^{-5}$ | $1.68 \times 10^{-4}$ | 75.5 | Yes |
| 8 | GEP01 | 1.328 (1.146–1.538) | $1.54 \times 10^{-4}$ | $5.69 \times 10^{-4}$ | 0.0 | Yes |
| 9 | GEP04 | 1.315 (1.141–1.516) | $1.61 \times 10^{-4}$ | $5.69 \times 10^{-4}$ | 43.8 | Yes |
| 10 | GEP12 | 1.359 (1.161–1.590) | $1.34 \times 10^{-4}$ | $5.69 \times 10^{-4}$ | 23.2 | Yes |
| 11 | GEP36 | 1.304 (1.136–1.495) | $1.53 \times 10^{-4}$ | $5.69 \times 10^{-4}$ | 16.6 | Yes |
| 12 | GEP15 | 1.272 (1.119–1.446) | $2.40 \times 10^{-4}$ | $7.81 \times 10^{-4}$ | 56.8 | No |
| 13 | GEP33 | 1.301 (1.129–1.498) | $2.71 \times 10^{-4}$ | $8.13 \times 10^{-4}$ | 58.0 | Yes |
| 14 | GEP25 | 1.317 (1.135–1.530) | $2.94 \times 10^{-4}$ | $8.20 \times 10^{-4}$ | 64.5 | Yes |
| 15 | GEP27 | 0.782 (0.671–0.912) | 0.002 | 0.004 | 31.2 | Yes |
| 16 | GEP29 | 0.826 (0.720–0.947) | 0.006 | 0.015 | 0.0 | Yes |
| 17 | GEP37 | 1.180 (1.030–1.351) | 0.017 | 0.038 | 0.0 | Yes |
| 18 | GEP35 | 1.178 (1.019–1.361) | 0.027 | 0.058 | 0.0 | Yes |
| 19 | GEP38 | 1.145 (1.009–1.299) | 0.036 | 0.075 | 68.5 | No |
| 20 | GEP17 | 1.137 (0.989–1.308) | 0.072 | 0.140 | 0.0 | Yes |
| 21 | GEP20 | 1.102 (0.974–1.248) | 0.124 | 0.231 | 0.0 | No |
| 22 | GEP06 | 0.905 (0.783–1.045) | 0.174 | 0.309 | 45.5 | No |
| 23 | GEP11 | 0.905 (0.781–1.049) | 0.187 | 0.316 | 18.0 | No |
| 24 | GEP28 | 0.914 (0.795–1.051) | 0.207 | 0.337 | 20.1 | No |
| 25 | GEP10 | 0.921 (0.807–1.052) | 0.224 | 0.350 | 45.2 | No |
| 26 | GEP32 | 1.090 (0.938–1.266) | 0.261 | 0.392 | 0.0 | No |
| 27 | GEP14 | 0.933 (0.815–1.068) | 0.315 | 0.425 | 82.4 | No |
| 28 | GEP24 | 0.929 (0.806–1.070) | 0.306 | 0.425 | 72.9 | No |
| 29 | GEP30 | 0.935 (0.821–1.066) | 0.316 | 0.425 | 0.0 | No |
| 30 | GEP19 | 1.041 (0.909–1.192) | 0.558 | 0.704 | 0.0 | No |
| 31 | GEP34 | 0.962 (0.844–1.096) | 0.560 | 0.704 | 0.0 | No |
| 32 | GEP18 | 0.967 (0.849–1.102) | 0.618 | 0.754 | 4.6 | No |
| 33 | GEP13 | 0.968 (0.843–1.111) | 0.640 | 0.756 | 25.7 | No |
| 34 | GEP07 | 0.970 (0.834–1.129) | 0.697 | 0.799 | 83.8 | No |
| 35 | GEP02 | 0.985 (0.864–1.123) | 0.827 | 0.906 | 0.0 | No |
| 36 | GEP05 | 1.010 (0.886–1.152) | 0.880 | 0.906 | 43.2 | No |
| 37 | GEP23 | 1.012 (0.862–1.189) | 0.882 | 0.906 | 24.6 | No |
| 38 | GEP26 | 1.014 (0.883–1.164) | 0.844 | 0.906 | 0.0 | No |
| 39 | GEP03 | 0.994 (0.856–1.154) | 0.936 | 0.936 | 85.3 | No |

Hazard ratios (HRs), 95% confidence intervals (CIs) and $P$ values are from fixed-effect inverse variance meta-analysis of the four covariate-adjusted cohort Cox models. False discovery rates (FDRs) use Benjamini–Hochberg correction across all 39 signal GEPs. “Same direction” requires all four cohort HRs to be either above or below 1.

Supplementary Table 3: Representative enrichment terms for all signal GEPs in cross-species cerebellum.

| GEP | Top ten positive-weight genes | Representative enriched terms |
|---|---|---|
| **GEP01** | *KCNIP4*, *KCND2*, *RBFOX1*, *PRDX5*, *SYT1*, *CADPS2*, *NRG1*, *GRIK2*, *FGF14*, *LRP1B* | Synaptic signaling<br>Neuron projection development<br>Synaptic vesicle cycle |
| **GEP02** | *CCDC3*, *SELENOW*, *ANO6*, *PPIA*, *HSP90AB1*, *GPM6B*, *FTH1*, *DBI*, *SPARC*, *B2M* | Cytoplasmic translation<br>Peptide metabolic process<br>Translation at presynapse |
| GEP03 | *HSPA8*, *FAU*, *HSP90AB1*, *PCP4*, *TTR*, *PRDX5*, *DBI*, *CPE*, *RPL37*, *SPARCL1* | Proton motive force driven ATP synthesis<br>Aerobic respiration<br>Oxidative phosphorylation |
| **GEP04** | *CALB1*, *YWHAH*, *CALM2*, *NSG1*, *PEBP1*, *HOMER3*, *PVALB*, *HPCAL1*, *ITM2B*, *PPP1R17* | Synaptic signaling<br>Transport along microtubule<br>Neuron development |
| GEP05 | *RPL18*, *RPS8*, *RPS9*, *RPS7*, *RPL8*, *RPS19*, *FAU*, *STMN1*, *RPLP0*, *RPL7* | Cytoplasmic translation<br>Peptide biosynthetic process<br>Translation at presynapse |
| GEP06 | *PCP2*, *ITPR1*, *GNG13*, *CALB1*, *PPP1R17*, *INPP5A*, *IGSF5*, *SLC1A6*, *RGS8*, *FABP7* | Synaptic signaling<br>Anterograde axonal transport<br>Modulation of chemical synaptic transmission |
| **GEP07** | *PLP1*, *CRYAB*, *MOBP*, *CNP*, *PPP1R14A*, *QDPR*, *GFAP*, *MAG*, *FTH1*, *SPP1* | Myelination<br>Oligodendrocyte development<br>Glial cell development |
| GEP08 | *NDRG2*, *SNAP25*, *FAM107A*, *CALM2*, *PEBP1*, *RPL37*, *GALNT12*, *RPS8*, *PCP2*, *RPS24* | Cytoplasmic translation<br>Peptide biosynthetic process<br>Ribosomal small subunit biogenesis |
| **GEP09** | *SLC1A3*, *SPARCL1*, *CPE*, *FABP7*, *CD81*, *GSTM3*, *APOE*, *GPM6A*, *SLC4A4*, *LCAT* | Positive regulation of synaptic transmission<br>Neuron projection development<br>Gliogenesis |
| GEP10 | *PSAP*, *HPCAL1*, *SCN1B*, *ATP1A1*, *PRDX5*, *SNAP25*, *GAPDH*, *SLC24A3*, *HSP90AB1*, *CST3* | Cytoplasmic translation<br>Aerobic respiration<br>Ribosome biogenesis |
| GEP11 | *PEBP1*, *RPL30*, *SNAP25*, *CPE*, *PRDX5*, *SPARCL1*, *DBI*, *RPL37*, *FTH1*, *FSIP1* | Cytoplasmic translation<br>Peptide biosynthetic process<br>Translation at presynapse |
| GEP12 | *ATP1B1*, *CST3*, *CSMD1*, *RPL30*, *SLC24A3*, *NRXN3*, *HSPA8*, *KCNIP4*, *PVALB*, *AUTS2* | Synaptic signaling<br>Synapse assembly<br>Receptor localization to synapse |

Continued on next page

**Supplementary Table 3 (continued)**

| GEP | Top ten positive-weight genes | Representative enriched terms |
|---|---|---|
| GEP13 | *HSP90AB1*, *ARHGAP32*, *CALM2*, *EIF1*, *SLC24A3*, *RPL27A*, *RPL9*, *RPS10*, *TMSB10*, *FSIP1* | Cytoplasmic translation<br>Peptide biosynthetic process<br>Translation at presynapse |
| **GEP14** | *HSP90AB1*, *NEGR1*, *GRIK2*, *NRXN1*, *FGF14*, *SYNE1*, *NKAIN2*, *RBFOX1*, *CHN2*, *EIF1* | Synaptic membrane adhesion<br>Synapse organization<br>Synaptic signaling |
| **GEP15** | *RPL30*, *PVALB*, *EIF1*, *NDRG2*, *PCP4*, *SNAP25*, *FAU*, *SLC1A3*, *HOMER3*, *GAPDH* | Synaptic signaling<br>Long term synaptic potentiation<br>Receptor localization to synapse |
| GEP16 | *RPS21*, *CST3*, *RPS19*, *CLU*, *SPARCL1*, *FAU*, *CD81*, *CALM2*, *RPL30*, *RPL35* | Cytoplasmic translation<br>Peptide biosynthetic process<br>Translation at presynapse |
| GEP17 | *MBP*, *GFAP*, *CST3*, *FTH1*, *MOBP*, *RGCC*, *CALM2*, *RPS19*, *RPS7*, *RPS6* | Cytoplasmic translation<br>Peptide biosynthetic process<br>Translation at presynapse |
| GEP18 | *RPS15*, *RPS19*, *SLC1A3*, *NDRG2*, *PVALB*, *RPL30*, *FABP7*, *HSPA8*, *PLP1*, *FAM107A* | Proton motive force driven ATP synthesis<br>Oxidative phosphorylation<br>Nucleoside triphosphate metabolic process |
| GEP19 | *PCP4*, *SNAP25*, *RPL9*, *FTH1*, *RPS15*, *RPS23*, *CST3*, *RPS8*, *RPL27A*, *RPS10* | Cytoplasmic translation<br>Peptide biosynthetic process<br>Ribosome biogenesis |
| GEP20 | *S100B*, *CST3*, *FTH1*, *GALNT12*, *RPL37*, *GFAP*, *GALNTL6*, *DBI*, *CALM2*, *KIF5C* | Cytoplasmic translation<br>Peptide biosynthetic process<br>Translation at presynapse |
| GEP21 | *GRID2*, *SNAP25*, *CSMD1*, *UNC5C*, *AUTS2*, *FRMD4A*, *EIF1*, *NRXN3*, *KCNIP4*, *NLGN1* | Synapse assembly<br>Postsynapse organization<br>Synaptic signaling |
| GEP22 | *PTGDS*, *APOD*, *VIM*, *CLU*, *FTH1*, *IGFBP7*, *TPT1*, *RPLP2*, *TMSB10*, *MGP* | Cytoplasmic translation<br>Peptide biosynthetic process<br>Organonitrogen compound biosynthetic process |
| GEP23 | *TTR*, *FTH1*, *GNG13*, *CST3*, *DBI*, *NKAIN2*, *LDHB*, *PVALB*, *IGSF5*, *RPL37* | Proton motive force driven ATP synthesis<br>Nucleoside triphosphate metabolic process<br>Aerobic respiration |
| GEP24 | *APOE*, *FTH1*, *RPS19*, *RPL9*, *RPS23*, *CALM2*, *PCP2*, *RPL35*, *STMN2*, *SNAP25* | Cytoplasmic translation<br>Translation<br>Translation at presynapse |

All 24 signal GEPs are listed; background GEP00 is excluded. Bold identifiers denote GEPs discussed in the corresponding Results section. The ten genes with the largest positive $Q$ weights are shown in descending order using standardized gene symbols. Enrichment used the larger set of positive-weight genes in the top 2% of the complete profile ranking and Gene Ontology Biological Process. Three to five significant terms were selected to summarize enriched themes and reduce repetitive entries. All displayed terms have Benjamini–Hochberg FDR $< 0.05$, corrected across all signal GEPs and tested terms within each collection. Significant enrichment results and FDR values are provided in Supplementary Data 1.

Supplementary Table 4: Representative enrichment terms for all signal GEPs in mouse embryogenesis.

| GEP | Top ten positive-weight genes | Representative enriched terms |
|---|---|---|
| GEP01 | *TUBB3*, *HBA-X*, *CRABP2*, *MARCKSL1*, *H2AFV*, *RPL35*, *NES*, *SERF2*, *RPL34*, *DDAH2* | Cytoplasmic translation<br>Translation at presynapse<br>Translation at postsynapse |
| **GEP02** | *MAPT*, *RTN1*, *STMN2*, *TUBB2B*, *GAP43*, *INA*, *DPYSL3*, *MAP2*, *CRMP1*, *TUBB2A* | Neuron projection morphogenesis<br>Axonogenesis<br>Synapse organization |
| GEP03 | *HBB-Y*, *MDK*, *MIF*, *LRBA*, *NNAT*, *SNRPB*, *PPIA*, *H2AFZ*, *YBX1*, *HSPE1* | ATP synthesis coupled electron transport<br>Oxidative phosphorylation<br>Aerobic respiration |
| GEP04 | *AFP*, *ALB*, *MT1*, *GPX1*, *APOA2*, *HBB-BT*, *SLC25A37*, *HBB-BS*, *HBA-A2*, *TRF* | Erythrocyte development<br>Erythrocyte differentiation<br>Porphyrin containing compound metabolic process |
| GEP05 | *COL1A2*, *POSTN*, *COL3A1*, *SPARC*, *COL5A2*, *MALAT1*, *FSTL1*, *COL6A3*, *COL5A1*, *COL12A1* | Skeletal system development<br>Extracellular matrix organization<br>Collagen fibril organization |
| **GEP06** | *ACTA1*, *MYH3*, *MYLPF*, *ACTC1*, *MYL1*, *TNNT1*, *TPM2*, *TNNC2*, *TNNI1*, *MYL4* | Striated muscle contraction<br>Sarcomere organization<br>Muscle cell differentiation |
| GEP07 | *HBB-Y*, *GM19951*, *JARID2*, *CD44*, *ZC3H7A*, *CMSS1*, *GM26561*, *TMTC2*, *D130009I18RIK*, *GM15564* | RNA splicing<br>DNA metabolic process<br>Epigenetic regulation of gene expression |
| GEP08 | *HBB-BS*, *HBA-A1*, *HBA-A2*, *HBB-BT*, *TUBA1A*, *HBB-Y*, *MT-ND4*, *COL1A2*, *NNAT*, *COL3A1* | Cytoplasmic translation<br>Supramolecular fiber organization<br>Enzyme linked receptor protein signaling pathway |
| GEP09 | *HMGB2*, *HBB-BS*, *HMGB1*, *HNRNPA3*, *TOP2A*, *HMGN2*, *H2AFZ*, *MKI67*, *IGFBP2*, *SET* | RNA processing<br>RNA splicing<br>Mitotic cell cycle<br>Chromosome organization |
| GEP10 | *KRTDAP*, *DMKN*, *KRT10*, *S100A6*, *KRT1*, *COL3A1*, *HBB-BS*, *GM19951*, *KRT77*, *COL1A2* | Cytoplasmic translation<br>Peptide metabolic process<br>Ribosome biogenesis |
| GEP11 | *TTN*, *NEB*, *MEF2C*, *DMD*, *PALLD*, *MYBPC1*, *LDB3*, *DLG2*, *NEXN*, *MALAT1* | Muscle cell differentiation<br>Sarcomere organization<br>Muscle contraction |
| GEP12 | *ID4*, *LGALS1*, *CCND2*, *IGF2*, *HBB-BH1*, *TPM1*, *GPC3*, *CNN2*, *COL3A1*, *MT2* | Supramolecular fiber organization<br>Actin cytoskeleton organization<br>Cytoplasmic translation |
| GEP13 | *KCTD12*, *HBB-Y*, *MT-ND4*, *KRTDAP*, *GM42418*, *HMGB2*, *FOXP2*, *IGFBP5*, *EBF3*, *AFF3* | Growth<br>Heart development<br>Central nervous system development |
| GEP14 | *MT2*, *GM45774*, *MT1*, *ASXL3*, *HBA-X*, *RPL35*, *RPL34*, *MGST3*, *SMIM1*, *RPS6* | Cytoplasmic translation<br>Translation at presynapse<br>Erythrocyte development |
| GEP15 | *GM42418*, *CDK8*, *IL31RA*, *CAMK1D*, *GPHN*, *LARS2*, *CMSS1*, *GM26561*, *MT-ND1*, *JARID2* | Cytoplasmic translation<br>Oxidative phosphorylation<br>Aerobic respiration |

Continued on next page

**Supplementary Table 4 (continued)**

| GEP | Top ten positive-weight genes | Representative enriched terms |
|---|---|---|
| GEP16 | *LUM*, *COL3A1*, *LGALS1*, *HMGB2*, *COL5A2*, *FN1*, *CSRP2*, *TAGLN*, *COL1A2*, *SFRP2* | Collagen fibril organization<br>Extracellular matrix organization<br>Oxidative phosphorylation |
| GEP17 | *MT-ND6*, *PABPC1*, *MT-ND1*, *RPS21*, *MT-ND2*, *MT-CYTB*, *MT1*, *HBA-X*, *LDHA*, *RPL22L1* | Cytoplasmic translation<br>Ribosome biogenesis<br>Chromosome organization |
| GEP18 | *PTN*, *TUBB3*, *HBB-BS*, *HMGB2*, *ACTA1*, *ALDOC*, *NDUFA4*, *RPL22L1*, *TNC*, *PCP4* | Oxidative phosphorylation<br>Aerobic respiration<br>ATP synthesis coupled electron transport |
| GEP19 | *MYL2*, *MYL7*, *MYL3*, *MYH7*, *TNNT2*, *MYH6*, *ACTC1*, *TNNI3*, *MYL4*, *NPPA* | Aerobic respiration<br>Cardiac muscle tissue development<br>Cardiac muscle contraction |
| GEP20 | *FABP7*, *VIM*, *CKB*, *PEA15A*, *QK*, *HES5*, *NPAS3*, *PANTR1*, *PTPRZ1*, *NFIA* | Central nervous system development<br>Glial cell differentiation<br>Neural precursor cell proliferation |
| GEP21 | *ATXN7L3B*, *HBA-A1*, *HBB-BS*, *HBA-A2*, *RPL34*, *HBB-BT*, *MT2*, *RPS21*, *RPL35A*, *MT-ND6* | Cytoplasmic translation<br>Peptide biosynthetic process<br>Ribosome biogenesis |
| GEP22 | *TENM3*, *RBFOX1*, *NRXN1*, *HOXB8*, *EBF1*, *LINGO2*, *PCP4*, *PRRXL1*, *EBF3*, *GRM7* | Synapse organization<br>Neuron projection morphogenesis<br>Cell junction organization |
| GEP23 | *TSC22D1*, *MSX1*, *RPS12*, *ACTC1*, *MYLPF*, *UBE2C*, *KRT15*, *MYH3*, *ACTA1*, *HBB-BS* | Striated muscle cell development<br>Myofibril assembly<br>Intermediate filament based process |
| GEP24 | *GPC6*, *KRTDAP*, *HMGA2*, *HBB-Y*, *MT-ND4*, *EFNA5*, *ROBO1*, *PBX1*, *HMGB1*, *MDK* | Central nervous system development<br>Chromatin remodeling<br>Cell cycle |
| **GEP25** | *COL2A1*, *COL11A1*, *COL9A1*, *COL9A3*, *WWP2*, *SOX9*, *COL9A2*, *COL11A2*, *SOX6*, *CNMD* | Cartilage development<br>Chondrocyte differentiation<br>Extracellular matrix organization |
| GEP26 | *HBA-X*, *HBB-Y*, *PABPC1*, *HBA-A2*, *HBA-A1*, *MKRN1*, *EPB41*, *APEX2*, *ALAS2*, *FTH1* | Erythrocyte differentiation<br>Supramolecular fiber organization<br>Hydrogen peroxide catabolic process |
| **GEP27** | *GM26917*, *HBB-Y*, *MT-ND6*, *HBA-X*, *MT-ND1*, *YBX1*, *LRBA*, *HMGB2*, *PTMS*, *PABPC1* | Cytoplasmic translation<br>Peptide metabolic process<br>Translation at presynapse |
| GEP28 | *NEFL*, *NEFM*, *SNCG*, *PRPH*, *S100A10*, *GAP43*, *STMN2*, *UCHL1*, *FABP7*, *RBMS3* | Cell morphogenesis involved in neuron differentiation<br>Neuron projection morphogenesis<br>Axon development |
| GEP29 | *FABP5*, *HBB-Y*, *IGF2*, *HMGB2*, *TCF4*, *PAX1*, *ACTA2*, *HBA-X*, *H19*, *CAMK1D* | Enzyme linked receptor protein signaling pathway<br>Tissue morphogenesis<br>Heart development |
| GEP30 | *TTR*, *HBA-X*, *HBB-Y*, *CLIC6*, *SLC6A15*, *KCNE2*, *TUBB3*, *EZR*, *TRPM3*, *DEFB11* | Central nervous system development<br>One carbon compound transport<br>Regulation of cellular response to growth factor stimulus |

Continued on next page

**Supplementary Table 4 (continued)**

| GEP | Top ten positive-weight genes | Representative enriched terms |
|---|---|---|
| **GEP31** | *KRT15*, *KRT5*, *PERP*, *KRT14*, *DSP*, *KRT10*, *SFN*, *KRT1*, *S100A6*, *KRT4* | Keratinocyte differentiation<br>Epidermis development<br>Keratinization<br>Translation at presynapse |
| GEP32 | *CRABP1*, *HMGB2*, *HBB-BS*, *ACTA2*, *IGFBP2*, *COL1A1*, *FABP7*, *HMGB1*, *NFIB*, *HBB-BT* | Muscle tissue development<br>Central nervous system development<br>Cardiac muscle tissue morphogenesis |
| GEP33 | *COL1A1*, *COL1A2*, *SPARC*, *GM42418*, *SERPINH1*, *TUBA1A*, *HBB-Y*, *RCN3*, *NNAT*, *HMGB2* | Extracellular matrix organization<br>Collagen fibril organization<br>Bone development |
| GEP34 | *NFIB*, *CCND2*, *FOXG1*, *TCF4*, *NEUROD6*, *SOX11*, *NFIA*, *GM43126*, *CELF2*, *NEUROD2* | Axon development<br>Cell morphogenesis involved in neuron differentiation<br>Telencephalon development |
| GEP35 | *AUTS2*, *HBB-Y*, *MT1*, *COL1A1*, *SNCG*, *LGALS1*, *PRPH*, *LDHA*, *S100A10*, *LUM* | Chordate embryonic development<br>Microtubule cytoskeleton organization<br>Striated muscle contraction |
| GEP36 | *IGFBP2*, *ATP1A2*, *PTGDS*, *GJA1*, *RBP1*, *VTN*, *SELENOP*, *TRPM3*, *MT-CO1*, *CLDN11* | Tube morphogenesis<br>Regulation of cell motility<br>Blood vessel development<br>Extracellular matrix organization |
| GEP37 | *RPS12*, *HBB-Y*, *CAMK1D*, *RPS21*, *PPIA*, *RPS27A*, *TPT1*, *CMSS1*, *HBA-X*, *RPL34* | Cytoplasmic translation<br>Translation at presynapse<br>Ribosome biogenesis |
| GEP38 | *HBB-BH1*, *ASXL3*, *HMGB2*, *HBB-Y*, *HBA-X*, *MT1*, *NUDT4*, *HBA-A1*, *CRABP1*, *KRTDAP* | Cytoplasmic translation<br>Peptide biosynthetic process<br>Ribonucleoprotein complex biogenesis |
| GEP39 | *MSX1*, *PRRX1*, *WNT5A*, *CXCL14*, *TWIST1*, *TRPS1*, *PRRX2*, *ALX1*, *HPSE2*, *LGALS1* | Embryonic limb morphogenesis<br>Skeletal system development<br>Ossification |
| GEP40 | *IGFBPL1*, *SOX11*, *TAGLN3*, *EBF1*, *CD24A*, *HIST3H2BA*, *NEUROD6*, *BCL11B*, *DCC*, *H3F3B* | Regulation of neuron projection development<br>Axon development<br>Cell morphogenesis involved in neuron differentiation |
| GEP41 | *TCF7L2*, *LHX9*, *NR2F1*, *TENM2*, *TFAP2B*, *ZFHX3*, *SHOX2*, *CCND2*, *GAP43*, *NR2F2* | Forebrain development<br>Telencephalon development<br>Neuron projection morphogenesis |
| GEP42 | *EBF2*, *EBF3*, *EBF1*, *GAS2*, *AMOT*, *ACTA2*, *HMGB2*, *FN1*, *CRABP2*, *IGFBP5* | Skeletal system development<br>Anterior posterior pattern specification<br>Connective tissue development |
| **GEP43** | *ACTA2*, *MYL9*, *ACTG2*, *MYLK*, *TAGLN*, *MYH11*, *MYL6*, *FLNA*, *LSP1*, *CALD1* | Smooth muscle contraction<br>Actomyosin structure organization<br>Muscle structure development<br>Heart development |
| GEP44 | *DBI*, *MT3*, *ALDOC*, *SLC1A3*, *TUBB3*, *SPARCL1*, *HMGB2*, *TNC*, *ALDH1L1*, *IGFBPL1* | Central nervous system development<br>Regulation of neuron projection development<br>Gliogenesis |
| GEP45 | *PCDH9*, *HBB-Y*, *HMGB2*, *IGFBPL1*, *MALAT1*, *TENM2*, *ZIC1*, *PTPRD*, *EPHA3*, *LRRC4C* | cell–cell adhesion via plasma membrane adhesion molecules<br>Synapse organization<br>Synaptic membrane adhesion |

Continued on next page

**Supplementary Table 4 (continued)**

| GEP | Top ten positive-weight genes | Representative enriched terms |
|---|---|---|
| GEP46 | *PANTR1*, *NEFL*, *NEFM*, *TPM1*, *IGF2*, *GM26917*, *GPC3*, *RBMS1*, *PRPH*, *HMGA2* | Supramolecular fiber organization<br>Muscle structure development<br>Regulation of cell motility |
| GEP47 | *PEG3*, *NNAT*, *IGFBPL1*, *COPG2*, *MAP1B*, *IGFBP2*, *SNHG11*, *FABP7*, *KRTDAP*, *NDN* | Synapse organization<br>Cell morphogenesis involved in neuron differentiation<br>Axonogenesis |
| GEP48 | *TSHZ1*, *HOXB8*, *HOXA5*, *EBF3*, *HOXB9*, *HOXB3*, *RBFOX1*, *TSHZ2*, *ZIC1*, *EBF1* | Pattern specification process<br>Regulation of neuron projection development<br>Central nervous system development |
| GEP49 | *LSAMP*, *IGFBPL1*, *CSMD1*, *TUBB3*, *SNHG11*, *MEG3*, *RBFOX1*, *PCP4*, *FABP7*, *NRG3* | Synapse organization<br>cell–cell adhesion via plasma membrane adhesion molecules<br>Generation of precursor metabolites and energy |
| GEP50 | *PLAGL1*, *DLK1*, *HBB-Y*, *PEG10*, *GM42418*, *MSX1*, *MT-ND4*, *NRK*, *ITM2A*, *ASB4* | Chromatin remodeling<br>Negative regulation of DNA templated transcription<br>In utero embryonic development<br>Cell cycle |
| GEP51 | *APOE*, *SELENOP*, *CTSB*, *RBP4*, *AFP*, *APOA2*, *TRF*, *CTSD*, *HMGB2*, *PF4* | Blood coagulation<br>Fibrinolysis<br>Inflammatory response |
| GEP52 | *COL4A1*, *SPARCL1*, *COL18A1*, *CLDN5*, *ZEB2*, *NR2F2*, *HBB-BS*, *RPS12*, *HNRNPA3*, *CXCL12* | Tube morphogenesis<br>Blood vessel development<br>Regulation of cell migration |
| GEP53 | *DCN*, *COL3A1*, *DLK1*, *S100A6*, *COL2A1*, *ANXA2*, *KRT10*, *CMA1*, *VIM*, *CRIP1* | Regulation of epithelial cell proliferation<br>Collagen fibril organization<br>Cellular response to growth factor stimulus |
| GEP54 | *CDKN1C*, *HMGB2*, *MT-ND1*, *MT-ND6*, *MSX1*, *HBB-BS*, *ACTA2*, *MT-CYTB*, *NNAT*, *MT-ND4* | Cardiac ventricle development<br>Muscle tissue development<br>Cellular component morphogenesis |
| GEP55 | *PKM*, *GPHN*, *LARS2*, *CAMK1D*, *GM15564*, *IL31RA*, *CD44*, *GM26561*, *ALCAM*, *MALAT1* | Cytoplasmic translation<br>Peptide biosynthetic process<br>Cellular component morphogenesis |
| GEP56 | *MALAT1*, *HBB-BS*, *NNAT*, *H3F3B*, *SOX11*, *PRPH*, *H2AFV*, *TUBA1A*, *HBB-BT*, *RPL34* | RNA processing<br>Ribosome biogenesis<br>Cytoplasmic translation |
| GEP57 | *KIF26B*, *IGFBP2*, *HPSE2*, *COL1A1*, *WNT5A*, *ALX1*, *MAP1B*, *FABP7*, *RPL35A*, *POSTN* | Cytoplasmic translation<br>Ribosome biogenesis<br>Embryonic morphogenesis |
| GEP58 | *H19*, *FABP7*, *MSX1*, *PPIA*, *HMGB1*, *HMGN2*, *STMN1*, *H2AFZ*, *HMGB2*, *HNRNPA3* | Central nervous system development<br>Neuron projection morphogenesis<br>Axon development |
| GEP59 | *MAP1B*, *CCND2*, *GM43126*, *NES*, *ZIC1*, *HBB-Y*, *NCL*, *ANP32B*, *VIM*, *ZEB1* | Supramolecular fiber organization<br>Regulation of cell motility<br>Collagen fibril organization<br>Embryonic morphogenesis |

All 59 signal GEPs are listed; background GEP00 is excluded. Bold identifiers denote GEPs discussed in the corresponding Results section. The ten genes with the largest positive $Q$ weights are shown in descending order using standardized gene symbols. Enrichment used the larger set of positive-weight genes in the top 2% of the complete profile ranking and Gene Ontology Biological Process. Three to five significant terms were selected to summarize enriched themes and reduce repetitive entries. All displayed terms have Benjamini–Hochberg FDR $< 0.05$, corrected across all signal GEPs and tested terms within each collection. Significant enrichment results and FDR values are provided in Supplementary Data 1.

Supplementary Table 5: Literature-informed gene sets used in the mouse embryogenesis concordance analyses.

| GEP | Gene-set label | Fixed genes | Literature-supported interpretation | Representative literature |
|---|---|---|---|---|
| GEP25 | Chondrogenic | *Sox5*, *Sox6*, *Sox9*, *Acan*, *Col2a1*, *Col9a1*, *Col11a1* | Chondrogenic regulation and cartilage matrix formation. | Lefebvre et al. [5]; Chen et al. [6] |
| GEP31 | Epithelial-associated | *Krt5*, *Krt14*, *Krt15*, *Perp*, *Dsp*, *Sfn*, *Cxcl14*, *S100a6* | Basal epithelial identity, desmosomal adhesion and keratinocyte-associated expression. | Joost et al. [7]; Ihrie et al. [8]; Tsujihana et al. [9]; Dyring-Andersen et al. [10] |
| GEP02 | Neural-associated | *Mapt*, *Ina*, *Celf4*, *Map2*, *Nsg2*, *Cdk5r1*, *Dcx*, *Rtn1* | Neuronal differentiation and cytoskeletal organization. | Zeisel et al. [11]; Chen et al. [6] |
| GEP06 | Sarcomeric muscle | *Myh3*, *Tnnt3*, *Actn2*, *Des* | Embryonic and striated muscle differentiation and sarcomeric structure. | Agarwal et al. [12]; Cao et al. [13]; Buckingham and Rigby [14] |
| GEP43 | Smooth-contractile | *Acta2*, *Myl9*, *Actg2*, *Tagln* | Visceral smooth muscle and broader smooth-contractile programs. | Muhl et al. [15]; Owens et al. [16]; Li et al. [17] |

Gene sets were fixed before concordance analysis. The cited studies support the set-level functional annotations.

Supplementary Table 6: Dataset scale and input matrices.

| Analysis | Dataset | Sections / samples | Locations | Genes before selection | Genes used | SORT input |
|---|---|---|---|---|---|---|
| Simulation | Simulated spatial atlas | 20 replicates; 8 sections each | 10,368 / replicate | 8,000 | 5,000 HVGs | Scran-normalized log-expression |
| PDAC subtype | GSE282302 Visium | 20 subsets; 10 sections from 10 patients per subset | Subset-specific | 19,144 | SORT/ NMF 3,000; STAMP 6,000 | Normalized log-expression |
| PDAC full atlas | GSE282302 Visium | 108 sections; 39 patients | 352,102 fitted | 19,144 | 10,000 HVGs | Normalized log-expression |
| Cerebellum | Cross-species Stereo-seq | 9 sections; 3 species; 1 animal per species | 1,546,637 | 12,599 common | 4,000 HVGs | Normalized log-expression |
| Embryogenesis | MOSTA Stereo-seq | 8 sections; one per stage, E9.5–E16.5 | 520,389 | 8,000 retained | 8,000 | Normalized log-expression |

HVGs, highly variable genes.

Supplementary Table 7: SORT analysis-specific settings.

| Analysis | $K$ | HVGs | Graph $k$ | $\lambda_{\text{TV}}$ | $\lambda_{1,W}$ | $\lambda_{1,Q}$ | Epochs (stage 1/2) |
|---|---|---|---|---|---|---|---|
| Simulation main | 24 | 5,000 | 8 | 1.0 | 0.5 | 300 | 100/100 |
| PDAC subtype | 20 | 3,000 | 24 | 0.3 | 0.3 | 300 | 50/100 |
| PDAC full atlas | 40 | 10,000 | 24 | 0.3 | 0.3 | 300 | 50/100 |
| Cerebellum | 25 | 4,000 | 8 | 0.3 | 0.3 | 300 | 50/100 |
| Embryogenesis | 60 | 8,000 | 8 | 0.4 | 0.4 | 800 | 50/100 |

Unless otherwise stated, SORT used normalized log-expression, section-wise graphs without cross-section edges and component 0 as background. Graph TV was applied during stage 2 with $\varepsilon_{\text{TV}} = 10^{-2}$, with weights updated every five epochs. The manuscript parameter $\lambda_{\text{TV}}$ is called $\alpha$ in the implementation. The cerebellum analysis used adaptive sparsity weights for signal profiles as described in Supplementary Note 1.

Supplementary Table 8: Baseline method settings.

| Benchmark | Method | Input | $K$ | HVGs | Graph | Covariate |
|---|---|---|---|---|---|---|
| Simulation | PCA | Log-expression | 24 | 5,000 | None | None |
| Simulation | NMF | Nonnegative log-expression | 24 | 5,000 | None | None |
| Simulation | GraphPCA | Log-expression | 24 | 5,000 | kNN; Visium mode | None |
| Simulation | STAMP | Counts | 24 | 5,000 | kNN = 15 | Library ID |
| Simulation | NNMF | Nonnegative expression | 24 | 5,000 | Spatial coords. | Section batch |
| PDAC subtype | NMF | Log-expression | 20 | 3,000 | None | None |
| PDAC subtype | STAMP-noCov | Counts | 20 | 6,000 | kNN = 8 | None |
| PDAC subtype | STAMP | Counts | 20 | 6,000 | kNN = 8 | Patient ID |

kNN, k-nearest-neighbor graph.

Supplementary Table 9: Simulation sensitivity and ablation settings.

| Setting | $\lambda_{\text{TV}}$ | $\lambda_{1,W}$ | Simulation datasets |
|---|---|---|---|
| Main simulation | 1.0 | 0.5 | 20 |
| Conservative grid point | 0.3 | 0.3 | 10 |
| Nonzero grid | 0.3, 0.6, 1.0 | 0.3, 0.5, 0.7 | 10 |
| No spatial | 0 | 0.5 | 10 |
| No $W$-sparsity | 1.0 | 0 | 10 |
| Double ablation | 0 | 0 | 10 |
| Unit norm, nonorthogonal | 1.0 | 0.5 | 10 |

The nonzero grid contains all nine combinations of the listed $\lambda_{\text{TV}}$ and $\lambda_{1,W}$ values. All settings used $K = 24$, 5,000 HVGs, graph $k = 8$, $\lambda_{1,Q} = 300$ and stage 1/2 epochs of 100/100. Supplementary Note 2 defines the orthogonality design ablation.

Supplementary Table 10: Metrics and statistical tests.

| Analysis | Metric | Inputs | Unit | Summary / test | Interpretation |
|---|---|---|---|---|---|
| Simulation | Recovery PCC | $W_{\text{true}}, \widehat{W}$ | Program × replicate | Mean over 13 programs; paired Wilcoxon | Higher |
| Simulation | Boundary AUPRC | True and inferred edge gradients | Program × replicate | Mean over programs; paired summary | Higher |
| Simulation | Blur ratio | Boundary-adjacent gradients | Program × replicate | Mean over programs; paired summary | Lower |
| Simulation | Moran's $I$ | Selected component map | Program × replicate | Descriptive with boundary metrics | Descriptive |
| Simulation orthogonality | Profile cosine, duplicate matches and reconstruction error | $Q$, $X$ and $WQ^\top$ | Paired seed | Paired medians; exploratory exact Wilcoxon tests | Varies by metric |
| PDAC subtype | Reference-concordance PCC; target specificity | Target and alternative UCell scores with matched component activity | Subtype × iteration | Paired Wilcoxon for PCC; specificity descriptive | Higher |
| PDAC colocalization | Colocalization score | Global 90th-percentile selection; section-specific 80th-percentile scoring | Pair × section | Descriptive ranking; permutation assessment | Higher |
| PDAC survival | Cox HR | Bulk GEP score and OS | GEP × cohort | Covariate-adjusted Cox; fixed-effect meta-analysis; all-39 BH FDR | Effect sign |
| Cerebellum | Profile PCC | Normalized distance profiles | GEP | Mean pairwise PCC | Higher |
| Cerebellum | Coefficient of variation (CV) of mean activity | Mean unstandardized smoothed section-profile activity over $[-1, 1]$ | GEP | Descriptive across species means | Descriptive |
| Cerebellum | GEP02 and GEP14 detection effects | Domain-specific detection fractions | Gene × animal set | Matched-control calibration and orthology sensitivity | Effect sign |
| Cerebellum | Reciprocal GEP00–GEP01 contrast | Section-level detection fractions | Gene × section | Exact label permutation and BH correction across 24 genes | Effect sign |
| Embryogenesis | Temporal activity | Stage-wise mean GEP activity | GEP × stage | Descriptive groups; within-group correlation ordering | Descriptive |
| Embryogenesis | Gene-set concordance | Continuous $W$ and fixed gene-set score | Stage × annotation | Spearman and coverage-adjusted rank correlation; descriptive sensitivity | Descriptive |

PCC, Pearson correlation coefficient; AUPRC, area under the precision–recall curve; HR, hazard ratio; OS, overall survival; BH, Benjamini–Hochberg.

[5] Véronique Lefebvre, Richard R. Behringer, and Benoit de Crombrugghe. L-Sox5, Sox6 and Sox9 control essential steps of the chondrocyte differentiation pathway. *Osteoarthritis and Cartilage*, 9(Suppl A):S69–S75, 2001.
